# L-shape distribution of the relative substitution rate (c/μ) observed for SARS-COV-2’s genome, inconsistent with the selectionist theory, the neutral theory and the nearly neutral theory but a near-neutral balanced selection theory: implication on “neutralist-selectionist” debate

**DOI:** 10.1101/2023.01.01.522435

**Authors:** Chun Wu, Nicholas J. Paradis, Phillip M. Lakernick, Mariya Hryb

## Abstract

The genomic substitution rate (GSR) of SARS-CoV-2 exhibits a molecular clock feature and does not change under fluctuating environmental factors such as the infected human population (10^0^-10^7^), vaccination etc.. The molecular clock feature is believed to be inconsistent with the selectionist theory (ST). The GSR shows lack of dependence on the effective population size, suggesting Ohta’s nearly neutral theory (ONNT) is not applicable to this virus. Big variation of the substitution rate within its genome is also inconsistent with Kimura’s neutral theory (KNT). Thus, all three existing evolution theories fail to explain the evolutionary nature of this virus. In this paper, we proposed a Segment Substitution Rate Model (SSRM) under non-neutral selections and pointed out that a balanced mechanism between negative and positive selection of some segments that could also lead to the molecular clock feature. We named this hybrid mechanism as near-neutral balanced selection theory (NNBST) and examined if it was followed by SARS-CoV-2 using the three independent sets of SARS-CoV-2 genomes selected by the Nextstrain team. Intriguingly, the relative substitution rate of this virus exhibited an L-shaped probability distribution consisting with NNBST rather than Poisson distribution predicted by KNT or an asymmetric distribution predicted by ONNT in which nearly neutral sites are believed to be slightly deleterious only, or the distribution that is lack of nearly neutral sites predicted by ST. The time-dependence of the substitution rates for some segments and their correlation with the vaccination were observed, supporting NNBST. Our relative substitution rate method provides a tool to resolve the long standing “neutralist-selectionist” controversy. Implications of NNBST in resolving Lewontin’s Paradox is also discussed.

## Introduction

The COVID-19 pandemic was caused by severe acute respiratory coronavirus 2 (SARS-CoV-2) in December 2019 in Wuhan, China, with the human population undergoing six infection waves, ranging from 10^0^ to 10^7^ infected cases (**Figure 1A**). As of October 2022, SARS-CoV-2 resulted in 626 million infections and 6.56 million deaths worldwide, largely surpassing the infections and deaths caused by SARS-CoV and Middle East respiratory syndrome (MERS-CoV) (**Table 1**). Despite the rapid increase in global vaccinations since December 2020 (**Figure 1B**), multiple SARS-CoV-2 variants have emerged (alpha-omicron), of which the omicron variants have led to higher infection rates but lower virulence than previous variants, which may increase recurring infections. Continual mutations also raise concern about possible antibody-dependent enhancements (ADE) [1], which might increase the severity of COVID-19 infections. Furthermore, only a few FDA-approved therapeutics (e.g., Remdesivir, Paxlovid, and Molnupiravir) are available to treat COVID-19 infection and many promising drug candidates are still under development [2, 3].

**Figure 1.**
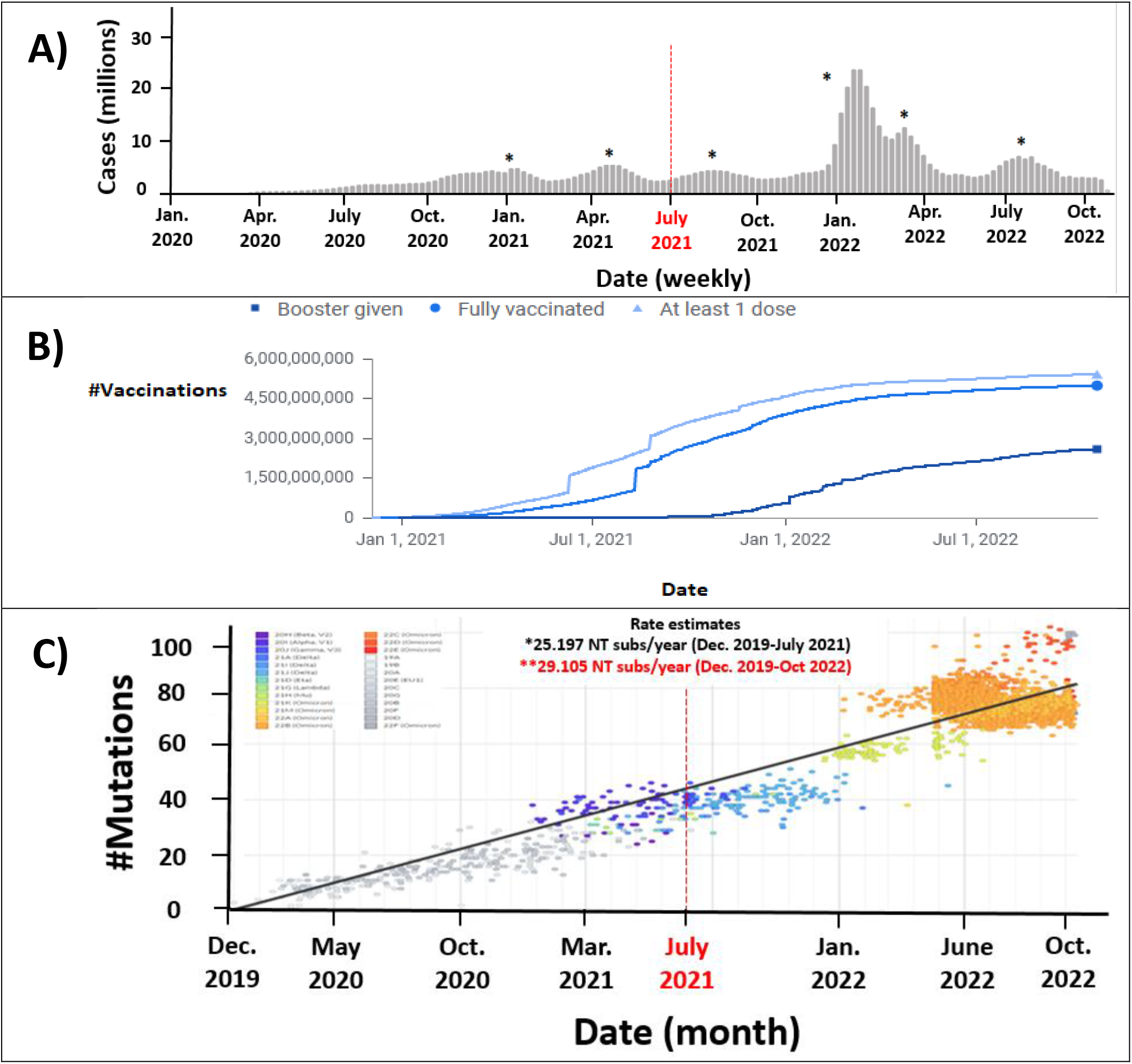
Timelines. **A**: Six waves of infections (https://covid19.who.int/) ranged from 10° to ~10^7^ human infections. **B**: Number of vaccinations starting from December 1 2021 (https://ourworldindata.org/covid-vaccinations?country=OWID_WRL). **C**: Constant NT substitution rate (# NT site per year per genome). Please refer to the Nextstrain web site for the present figures: https://nextstrain.org/ncov/gisaid/global.

**Table 1.** (**Top**) Number of global infections, global deaths, percent mortality rate and reported date of outbreak for SARS-CoV-2, SARS-CoV and MERS-CoV. (**Bottom**) Three independent datasets (11,198 genomes), collected from December 2019 to July 2021, to investigate the evolution of SARS-CoV-2.

| **Virus** | **# Global Infections** | **# Global Deaths** | **Mortality Rate** | **Date of Outbreak** |
| --- | --- | --- | --- | --- |
| SARS-CoV | 9,422 | 919 | 11% | 3/16/2003 |
| MERS-CoV | 2,494 | 858 | 34% | 9/23/2012 |
| SARS-CoV-2* | ~645 Million | ~6.63 Million | 1.0% | 12/31/2019 |
| **Genome** | **Dataset (# Genomes)** | **Virus Host** | **Genomic Variation<sup>1</sup>** | **Divergent Time** |
| SARS-CoV-2 | A1a (3856) | Human |  |  |
|  | A1b (3416) | Patients | <0.12% | <2 years |
|  | A1c (3926) |  |  |  |

\*Updated as of 12/14/2022. (World Health Organization. https://covid19.who.int/)
<sup>1</sup>The reference genome is the first human SARS-CoV-2 (Wuhan-Hu-1).

Understanding the molecular evolution nature of the SARS-CoV-2 genome is thus pressing to identify critical sites or regions in developing a rational strategy to mitigate COVID-19 in the long term. So far, three competing evolution theories have been developed to explain the evolution nature of a species. The first one is selectionist theory (ST) [4], in which the fittest mutations will survive through positive selection. The second one is Kimura’s neutral theory (KNT) [5, 6], in which the luckiest mutations will survive through genetic drifting.[7] The third one is Ohta’s nearly neutral theory (ONNT) [8–16], in which the fate of nearly neutral mutations, mostly being slightly deleterious, depends on the effective population size. Nonetheless, all three basic theories agree that deleterious mutations will be purified through evolution. Given the three basic theories, which one of them is true for this virus? Are the observed substitutions neutral, nearly neutral or advantageous? How can this be determined?

SARS-CoV-2 is a ~30 kb, single-stranded, positive-sense membrane-bound RNA virus, which shares high sequence similarity to other beta-coronaviruses such as SARS-CoV [17, 18] and MERS-CoV [19] [20]. The genomes of these beta-coronaviruses contain a Translated Region (TR) and Un-Translated Region (UTR). TR encodes a set of proteins critical to the virus life cycle, including open reading frame 1ab (Orf1ab) comprising 15 non-structural polyproteins (Nsp), major structural proteins (E, N, M and S) and accessory proteins (Orf3a, Orf6, Orf7a, Orf8 and Orf10) (**Tables S1a-S1b**). UTR does not encode protein but regulates viral replication and transcription [21]. 5’-untranslated regions (5’-UTR) occur at the beginning of each gene and a 3’-untranslated region (3’-UTR) occurs at the end of hypothetical Orf10 (**Table S1c**). 5’-UTRs contain Transcriptional Regulatory Sequences (TRS) that comprise 6-7 conserved core nucleotides (NTs) with variable positions in each gene (**Table S1d**). TRSs are involved in subgenomic RNA (sgRNA) biogenesis; during negative-sense strand synthesis, RdRp pauses RNA transcription when it reaches a TRS in the body (TRS-B) and switches the positive-sense RNA template strand to the TRS in the leader (TRS-L) in Orf1ab 5’-UTR, resulting in discontinuous transcription of sgRNA [22] [23].

In this paper, the genomic evolution of SARS-CoV-2 is assumed to be a repeating process of two basic steps (**Figure 2**). First, random mutations occur at any NT site of the genome with a putative constant mutation rate (*μ*) due to the polymerase error. Second, fixation of the mutations in the viral population (i.e., the census viral population subjected to genomic sequencing) leads to a substitution rate (*c*) through genetic drifting, environmental selection and other factors. A new substitution is classified as under positive selection if *c>μ*, as under neutral if *c=μ*, and as under negative selection if *c<μ*. The assumption that all NT sites in a genomic sequence solely follow neutral selection leads to a constant genomic substitution rate (*c=μ*), and this is defined as the Genomic Substitution Rate Model (GSRM). GSRM, as the strongest form of KNT (*f_0_*=1, where *f_0_* is the fraction of NT sites under neutral selection), serves as a null hypothesis that is rejectable if a NT site follows non-neutral selection (purifying/- or adaptive/+). It is clear that substitution rates vary within viral [24, 25] [26] [27] and eukaryotic genomes [28–32] and in different lineages of species (e.g., substitution rates much slower in humans compared to rabbits or Dengue-4 virus). [33, 34] Thus, strong variations of the substitution rate within the genome of SARS-CoV-2 should be expected. If so, GSRM and KNT would be untrue for SARS-CoV-2. It is therefore critical to investigate the substitution rate variations in the SARS-CoV-2 genome for identifying key NT sites [35, 36] and key genomic regions under positive selections. The key mutations and regions associated with high infection and virulence can be potential drug targets in treating COVID-19 [37–39].

**Figure 2.**
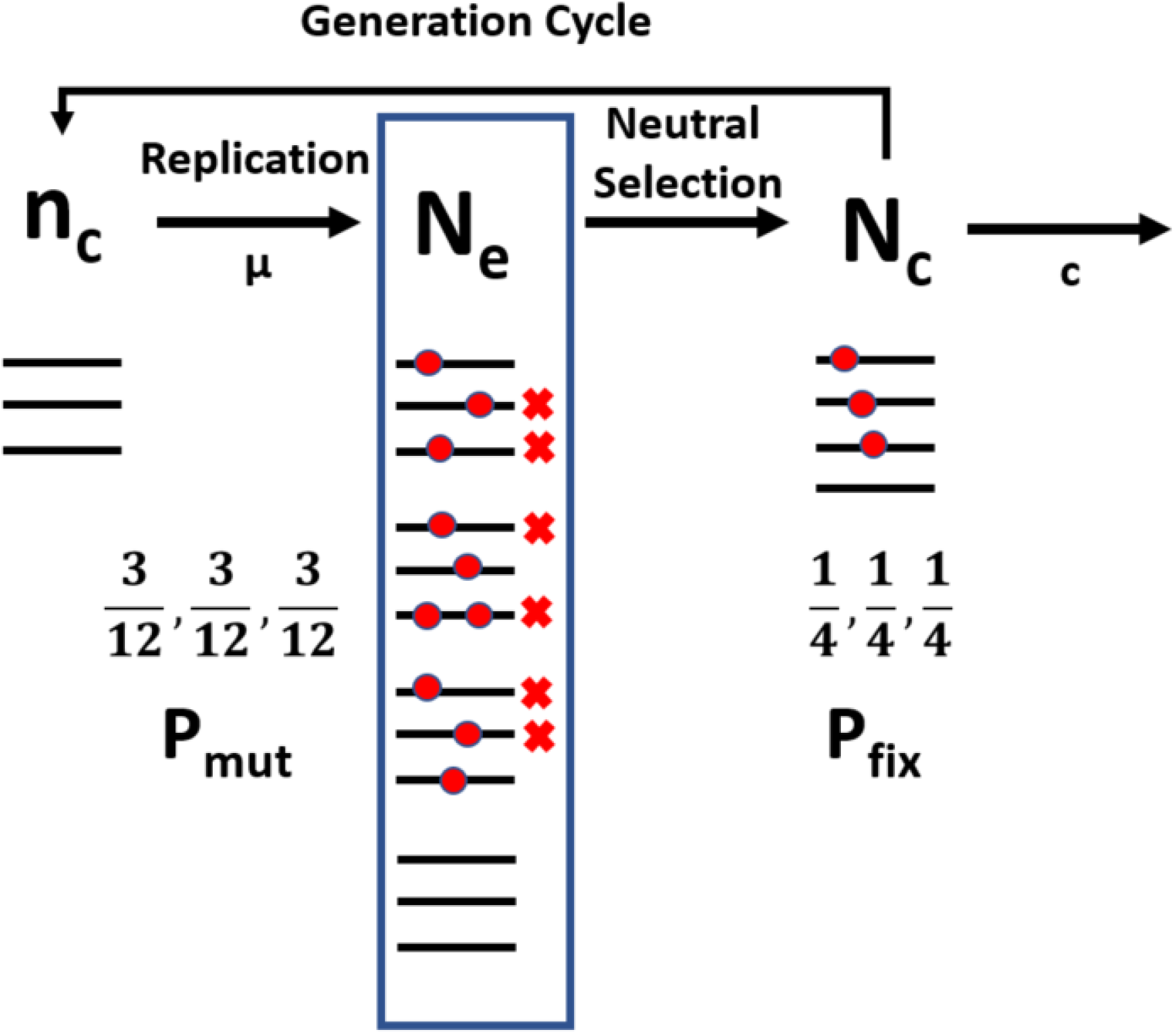
Simplified diagram of the Genomic Substitution Rate Model (GSRM) for a virus population over time under neutral selection only. A viral population (n_c_) undergoes replication and can undergo mutation at rate (μ) with probability (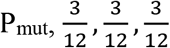) of three different mutants,12 12 12generating a produced population (N_e_) prior to the neutral selection. In the blue box, the mutant population (N_e_) undergoes neutral selection from the host (i.e., host immune response, host dies, vaccine pressure, etc.) and only some offspring survive, generating the observed viral population with fixed mutations (N_c_) with the probability (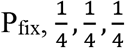) for the three different mutants. This process continues for succeeding generations within the host and after infecting other hosts. From N_c_, the substitution rate (c) can be observed. Black lines represent viral genomes. Red circles represent substitution mutations in the viral genomes.

Intriguingly, a near-constant, time-independent NT genomic substitution rate (GSR) in SARS-CoV-2 genomes, also known as a molecular clock feature (**Figure 1C**, 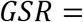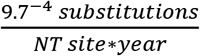) [40, 41] is evident despite dynamic changes in the infected human population **Figure 1A**) and significant increases in global vaccinations (**Figure 1B**). These observations appear to suggest that the molecular evolution of SARS-CoV-2 follow the GSRM (*c=μ)* or KNT [5, 6] (*c=μ*f_0_*) rather than ONNT[8–16] and ST.[4] This is because while GSRM and KNT predict the substitution rate is limited by the mutation rate (*μ*), which is a time-independent constant regardless of fluctuating environmental factors, the substitution rate (c) predicted by ONNT and ST should be strongly time-dependent. The time-dependent rate for ONNT comes from its strong dependence on the size of the effective population (*N_e_*) of this virus in fixing slightly deleterious mutations in the effective population. While the generation time of this virus (~5 days) does not change much, *N_e_* must have strong time dependence and a change of several orders of magnitude; because the virus census population (*N_c_*) is proportional to the host population size and the infected human population as the host population underwent six waves, with a range of 10^0^ to 10^7^ recorded cases so far (**Figure 1A**). In a stronger perspective to the ONNT, ST would also predict a strong time dependent substitution rate as the adaption response to the fluctuating environmental factors.

While the temporal features of this virus’s GSR appear to be inconsistent with ONNT and ST, the big variation of the substitution rate in the sequence space is also likely inconsistent with GSRM and KNT. What then can explain the molecular evolution of SARS-CoV-2? While the theoretical proofs for GSRM, KNT and ONNT to be consistent with the molecular clock has been provided, the proof that ST must be inconsistent with the molecular clock feature does not exist. Is it possible that some NT sites/segments are under (weakly) positive selection and other sites are under (weakly) negative selection, and that both selection types might exhibit time-dependent substitution rates, but are “balanced out” to generate the overall time-independent GSR? In other words, under which conditions can ST also lead to a molecular clock feature? There is also no reason to exclude some NT sites under the nearly neutral selection for ST. If so, a hybrid between ST and the neutral theories would be able to explain the evolution of this virus and many other species. In this paper, we proposed a Segment Substitution Rate Model (SSRM) under non-neutral selections and pointed out a balanced mechanism between negative selection and positive to obtain the molecular clock feature. We named this mechanism as a near-neutral balanced selection theory (NNBST) and examined if it was followed by SARS-CoV-2.

Although the GSR of SARS-CoV-2 virus has been reported (**Table S2**), the segment substitution rate (*c*, SSR) for the TR coding genes and non-coding UTR and TRS segments are not yet available [12, 42–47]. Although the selection type for the TR and each coding gene has been predicted via conventional Ka/Ks analysis (Ka: non-synonymous substitution rates; Ks: synonymous substitution rates), a convergent selection type for some genes (i.e., S-, E- and N-genes) is under debate (**Table S3**) [10–16]. Because the Ka/Ks method is not applicable to non-coding regions, UTR and TRS selection types have not been assigned. Although most evolutionary biologists consider most non-coding eukaryotic DNA evolves neutrally [48], some UTRs and TRSs of this virus are functionally important and likely under non-neutral selection. Indeed, several studies report that NT substitutions within the 5’UTR and 3’UTR of SARS-CoV-2 [49–56] impacted viral replication and transcription processes [57, 58], thus deciphering the variation of their substitution rates is also critical for determining their selection type.

In this study, we formulated the relative substitution rate (*c/μ*) test to quantify the selection type of an arbitrary genomic fragment or a NT site. A theoretical proof on the consistency between *c/μ* and the conventional Ka/Ks in classifying selection type for a gene in TR under near neutral selection for the genome was provided in the methods section. Systematic analysis using *c/μ* was done on three independent sets of SARS-CoV-2 genomes (a total of 11,198 sequences) from human patients within the first 19 months of the COVID-19 pandemic, which were selected and compiled by the Nextstrain team (**Table 1**) [40] [59]. In the first part of the results section, we investigated the substitution rate variation in the NT sequence space to see if the GSRM or KNT was not followed, respectively, by all NT sites or all substituted sites of SARS-CoV-2 using our *c/μ* test, including each coding gene in TR and UTR and TRS non-coding segments. The empirical probability distribution of *c/μ* was examined to check against the Poisson distribution as predicted by GSRM or KNT. The selection type of the 10 genes, 11 UTRs and 9 TRSs were determined using the *c/μ* test. The empirical check on the consistency of the selection type by *c/μ* with by Ka/Ks for the 10 coding genes was conducted. At NT level, the top 54 NT sites in UTR and 247 codon sites in TR that followed positive selection were identified. In the second part of the results section, the time-dependency of the substitution rates (*c* for the coding genes, UTRs and TRS; Ka/Ks for coding genes only) were examined using our SSRM to probe their time-dependent features for non-neutral selections. Additionally, the impact of the vaccine on viral genomic evolution was investigated using time correlation plots. In the discussion, our *c/μ* test in solving the long-standing “neutralist-selectionist” controversy on how to determine the selection type of a gene or a NT site in a genome exhibiting a high substitution rate (i.e., neutral VS positive/adaptive selection type) will be discussed. The implications of NNBST in the possible resolution of Lewontin’s Paradox, how sequence variability among species is far less than from what is expected based on differences in their population sizes will be also discussed, given the observation that the sequence variation was independent from the census population for this virus.

## Methods

### Genomic Substitution Rate Model (GSRM), a null hypothesis

A virus genome of length *N* consists of *i* functional segments (i.e., genes in TR, UTRs and TRSs) and segment *i* consists of *n_i_* NTs to give the following for non-overlapping segments:

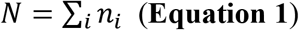

For SARS-CoV-2, *N*=29,903 NTs and *i*=10+11+9 (genes+UTRs+TRSs), if Orf1ab instead of Orf1a is used in this study. At time *t=0*, the number of NT mutations in segment ***i*** is zero for all segments to give:

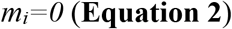

SARS-CoV-2 starts to replicate its genome using its replication machine (RNA-dependent RNA polymerase (RdRp) with a 3’ proofreading exonuclease (ExoN)) to maintain replication fidelity. Two assumptions are then made. **1**). The replication error rate at any NT site of the genome (the mutation rate) is the same 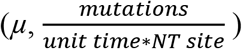, because the replication machine cannot sense the position of NTs in producing the nascent genome under random environmental perturbations. This spontaneous mutation process can be modeled as a Poisson process at a constant mutation rate of *μ*. **2**). The generated mutations in the produced virus population after virus replication (*N*_*e*_) is undergoing neutral selection only (i.e., no selection preference for any mutation type), such that the substitution rate (*c*, GSR) observed in the virus population generated after having undergone selection (i.e., the census population *N*_*c*_ subject to genomic sequencing) is the same as the mutation rate *μ* in the produced virus population (*N*_*e*_) in **Figure 1**:

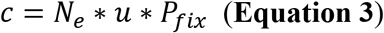

where μ *is* 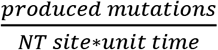 and *N_e_* * μ *is* 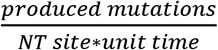 in the produced virus population prior to selection (*N_e_*). It’s important to note that the observed viral population *N*_*c*_ does not include viruses that have been eliminated through negative selection. *P*_*fix*_ is the fixation probability for a mutation or the observed probability of a mutation in the virus population after selection (*N*_*c*_), whereas *P*_*mut*_ is the probability for a mutation in the virus population prior to selection (*N*_*e*_). The fixation probability in the virus population after selection (*N*_*c*_) is equal to the probability of the mutation in the produced population (*N*_*e*_) prior to selection (i.e., *P*_*fix*_ = *P*_*mut*_) due to the neutral selection condition, and *P*_*mut*_ is proportional to the abundance of the mutation in the produced whole population (*N*_*e*_) prior to the selection (i.e., 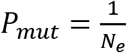 for a mutation) under the neutral selection condition, because any virus in *N*_*e*_ has an equal chance to be selected under the neutral selection only, thus simplifying **Equation 3**:

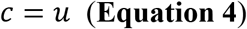

In the case where no selection exists and all viral offspring (non-mutants and mutants) survive, then N_c_ = N_e_. To observe this, the virus would need to reproduce in a reservoir free of selection pressure (i.e., no immune system) containing a constant supply of chemical and biological materials (i.e., NTs, host cells, etc.). The migration rate of the virus should also be sufficiently high to ensure mutation fixation within the global viral population. Humans, which are a migratory species, can enable the quicker virus spreading events between different countries and continents across the world; from early civilization to rapid globalization in the mid-20^th^ century, human-to-human virus spreading has increased at an unprecedented rate, given that the viruses exhibit low host fatality rates, high infectivity and slow incubation periods.

This time-independent constant substitution rate (*c*, GSR) under neutral selection only is equal to the mutation rate (*μ*) that is defined as the Fundamental Genomic Mutation Rate (FGMR), following the Jukes-Cantor substitution model [60]. The first assumption also implies that the FGMR of the genomic replication machinery (RdRp with 3’ ExoN) is independent from the virus genome sequence change, thus the replication error rate does not change significantly over a short period of time. When *c=μ* for every site within the genome, this is the basis of the GSRM, which is the strongest form of the KNT [4]. Consequently, the observed substitution rate for a genomic segment 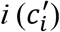 in the population is also a time-independent constant:

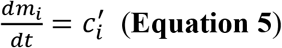

and

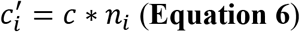

Where 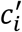 is the number of substitutions per unit time per segment *i*. Summing over evolution time *t*, the cumulative number of substitutions for segment ***i*** (*m_i_*) is equal to the product of the substitution rate per segment per time:

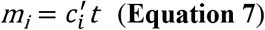

The integral substitution probability for a NT site within segment ***i*** (*P*_*i*_, substitution probability per NT site) at time *t* (the sequence variation from the initial NT site) is obtained by normalizing the number of substitutions (*m_i_*) by the total number of NTs of the segment ***i***:

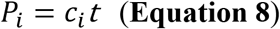

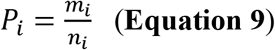

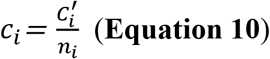

where *c*_*i*_ is the Segment Substitution Rate (SSR) in a unit of substitution per NT site per unit time. From the two assumptions, SSR of every segment is equal to FGMR:

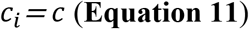

The time-independent substitution rate, which is independent of the virus population size and environmental conditions (e.g., vaccines), leads to a molecular clock feature (**Equation 8**) that can be used to estimate the divergent time between two or more species originating from a common ancestor using their sequence variances or predict sequence variance in the future [4, 7]. For example, given GSR of SARS-CoV-2 is 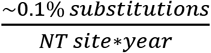, it will take ~1,031 years for the virus sequence to be 100% different from the first virus sequence.

While there are slight differences between our GSRM and KNT in terms of evolution, their fundamental nature is the same. The KNT assumes that in a genomic sequence, most sites undergo negative selection and are conserved; some sites undergo neutral selection; very rarely, sites undergo positive selection [4]. Because the mutations under negative selection will be purified in the population, these conserved sites will not contribute to the substitution rate of the virus. And because the positive selection sites are rare, these sites will not make a significant contribution to the overall substitution rate. Therefore, the substitution rate of all genomic sites under the KNT are dominated by the neutral sites and is just a fraction of FGMR (*c=μ*f_0_*, where *f_0_* is the fraction of neutral sites). In our GSRM, all sites within a genomic sequence are assumed to undergo neutral selection only (i.e., *f_0_* =1) and exhibit a constant substitution rate which is equal to the FGMR (*μ*). The single time-independent substitution rate (*c*) for all NT sites from GSRM or for some NT sites from KNT implies a Poisson process with that rate due to genetic drift [61, 62].

### Segment Substitution Rate Model (SSRM) and a balanced selection mechanism

SSRM is introduced to take the time-dependent non-neutral selection into consideration by allowing a specific substitution rate for a specific segment or a NT site:

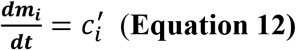

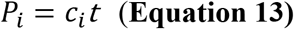

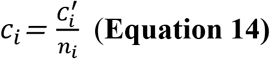

For example, while the substitution rate for the S-gene is 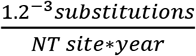, the substitution rate for the E-gene is 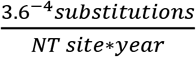. The integral substitution rate for each NT site within this viral genome (*P*) at time *t* (i.e., the genomic variation at time *t*) is product of time (*t*) with a GSR (*c*) in **Equation 15**. GSR (*c*) is a weighted sum of the substitution rate of each segment (*c*_*i*_) in **Equation 16**, and the weight is determined by the length of the segment over the length of the genome (*N*) in **Equation 18a**:

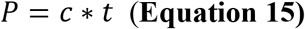

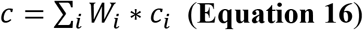

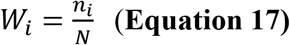

Please note that because the selection type on the virus could change over time, SSR 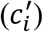 and GSR (*c*) could change over time due to time-dependent strong non-neutral selection types. Nonetheless, if the molecular evolution is under NNBST, the higher SSR of segment/NT sites under positive selection is balanced out by the lower SSR of the segments/NT sites, leading to a time-independent GSR that is close to FGMR (*μ*). Because the effect of ***N_e_*** on the SSR of weakly negative segments is also balanced out by that of the SSR of weakly positive segments, the overall substitution rate is independent from the ***N_e_***. This has been observed for SARS-CoV-2 in this study.

Because the virus evolution time is the same for each gene/UTR/TRS and the whole genome (e.g., 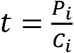), the variation ratio (e.g., 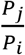) is proportional to the substitution rate ratio (e.g., 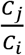) between between genes *j* and *i* or between gene *i* and the genome at a given time *t*:

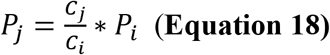

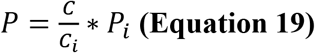

These substitution rate ratios can be also interpreted as relative substitution rates with the reference gene or the whole genome. The difference in the relative substitution rates reflect the difference in the net selection forces. **Equations 18-19** provide an alternative way to estimate the relative nucleotide substitution rates without requiring the explicit temporal information from the genomic data.

### c/μ Analysis for estimating selection type

Taking non-neutral selection into consideration, the single FGMR (*μ*) for all genes/UTRs/TRSs in GSRM is unlikely true [63]. The actual SSR of a segment (*c*_*i*_) depends on the type of selection acting on it. The three cases are thus:

SSR is reduced least if more NT sites are under nearly neutral selection:

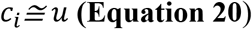

SSR is reduced most if more NT sites are under negative selection:

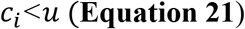

SSR is increased most if some NT sites are under positive selection:

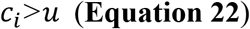

It is crucial to obtain the SSR of each segment for understanding the evolutionary dynamics of the SARS-CoV-2 virus [64]. It is worth noting that a higher substitution rate under positive selection than that under neutral selection is possible (*c*_*i*_>μ) due to the high fixation probability (*P*_*fix*_) through fixation of adaptive substitutions in the population.

Besides *c/μ*, selection type can also be determined using conventional Ka/Ks, where multiple methods can be used (see later in the methods section for our five selected Ka/Ks methods). However, Ka/Ks can only be used in the coding regions of a genome, whereas *c/μ* can be used in both coding and non-coding regions. Nonetheless, *c/μ* and Ka/Ks can be determined from the experimental sequence data, thus the selection type assignment from them is operational and objective. In contrast, a selection coefficient (*s*) is commonly used in population genetics to characterize the relative fitness for defining the selection type. Although it is theoretically important, empirical determination of such a quantity can be very difficult.

To use *c/μ*, the FGMR (*μ*) must first be determined. Under NNBST, GSR is a constant, which could be a good approximation for FGMR. In-vitro assays or in-vivo cell lines [64] without selection type might be used in the future to experimentally determine FGMR. In this study, the calculated GSR of SARS-CoV-2 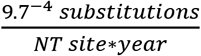 was used to represent the FGMR (*μ*) of this virus. Then, the ratio of SSR over FGMR (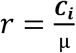) can be used to classify the selection type for a genomic segment including UTRs and TRSs that cannot be classified using conventional Ka/Ks methods. UTR/TRS is likely under near-neutral selection if *r* ≈ 1, positive selection if *r* > 1 or negative selection if *r* < 1. Because FGMR (*μ*) is always a positive number, even if *C*_*i*_ = 0 (i.e., conserved NT site), then *r* = 0 and suggests the strongest negative selection. In contrast, conventional Ka/Ks suffers if *Ks* = 0 or *Ks* ≈ 0, probably leading to a false positive classification. The SSR for each gene was also calculated. For example, the SSR for the N-gene is 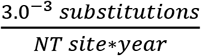, which is 3.1-fold faster than the GSR. Therefore, N-gene is under strong positive selection.

### Theoretical proof for 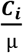 with conventional Ka/Ks in classifying selection type for a gene in TR under a near neutral condition

To validate the use of the relative substitution rate *c/μ* in classifying selection type for any given segment, we perform the following proofs against conventional Ka/Ks scheme under a near-neutral condition:

1. 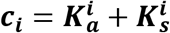, where 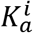 and 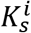 are the non-synonymous and synonymous mutation rates of gene *i* under non-neutral selection.
2. μ = ***K***_***a***_ + ***K***_***s***_ = **2*K***_***s***_, because *K*_*a*_ = *K*_*s*_ is under near neutral selection.
3. Assuming the synonymous mutation rate under neutral and non-neutral conditions is approximately equal to each other (i.e., 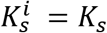).
4. Under neutral selection, 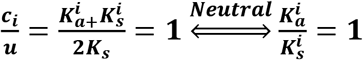
5. Under positive selection, 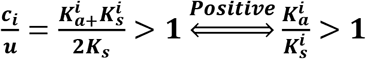
6. Under negative selection, 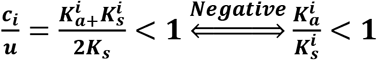

### Compilation of three genomic datasets

11,198 SARS-CoV-2 genomes collected between 12-24-2019 and 7-31-2021 were obtained from GISAID [59] (**Table 1**) and were from three independent datasets (A1a/3856, A1b/3416 and A1c/3926). These were selected and compiled by Nextstrain [40] and overlapping sequences between each dataset were removed prior to our analysis. Criteria for genome selection includes genome completeness (>27 kilobyte sequence length), no clustered substitutions and no excess divergence. The first SARS-CoV-2 sequence isolated from a human patient, Wuhan-Hu-1 (GenBank ID: MN908947.3, RefSeq ID: NC_045512.2), was used as the reference genome.

### Multiple Sequence Alignment (MSA) of genome, gene and protein sequences

MSA of all genomes in each dataset against the reference genome sequence was performed using MAFFT [65] under default settings. Aligned sequences were then ordered chronologically by their collection date.

### Extraction, compilation and MSA of gene and protein sequences

Following MSA, MATLAB Bioinformatics Toolbox was used to extract each coding gene and each non-coding UTR/TRS segment and to translate the coding genes into their respective protein sequences (**Tables S1A-S1D**) using the GenBank header file (GenBank ID: MN908947.3). MSA of each gene and protein was also done using MAFFT under default settings. Example alignments of non-redundant E-gene sequences were generated using Jalview (**Figures S2a-S2c**) and non-redundant E-protein and S-protein were generated using MultiAlin [66] (**Figures S3-4**).

### Calculating percent NT type and NT substitutions in genome, segments and codon position

The percentage of each NT type (Adenine (A), Thymine (T), Cytosine (C), Guanine (G)) in the reference genome sequence was calculated using the MATLAB Bioinformatics Toolbox script [66]. (**Table S4A**). Over the combined three datasets, total NT substitutions were decomposed into 4 transition types (purine-to-purine, pyrimidine-to-pyrimidine) and 8 transversion types (purine-to-pyrimidine and vice-versa) for the genome, each gene sequence combined (All-TR, All-UTR, All-TRS) and each individual gene (**Table S4B**). The percentage of NT substitutions occurring in positions 1, 2 and 3 (P1, P2 and P3) of each codon for All-TR and each coding gene was also calculated (**Table S4C**).

### Calculating percent amino acid (AA) type and codon substitutions

The percentage of each AA type in All-TR of the reference genome sequence was calculated using the MATLAB Bioinformatics Toolbox script [66] (**Table S5A**). Over the combined three datasets, AA substitution matrices for the 20 common AAs (20×20) were also generated for All-TR (**Table S5B**). Diagonal numbers represent synonymous substitutions and off-diagonal numbers represent non-synonymous substitutions.

### Calculating c/μ for the genome and each gene over the combined data sets

The percent substitution rate for segment *i* (*ci*) was calculated using the MATLAB Bioinformatics Toolbox script [66] to implement the following equation:

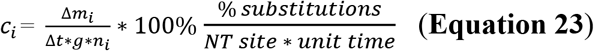

while dividing the number of substitutions (*Δm_i_*) with the number of genomes (*g*), the time interval (*Δt*) and the number of NTs (*n_i_*) within the segment i. For the genome and each segment, the percent NT substitution rate (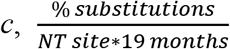) with reference to the genomic substitution rate (*μ*) and the relative substitution rate (*c/μ*) were obtained. *c* and *c/μ* values have been tabulated (**Table 3**).

**Table 2.** Percent total NT substitutions in codons containing 1, 2 or 3 NT substitutions in All-TR from the combined data sets.

| #NT Substitutions per Codon | 1 | 2 | 3 |
| --- | --- | --- | --- |
| %Total NT substitutions | 92.3 | 4.9 | 2.8 |

**Table 3.**
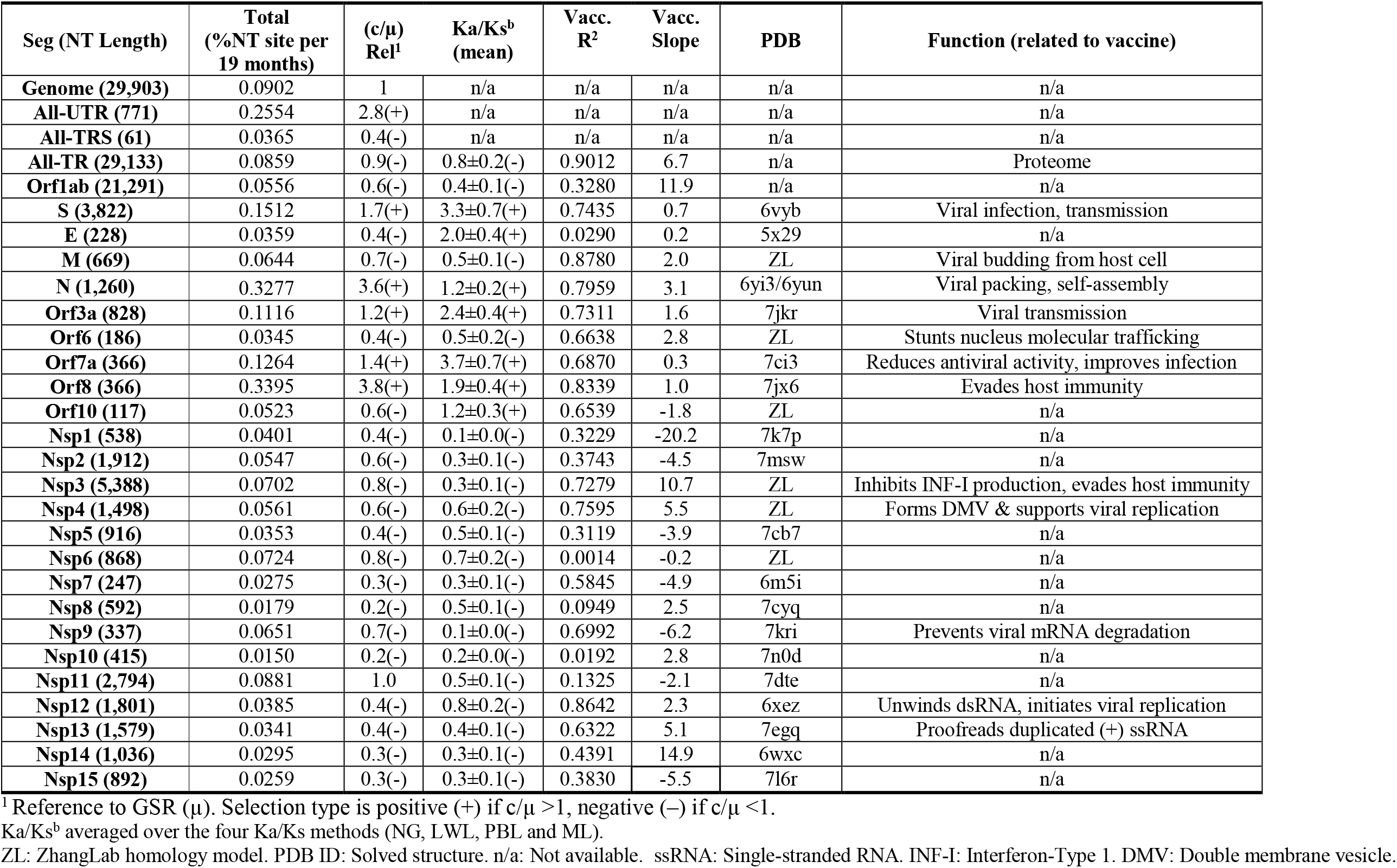
Total NT substitution rate, relative substitution rate (c/μ, Rel^1^), implied selection type (+/−) based on c/μ and Ka/Ks analysis, correlation coefficient (R^2^) and slope between the Ka/Ks timeline and the vaccination timeline (**Fig. 10-11**), protein structure model and associated functions of SARS-CoV-2 proteins.

### Calculating probability distribution of c/μ for each NT site in the genome and each gene

The probability distribution of *c/μ* was calculated over the whole genome and each segment on the combined datasets using the MATLAB Bioinformatics Toolbox script [66]. First, the distribution of the total NT substitution rate (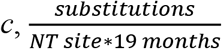) for the whole genome and each segment was obtained by calculating the substitution rate (*c*) at each NT site over all genomic sequences, followed by counting the occurring frequency of each *c*. Second, the mean total NT substitution rate within the whole genome (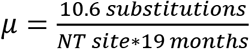) was obtained. Third, *c/μ* distribution plots were created by scaling *c* to *c/μ* for the probability distribution of the absolute substitution rate *c*. The *c/μ* probability distribution for the genome, All-UTR, All-TRS (**Figure 4C-F)**, UTRs and TRSs (**Figure S6**) are shown.

Within All-TR, we calculated the mean total NT substitution rate, mean synonymous and mean non-synonymous NT substitution rate using **Equation 15**. The NT *c/μ* distribution for the relative total NT substitution rate, relative synonymous and non-synonymous NT substitution rate was calculated for each coding gene (**Figure S7**) using the combined datasets. To compare with Ka/Ks analysis, the codon *c/μ* distribution for the relative total NT substitution rate (substitutions per codon per 19 months), relative synonymous and non-synonymous NT substitution rates were calculated for All-TR (**Figure S8**) and each gene (data not shown). Due to the averaging effect over three NTs, the codon-based *c/μ* distribution appeared flatter compared to the NT-based *c/μ* distribution (**Figure S8**).

### Selection type classification using c/μ at the NT and codon-level

A NT site in UTR or a codon site in TR can be classified as conserved (*c/μ*=0) or non-conserved (*c/μ*>0). Non-conserved NT sites can be further classified into near-neutral selection (0.5<*c/μ*<2.0), which decomposes into weak negative selection (0.5<*c/μ*<1.0) or weak positive selection (1.0<*c/μ*<2.0). Otherwise, NT sites are classified as not near-neutral and undergo either strong negative selection (0<*c/μ*<0.5) or strong positive selection (*c/μ*>2.0). The percentage of the sites under different selection types were calculated for All-TR and each major/accessory gene (**Table S6A**), Nsp1-15 (**Table S6B**), UTR (**Table S6C**) and TRS (**Table S6D**).

### Calculating Ka, Ks and Ka/Ks using five methods

Calculation of Ka, Ks and Ka/Ks was done using the MATLAB Bioinformatics Toolbox script [66]. We evaluated the consistency of Ka, Ks and Ka/Ks values, by calculating them using five different methods: Single Point mutation approximation (SP), Nei-Gojobori (NG) [67, 68], Li-Wu-Luo (LWL) [69], Pamilo-Bianchi-Li (PBL) [70] and the maximum likelihood (ML) methods [71]. SP is based on the assumption that most NT substitutions in TR are single-point substitutions in the codon, such that a point substitution can be easily classified as a non-synonymous substitution (codon change causes AA change), or a synonymous substitution (no codon/AA change). NG is the default method and is based on the Jukes-Cantor model [60] to calculate the number of Ka and Ks substitutions as well as the number of Ka and Ks sites [67]. LWL is based on the Kimura two-parameter model [72], which assumes equal NT base rates and calculates the number of Ka NT transition and transversion substitutions at three levels of codon degeneracy [69]. PBL performs similarly to LWL while including bias correction [70]. ML uses the Goldman-Yang method [73] to calculate Ka and Ks substitutions by accounting for NT transition/transversion substitution rate and NT base/codon rate biases [71].

### Selection type classification using Ka/Ks

Classification of selection type is defined for conserved (*Ka* = *Ks* = 0), strong negative (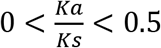), weak negative (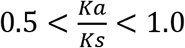), weak positive (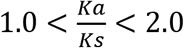) and strong positive (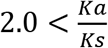) selection. For All-TR and each coding gene, Ka, Ks and Ka/Ks ratios were calculated using the five methods (SP, NG, PBL, LWL and ML) across each dataset. Averaged Ka/Ks values over the three datasets (**Table 3** and **Table S7-S8**) and Ka, Ks and Ka/Ks values for each dataset (**Table S8**) and for each of the four methods (NG, PBL, LWL and ML) were tabulated (**Tables S9A-S9D**).

### Selection type classification at the codon-level

For All-TR, the Ka/Ks ratio for each codon was calculated using three methods (NG, PBL and LWL) with a sliding window of 45 codons. Ka and Ks values were averaged over all sequences before the final Ka/Ks ratio was calculated. The percentage of codons following negative selection (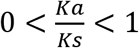) or positive selection (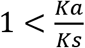) was calculated for All-TR, each major/accessory gene and Nsp1-15 using the Ka/Ks data from the three methods (**Tables S10A-S10D**).

### Ka/Ks Time-dependency correlation plots

To probe the time-dependency of Ka/Ks for All-TR and each gene/Nsp, its monthly value was calculated using the SP and NG method on the combined three datasets. Correlation plots were generated for each major/accessory gene (**Figure S12**) and for Nsp1-15 (**Figure S13**) using the MATLAB Bioinformatics Toolbox script [66].

### 1-Dose vaccine correlation plots

To probe the correlation between selection type change and 1-dose vaccinations, the timeline of vaccinations is included in NG Ka/Ks vs time for each major/accessory gene (**Figure 10**) and for Nsp1-15 (**Figure 11**) using the MATLAB Bioinformatics Toolbox script [66]. Correlation plots including both NG and SP Ka/Ks methods are included in the supporting document for major/accessory genes (**Figure S12**) and for Nsp1-15 (**Figure S13**). For UTR and TRS, vaccination timelines are included in the percent total NT variation vs time plots (**Figure S11**). The correlation coefficient, intercept and slope were calculated for each major/accessory gene and Nsp1-15 (**Figure S15**) and UTR and TRS (**Figure S14**) and were tabulated (**Table 3** and **Table 4**).

**Table 4.**
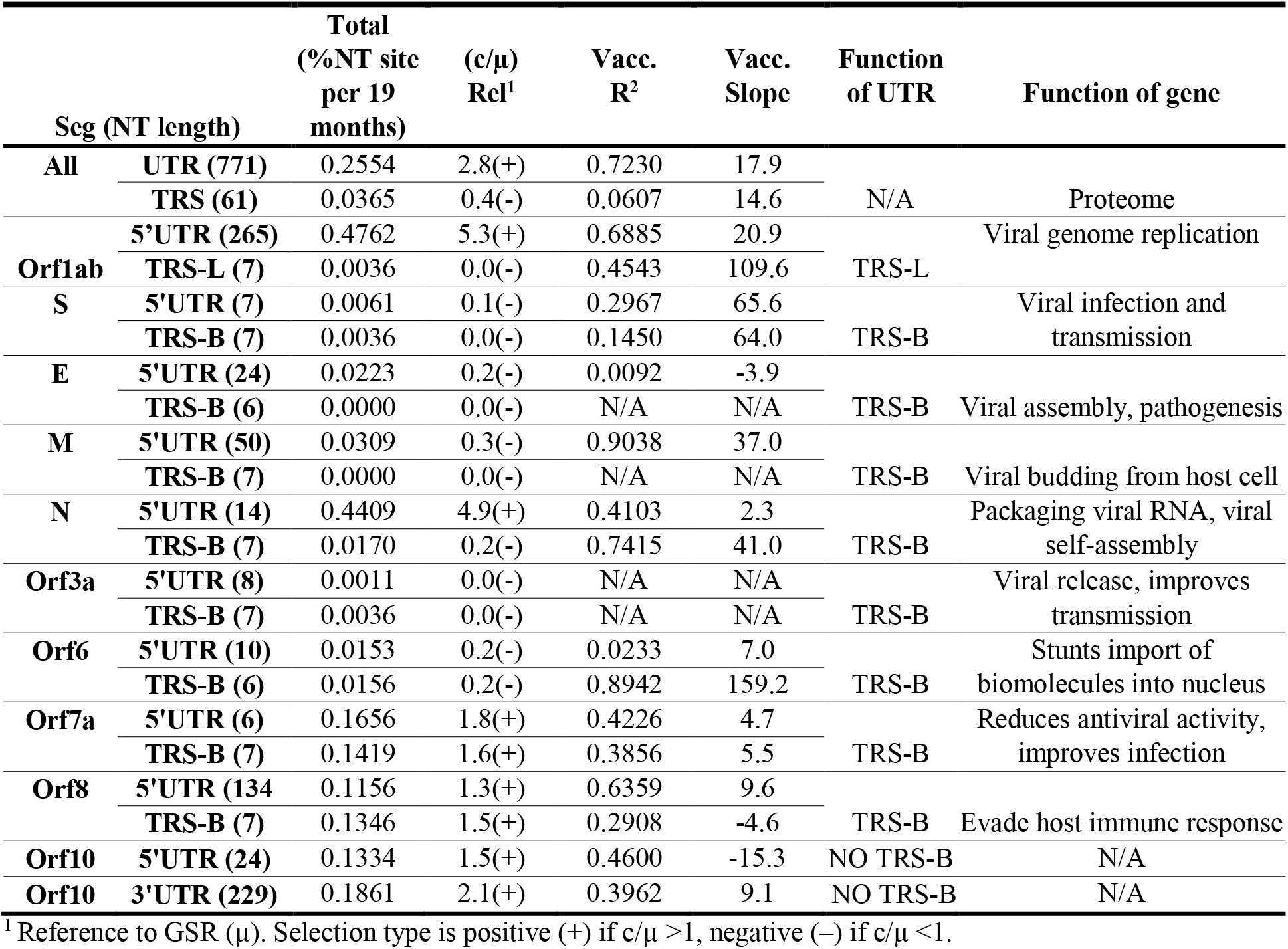
Total NT substitution rate, relative substitution rate (Rel^1^), implied selection type, correlation coefficient (R^2^) and slope between the total NT variation timeline and 1-dose vaccination timeline (**Fig. S11 and S14**), and UTR/TRS function where applicable.

### Protein structure models

Protein homology structure models were obtained from ZhangLab, in which they generated using their protocol [74]. The top 247 AA sites under positive selection based on *c/μ* and Ka/Ks analysis were mapped to each protein structure model using VMD v1.9.3. (**Figure 9 and Figure S10**). Protein secondary structure assignment is obtained from the Stride program in VMD.

### Data Files

The metadata including GISAID ID, collection date etc., the 9,711 AA sites in the proteome (translated All-TR) sorted by *c/μ* and by Ka/Ks with the SP method, the top 247 AA sites under positive selection (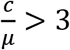 and 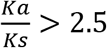), the 771 NT sites in UTR sorted by *c/μ* and top 54 NT sites under positive selection (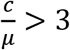) are provided in supporting Excel spreadsheet files.

## Results

### GSRM and KNT were not followed at each NT type in the genome

To demonstrate that the GSRM and KNT were not followed at the NT-level, we calculated the substitution rates for the four types of NTs (A, G, U and C), the 4 transition types and 8 transversion types in the genome and each segment as described in the methods section (**Table S4**). We otherwise would expect each NT type to exhibit similar transition/transversion substitution rates. The NT abundances in the SARS-CoV-2 reference genome (A:30%, G:20%, T:32% and C:18%) (**Table S4A**) showed high consistency with NT abundances for SARS-CoV and MERS-CoV [75]. Uneven substitution rates among the transition and transversion types were observed; Indeed, very high C-U transition substitution rates were exhibited in the genome (136,701 C-U substitutions, 4 to 6-fold higher than A-G, G-A and U-C substitutions) and each gene (e.g., S-gene: 13,142 C-U substitutions, 2 to 7-fold higher than G-A and U-C substitutions). The high C-U transition substitution rates might be due to a host-driven antiviral mechanism; human APOBEC deaminases induce C-U transition substitutions in foreign single-stranded DNA or RNA, driving C-U hyper mutations in SARS-CoV-2 genomes [76, 77]. Moderate G-U transversion substitution rates were also exhibited in the genome (39,241 G-U substitutions, 2 to 10-fold higher than the other 7 transversion types) and only some genes (e.g., S-gene: 5,502 G-U substitutions, 11-fold higher than U-A substitutions) (**Table S4B**). From the uneven transition and transversion substitution rates, GSRM and KNT were not followed at each NT type. Admittedly, it’s unclear whether ONNT or NNBST are applicable here, as the percentage of near-neutral selection types were calculated later.

### GSRM and KNT were not followed at each position of codon and at each codon type in All-TR

To demonstrate that GSRM and KNT were not followed at the codon-level, we calculated the percent NT substitution rates within positions 1, 2 and 3 (P1, P2 and P3) in each codon and averaged over each codon in All-TR (**Table S4C**). Otherwise, we expect the substitution rate to be similar at P1, P2 and P3 and within each codon. P2 had the highest percent substitution rate throughout the genome (44%), whereas P1 and P3 had comparable percent substitution rates (28%) (**Table S4C1**). Indeed, P2 had higher percent substitutions for all genes except Orf1ab (17%), Nsp1 (0%), Nsp2 (1%), Nsp3 (6%) and Nsp9 (0%) (**Table S4C1 and S4C2**). Typically, P1 and P2 should contain the most non-synonymous substitutions, whereas P3 (a.k.a., the wobble site) should contain the most synonymous substitutions. [78, 79] Therefore, P3 typically had a higher substitution rate than P1 and P2. Additionally, 93.3% of total NT substitutions occurred as single-point substitutions, 4.9% as 2-point substitutions and 2.8% as 3-point substitutions. This result supported the use of the four elaborate methods (NG, LWL, PBL and ML) rather than SP for getting more accurate estimates on Ka, Ks and Ka/Ks values. We then calculated the number of synonymous and non-synonymous NT substitutions for all possible 20*x*20 AA combinations (**Table S5**) using the SP method. If GSRM or KNT were assumedly followed, we expect the number of synonymous and non-synonymous NT substitutions to be the same (GSRM: Ka/Ks=1; KNT: Ka/Ks≈1). The calculated AA abundance ranged from 1% to 9% in the reference genome (**Table S5A**). Interestingly, highly uneven substitution rates at the codon level were observed (e.g., T-I, 7% of the total substitutions, **Table S5B**), clearly indicating GSRM and KNT were also not followed at the codon-level.

### GSRM and KNT were not followed at NT site level and at each coding gene and non-coding segment

To demonstrate that GSRM and KNT were not followed at the NT site-level and the segment-level, a diagram showing the percent NT substitutions at the whole genome, All-TR, All-UTR and each gene was generated (**Figures 3B-3H**). Total NT substitutions in TR (**Figure 3E**) were decomposed into non-synonymous (**Figure 3F**) and synonymous substitutions using the Ka/Ks SP method (**Figure 3G**); the Ka/Ks ratio is shown in **Figure 3H**. If GSRM is true, the substitution rate at every NT site should be similar. In fact, while 22% of NT sites showed no substitution, 78% NT sites showed variable substitution rates (**Figure 3B**). Thus, GSRM is not true for this virus. Although KNT allows conserved sites due to negative selection, the substitution rate at each substituted site should nonetheless be similar. Instead, a variable distribution of NT substitutions was observed in the genome and each gene segment. Many NT substitutions were observed in the genome (317,828 total NT substitutions), TR (294,631, ~93% from genomic NT substitutions), Orf1ab-gene (139,479 substitutions, ~44%), S-gene (68,050 substitutions, ~21%) and N-gene (48,627 substitutions, ~15%) (**Figure 3B**). High NT substitution rates were observed in UTR, primarily in Orf1ab 5’UTR (**Figure 3D**). Thus, KNT was not followed, Additionally, synonymous and non-synonymous NT substitution rates in TR should also be similar for KNT (KNT: Ka/Ks≈1). In Orf1ab-gene, the amount of NT substitutions that were synonymous (63,990, ~46%) were slightly less than non-synonymous substitutions (75,489, ~54%), whereas most NT substitutions in non-Orf1ab-genes were non-synonymous (138,697, ~89%), indicating that Nsp1-15 in Orf1ab are critical for maintaining the SARS-CoV-2 life cycle. This Ka/Ks analysis also suggests that KNT is not true for the TR of this virus.

**Figure 3.**
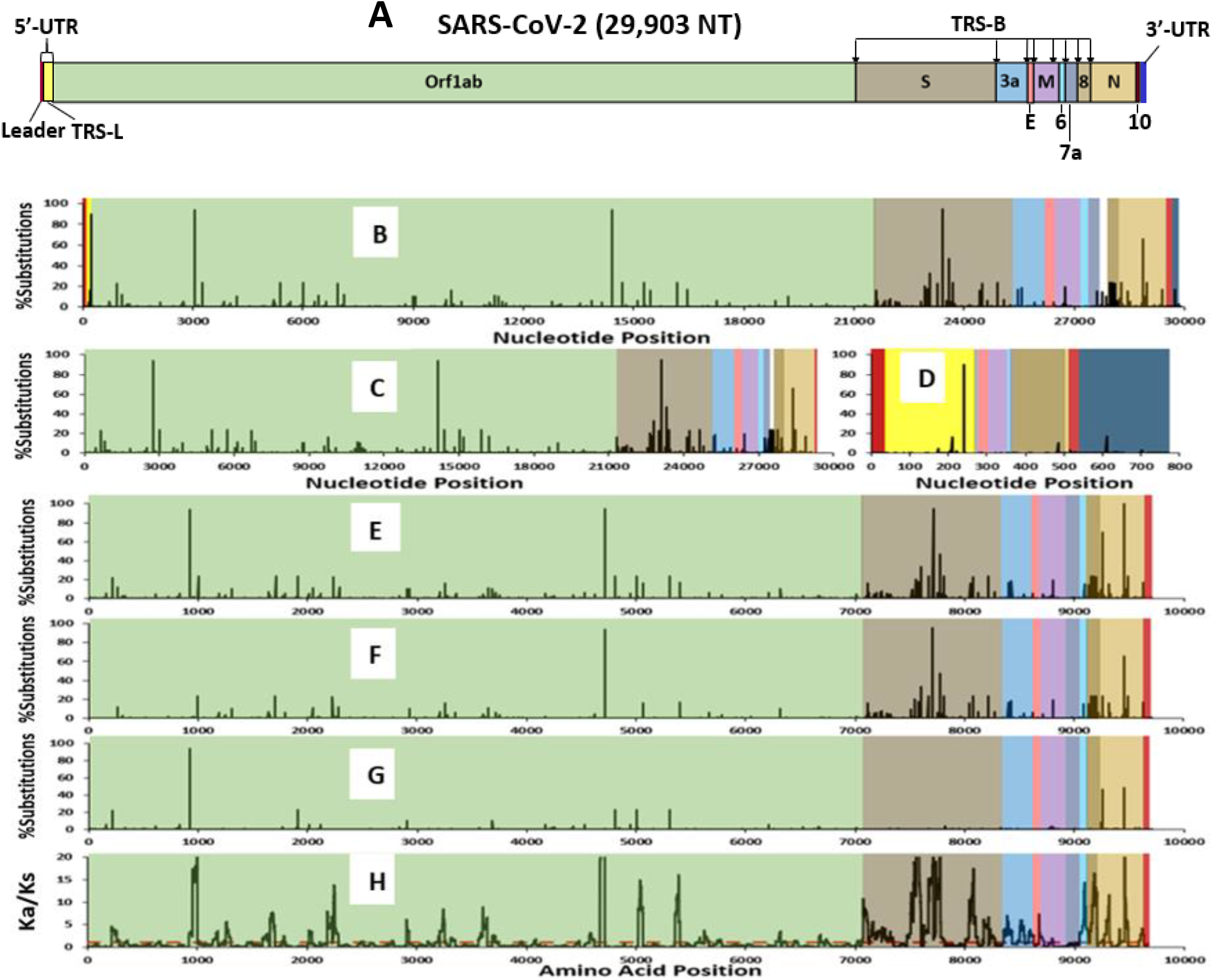
The genome structure of SARS-CoV-2 (**A**) and its substitution distribution (**B-H**) observed in the combined three data sets. (**A**) Major gene segments within the SARS-CoV-2 human reference genome (Wuhan-Hu-1). Total genome, NT, protein, UTR, TRS-L and TRS-B sequence lengths have been tabulated (**Table S1**). (**B**) Percent total NT substitutions across the entire genome, each gene and the 5’- and 3’-UTRs. (**C**) Percent total NT substitutions for each gene in TR and (**D**) their corresponding 5’-UTR and 3’-UTR regions. (**E**) Summed percent NT substitutions within three positions of each codon. (**F**) Percent non-synonymous NT substitutions in TR. (**G**) Percent synonymous NT substitutions in TR, calculated from the SP method. (**H**) Ka/Ks distribution across TR using NG method with a sliding window of 45 codons. The red dotted line in **(H)** represents the cutoff for residues undergoing positive, neutral or negative selection (Ka/Ks = 1).

**Figure 4.**
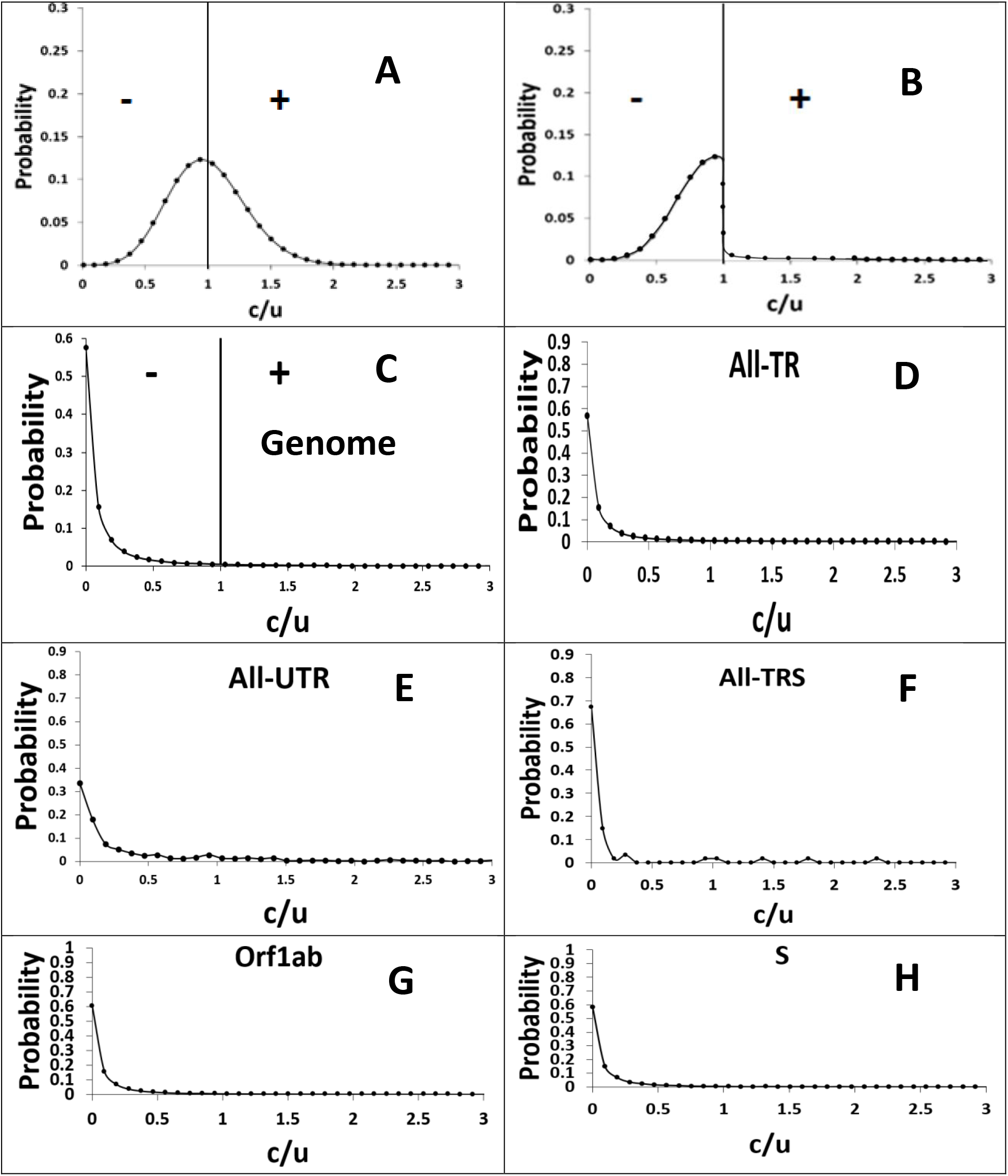
Comparison between the hypothetical probability distributions (**A-B**) and the empirical L-shape probability distribution (**C-H**) of relative NT substitution rates (*c/μ*). **A**: Given *μ* = 10.6 substitutions per NT site per 19 months for the genome, the hypothetical Poisson probability distribution implied by the NMT was calculated using the Poisson probability mass function equation for a genome containing x substitutions: 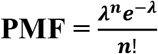, where *λ* = μ * *t* = 11, *t* =19 *months* and n is a whole integer and has the range [0, the NT segment length]. **B**: Asymmetric distribution implied by the NNT. **C-H**: The empirical L-distribution of the relative NT substitution rate for the genome, All-TR, All-UTR, All-TRS, Orf1ab and S of SARS-CoV-2, supporting our NNBST. **A-C**: Probability distribution showing negative (−), positive (+) and neutral selection (straight line).

Inspection of each coding gene and protein sequence, such as non-redundant sequences (i.e., non-overlapping mutant sequences) of the E-gene (**Figures S2a-S2c**) and E-protein and S-protein (**Figures S3a-S3c and Figures S4a-S4c**) confirmed random substitution sites but also uneven substitution rates across all sites. Interestingly, most NT substitutions in the E-gene were translated into non-synonymous substitutions in the E-protein (**Figures S3a-S3c**). Position 71 of the E-protein had the highest rate of non-synonymous substitutions (~5.00% per site per 19 months) of Pro71Lys, Pro71Phe or Pro71Ser substitutions. The S-protein exhibited higher substitution rates at some sites (**Figures S4a-S4d**), such as positions 501 (**Figure S4b**, ~25%) and 681 (**Figure S4c**). The variations in NT substitution rates and in Ka/Ks in each gene strongly indicates the GSRM and KNT were not followed at the gene-level and that near neutral and non-near neutral selection types must cause this. What percentage of the NT sites are near neutral or not-near neutral? Are near neutral sites solely slightly deleterious or also slightly positive?

### L-shaped c/μ probability distribution suggests that NNBST rather than ONNT is followed

To determine the selection type for each NT site and the relative abundance of each selection type, we generated probability distribution curves of the total relative NT substitution rate (*c/μ*) for the genome, All-TR/All-UTR/All-TRS and each coding/non-coding segment (**Figure 4**). The genomic *c/μ* probability distribution was then compared to the implied distribution curves assuming KNT and ONNT with a peak probability at *c/μ* = 1 (**Figure 4**). The KNT should exhibit a symmetric Poisson distribution curve, whereas the ONNT should exhibit an asymmetric distribution curve in which the area under the curve is significantly smaller for weak positive selection (right of the curve) than for weak negative selection (left of the curve), indicating the nearly neutral sites are deleterious. It’s important to note that the probability of conserved sites (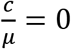) was included in these distribution curves. Intriguingly, an L-shaped distribution curve rather than a Poisson distribution curve was observed for the genome, All-TR, All-UTR, All-TRS, Orf1ab and S (**Figures 4A, C-H**) and all segments (**Figures S6-S8**). Here, this suggests that most NT sites are conserved (Genome: 60%; All-TR: ~57%; All-UTR: ~33%; All-TRS: ~67%; Orf1ab: ~60% and S: ~60%) or some are under strong negative selection, whereas weak negative, weak positive and strong positive selection is increasingly rare. Additionally, the distribution of the relative synonymous/non-synonymous NT substitution rates were also calculated for All-TR and each gene (**Figures S7-S8**), which too exhibited L-shaped distributions. The mean absolute substitution rate (*μ*) for All-TR was calculated for total NT substitutions (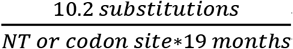), synonymous NT substitutions (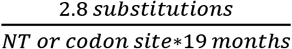) and non-synonymous NT substitutions (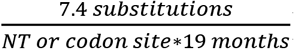). This L-shaped distribution clearly is consistent with NNBST, and it also suggests that the GSRM, KNT and the ONNT are not followed for the genome and each segment.

### The percentage of the positive and the negative selection types confirmed NNBST for this virus

To demonstrate that the KNT and ONNT are not followed in another way, we calculated the percentage of sites undergoing near neutral or not-near neutral selection for the genome and each segment using the *c/μ* test as defined in the methods section (**Table S6**). Pie charts depicting the percent selection types in All-TR, All-UTR and All-TRS using *c/μ* were also generated (**Figure 5A-C**). Assuming the ONNT is followed, we expect an asymmetric abundance of near neutral selection types (weak negative selection only). However, All-TR, All-UTR and All-TRS showed a similar abundance of both weak negative and weak positive selection types across their sites. Most sites in All-TR, All-UTR and All-TRS were under strong negative selection. For All-TR, 78% of sites were non-conserved (61% strong negative, 8% weak negative, 4% weak positive and 5% strong positive selection) (**Table S6A**). Orf1ab values were highly consistent with All-TR (77% non-conserved: 63% strong negative, 7% weak negative, 4% weak positive and 3% strong positive selection) (**Table S6A**). Coincidentally, non-Orf1ab genes were slightly less conserved (83% non-conserved: 59% strong negative, 11% weak negative, 5% weak positive and 8% strong positive selection) (**Table S6A**). For Nsp1-15, the values were highly consistent with Orf1ab (76% non-conserved: 62% strong negative, 7% weak negative, 3% weak positive and 4% strong positive selection) **(Table S6B**). For All-UTR, less sites were non-conserved (66% non-conserved: 63% strong negative, 2% weak negative, <1% weak positive and 1% strong positive selection) (**Table S6C**). For All-TRS, even less sites were conserved (33% non-conserved: 30% strong negative, 3% weak negative, <1% weak positive and <1% strong positive selection) (**Table S6D**). Although the combined percentages of near neutral selection are relatively small in All-TR (~12%), All-UTR (<3%) and All-TRS (<4%), the percentage of sites under weak negative and weak positive selection were mostly consistent within All-TR (4% - 8%), All-UTR (<1% - 2%) and All-TRS (<1% - 3%). Obviously, these data suggest NNBST is followed instead of ONNT or KNT.

**Figure 5.**
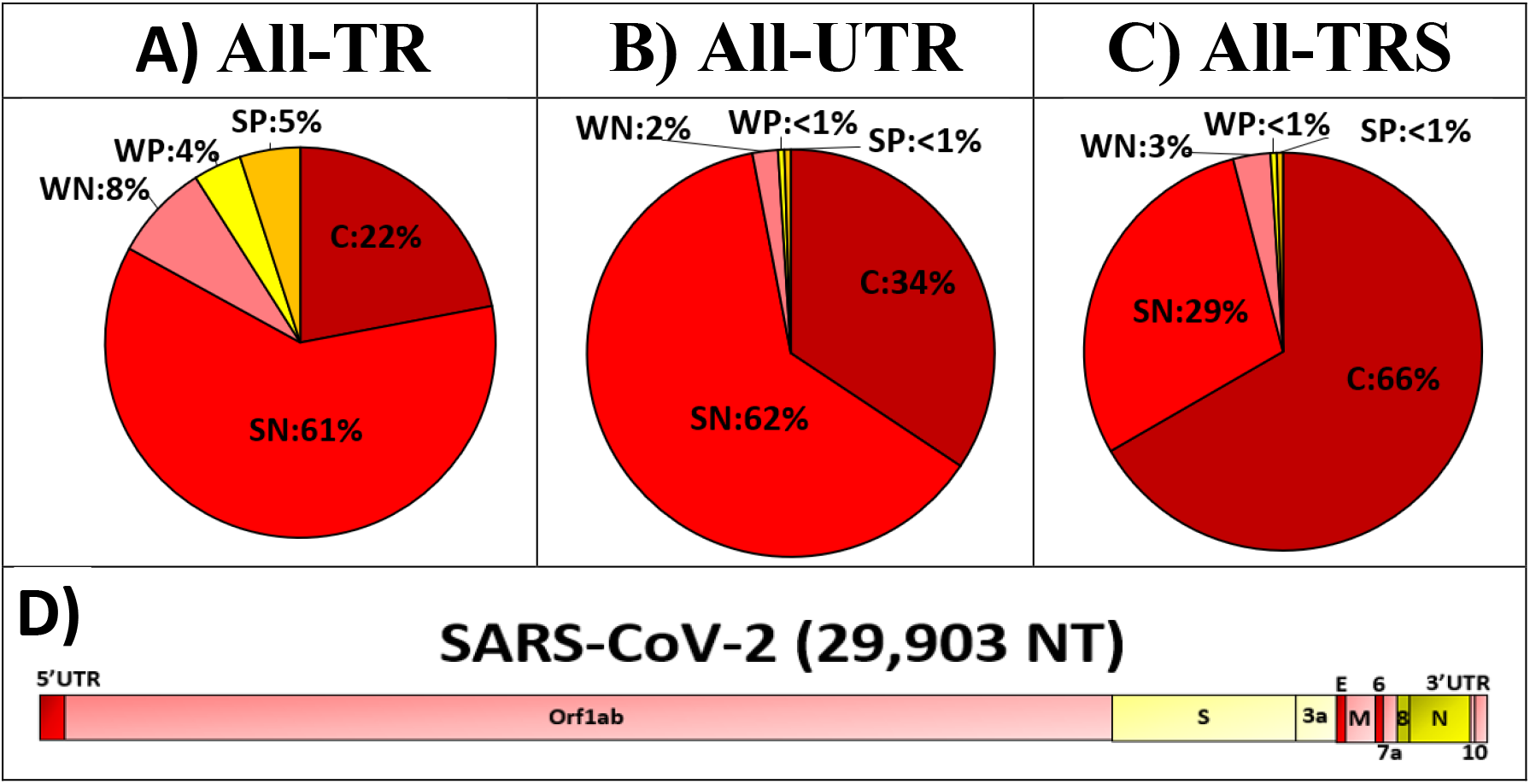
The molecular evolution nature of SARS-CoV-2. **A**-**C**. Percent decomposition of NT mutations into conserved selection (C, *c/μ* = 0, dark red), strong negative selection (SN, 0 < *c/μ*< 0.5, red), weak negative selection (WN, 0.5 < *c/μ* < 1.0, light red), weak positive selection (WP, 1.0 < *c/μ* < 2.0, light yellow) and strong positive selection (SP, *c/μ* > 2.0, dark yellow) for All-TR (**A**), All-UTR (**B**) and All-TRS(**C**). **D**: Overall nearly neutral selection on SARS-CoV-2 genome, produced by the balancing between weakly negative selection regions (Orf1ab, M, Orf7a, Orf10 and Orf10 3’UTR in light red), weakly positive selection regions (S and Orf3a in light yellow) and strongly negative selection regions (Orf1ab 5’UTR, E and Orf6 in dark red) and strongly positive selection regions (Orf8 and N in dark yellow).

### Varying relative substitution rates in each segment observed in TR and UTR, supporting NNBST

Some genes exhibited lower or higher percent NT substitutions across the genome and thus likely exhibit slower or faster substitution rates (*c*, SSR) relative to the genomic substitution rate (*μ*, GSR). By taking the ratio of these two values (*c/μ*), selection pressure types can be assigned to any coding or non-coding gene region similarly to conventional Ka/Ks methods (see the methods section).

To characterize the different selection types for each gene region, the total NT substitution rates for the genome (GSR, *c*), each segment (SSR, *c*), the relative segment substitution rate (Rel^1^, *c/μ*) and selection type classification based on the *c/μ* test were tabulated (**Table 3-4** and **Figure 6-7**). The overall average percent NT substitution rate for the GSR is 0.0902% (Rel^1^ = 1). Assuming GSRM and KNT are followed, the SSR of each gene should be consistent with the GSR (GSRM: *c = μ*; KNT: *c ≈ μ*), suggesting neutral selection. Assuming ONNT, we expect to observe slight variations in the SSR owed to mainly weak negative selection with very rare weak positive selection. Rel^1^ values suggested that All-UTR (2.8) is under strong positive selection, All-TR (0.9) is under weak negative selection and All-TRS (0.4) is under strong negative selection. For the major genes (**Table 3-4** and **Figure 6-7**), N (3.6) is under strong positive selection, S (1.7) is under weak positive selection, M (0.7) and Orf1ab (0.6) are under weak negative selection and E (0.4) is under strong negative selection. For the accessory genes, Orf8 (3.8) is under strong positive selection, Orf7a (1.4) and Orf3a (1.2) are under weak positive selection, Orf10 (0.6) is under weak negative selection and Orf6 (0.4) is under strong negative selection. For Nsp1-15, Nsp11 (1.0) is under near-neutral selection and the Nsp2-4, Nsp6 and Nsp9 (0.6-0.8) are under weak negative selection and Nsp1, Nsp5, Nsp7-8, Nsp10 and Nsp12-15) (0.2-0.4) are under strong negative selection. For each UTR (**Table 3-4** and **Figure 6-7**), Orf1ab 5’UTR (5.3), N 5’UTR (4.9) and Orf10 3’UTR (2.1) are likely under strong positive selection, Orf7a 5’UTR (1.8), Orf10 5’UTR (1.5) and Orf8 and 5’UTR (1.3) are likely under weak positive selection, M 5’UTR (0.3), Orf6a 5’UTR (0.2), E 5’UTR (0.2), S 5’UTR (0.1) and Orf3a 5’UTR (0.0) are likely under strong negative selection or are strongly conserved. For each TRS (**Table 4 and Figure 7**), Orf7a TRS-B (1.6) and Orf8 TRS-B (1.5) are likely under weak positive selection and Orf6 TRS-B (0.2), N TRS-B (0.2), S TRS-B (0.0), Orf3a TRS-B (0.0), Orf1ab TRS-L (0.0), E TRS-B (0.0) and M TRS-B (0.0) are likely under negative selection or are strongly conserved.

**Figure 6.**
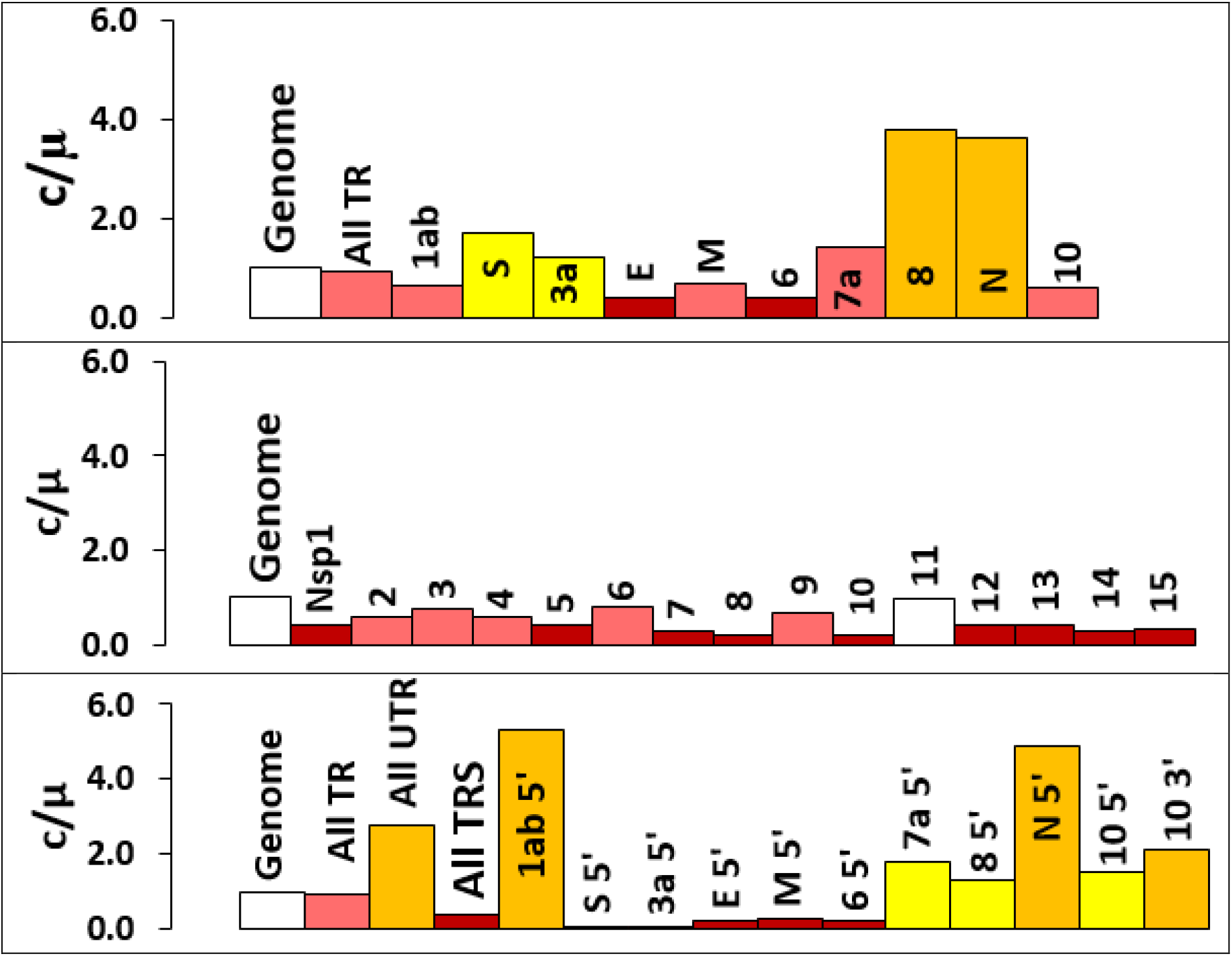
Relative substitution rate (*c/μ*) of All-TR and each gene (top), Nsp1-15 (middle) and All-UTR and each UTR (bottom). 5’ and 3’ represent 5’-UTR and 3’-UTR. Selection types are defined by color (strong negative selection (0 < *c/μ* < 0.5, red), weak negative selection (0.5 < *c/μ* < 1.0, light red), weak positive selection (1.0 < *c/μ* < 2.0, light yellow) and strong positive selection (*c/μ* > 2.0, dark yellow)).

**Figure 7.**
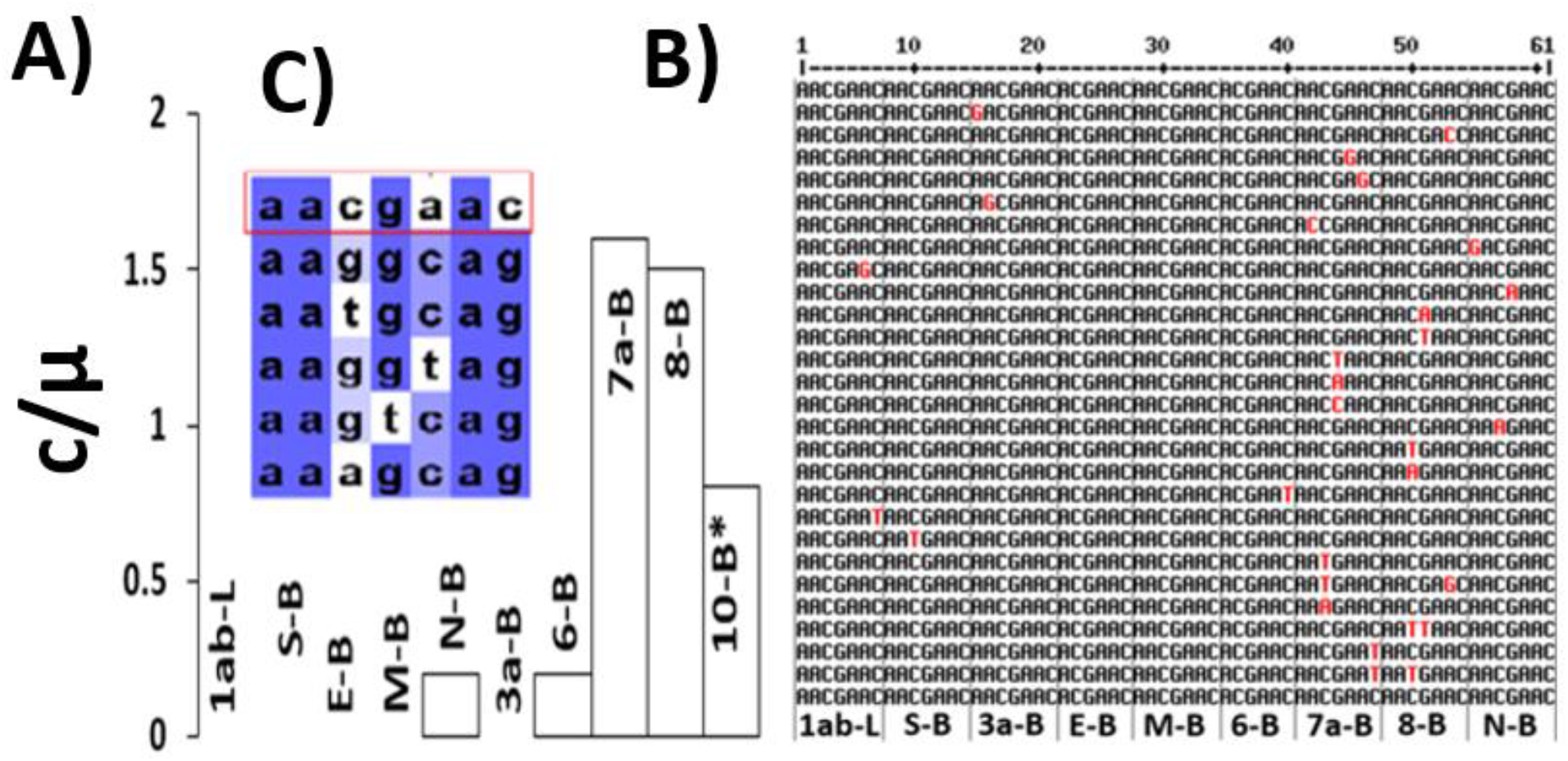
Relative substitution rate (*c/μ*) and unique sequence of each active TRS (**A-B**) and Hypothetical Orf10 TRS-B sequences (**C**). Mutations are in red font.

A mixture of near neutral and non-near neutral selection types was observed across each segment, strongly indicating that GSRM and KNT were not followed at the gene-level. Concerning non-near neutral selection, 17 gene segments exhibited strong negative or conserved selection (E, Orf6, Nsp1, Nsp5, Nsp7-8, Nsp10, Nsp12-15, M 5’UTR, Orf6a 5’UTR, E 5’UTR, S 5’UTR, Orf3a 5’UTR) and 5 gene segments exhibited strong positive selection (N, Orf8, Orf1ab 5’UTR, N 5’UTR and Orf10 3’UTR). Concerning near-neutral selection, 8 gene segments exhibited weak negative selection (M, Orf1ab, Orf10, Nsp2-4, Nsp6 and Nsp9) and 8 gene segments exhibited weak positive selection (S, Orf7a, Orf3a, Orf7a 5’UTR, Orf10 5’UTR, Orf8, 5’UTR, Orf7a TRS-B and Orf8 TRS-B). Therefore, NNBST rather than ONNT is supported by these relative substitution rates at the gene-level. In other words, the weak selection pressures combine and balance out, resulting in the overall near-linear GSR.

### *c/μ* and Ka/Ks selection type assignment was mostly consistent for All-TR

To validate the *c/μ* test, we compared our selection type assignment from the *c/μ* analysis with conventional Ka/Ks analysis by calculating average Ka/Ks values over 19 months and over each dataset using two classes of methods. The first method class (SP) assumes that only 1 NT mutation occurs within each AA codon (Ka/Ks^a^). The second method class (NG, LWL, PDL and ML) accounts for instances where ≥1 NT mutation occurs within an AA codon (Ka/Ks^b^, averaged over the four methods). Ka/Ks^a^ and Ka/Ks^b^ values for each TR segment were tabulated (**Table S7**). Ka/Ks^b^ values for each of the four methods (**Table S8**) and raw Ka, Ks and Ka/Ks value for Ka/Ks^b^ for each of the four methods and for each dataset were also tabulated (**Tables S9A-D)**. The SP method slightly overestimated Ka/Ks values (**Table S7**), whereas the other four methods exhibited very consistent Ka/Ks values (**Table S8**). Therefore, Ka/Ks^b^ values were used to classify selection type (**Table 3**).

For validation purposes, we compiled a list of seven Ka/Ks analyses from the existing literature (not including ours) to compare against our Ka/Ks^b^ results (**Table S3**) [10–16]. We observed that our Ka/Ks^b^ results for the genome and some genes (i.e., M, Orf10 and Nsp1-15) were consistent with those reported in the literature. Inconsistencies in the selection type for S, E and N-proteins were also observed. However, the time-dependency of these Ka/Ks values for S, E and N may be able to explain these inconsistencies. It must be noted that the longest time period analyzed for most of the literature studies is 13 months (Dec. 2019-Jan. 2021). From our Ka/Ks distribution over the first 13 months (**Figure 10**), S, E and N were reportedly under negative selection for most of the studies, which matches our approximate Ka/Ks^b^ within this time period. However, inclusion of the latter 6 months (Feb. 2021-July 2021) exhibited increased non-synonymous substitution rates for S, E and N and their Ka/Ks^b^ values indicated their positive selection.

Incredibly, the selection type assignment based on our *c/μ* test and Ka/Ks^b^ were consistent for each gene except E and Nsp11 (**Figure 8, Table 3)**. Based on Rel^1^, All-TR, Orf1ab, M, Orf6, Orf10, Nsp1-10 and Nsp12-15 likely undergo negative selection, whereas S, N, Orf3a and Orf8 likely undergo positive selection. From *c/μ* to Ka/Ks^b^, the selection type of E flips from negative selection to positive selection and Nsp11 flips from near-neutral selection to negative selection. Due to this observed consistency, it’s quite likely our *c/μ* test accurately assigned selection type to non-coding UTR and TRS. The *c/μ* test thus suggests that All-UTR, several UTRs (i.e., Orf1ab 5’UTR) and some TRSs (i.e., Orf7a TRS-B) likely undergo positive selection, whereas All-TRS, some UTRs (i.e., M 5’UTR) and most TRSs (i.e., S TRS-B) likely undergo negative selection.

**Figure 8.**
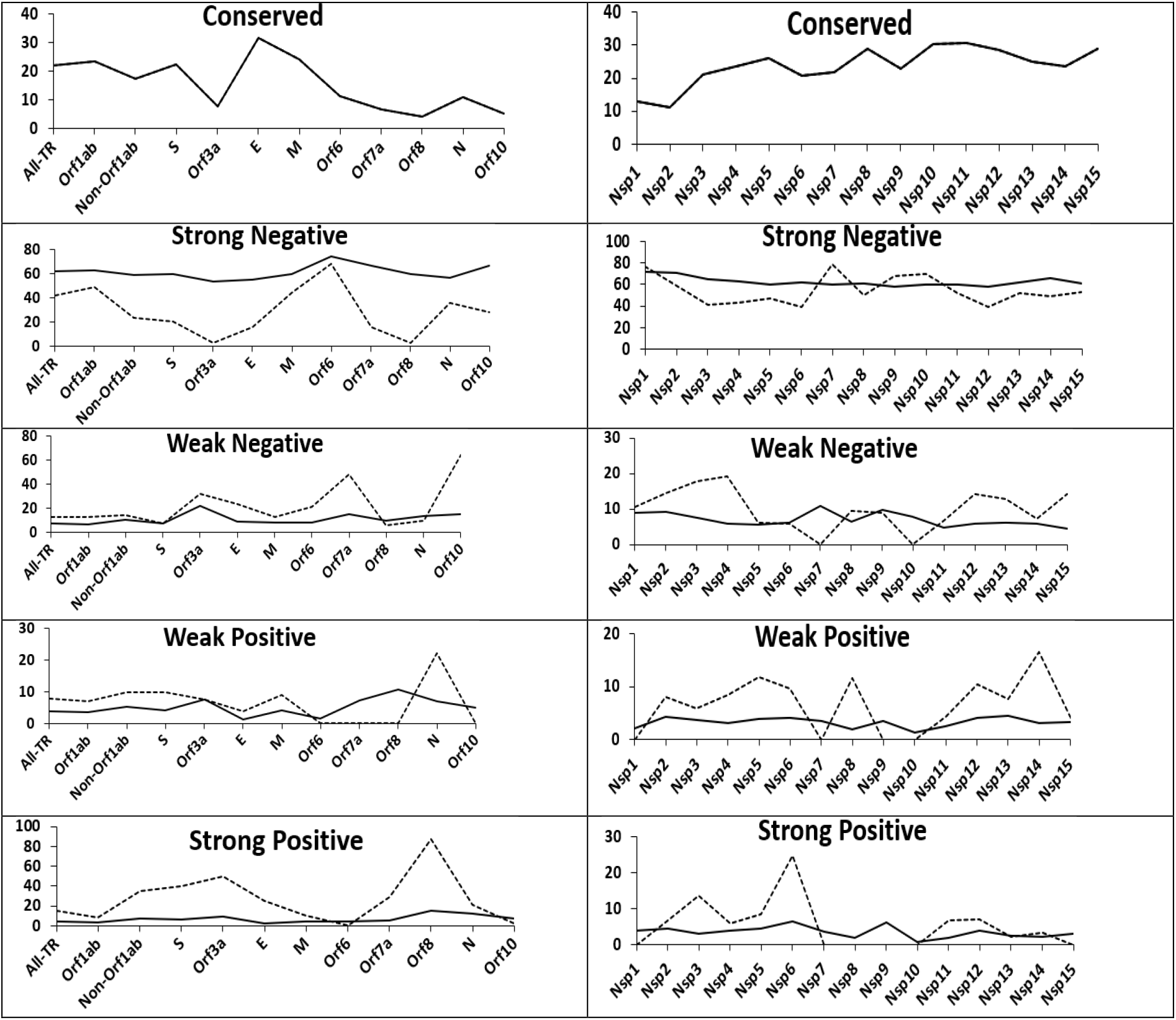
Percent decomposition of codon sites into conserved, strong negative, weak negative, weak positive and strong positive selection for All-TR, major/accessory genes (**left column**) and Nsp1-15 (**right column**) by the two methods (c/u: solid line; Ka/Ks using NG, LWL, PBL and ML: dashed line).

**Figure 9.**
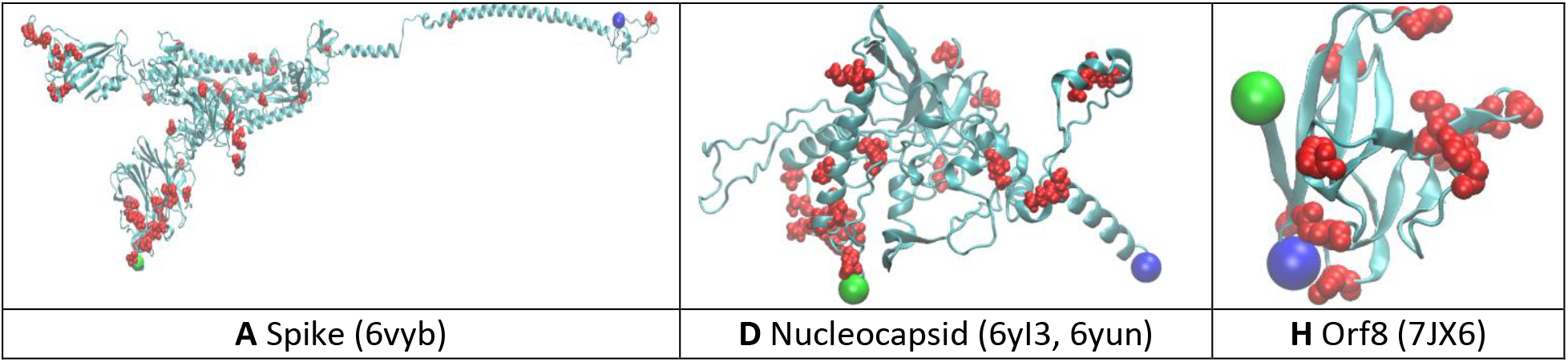
Top AA sites (red spheres) under strong positive selection for three selected proteins (See **Figure S10** for the complete set of proteins). Protein structure models were generated from the PDB databank. Protein backbone is represented as blue ribbons. N- and C-termini are represented as green and blue VDW spheres, respectively.

**Figure 10.**
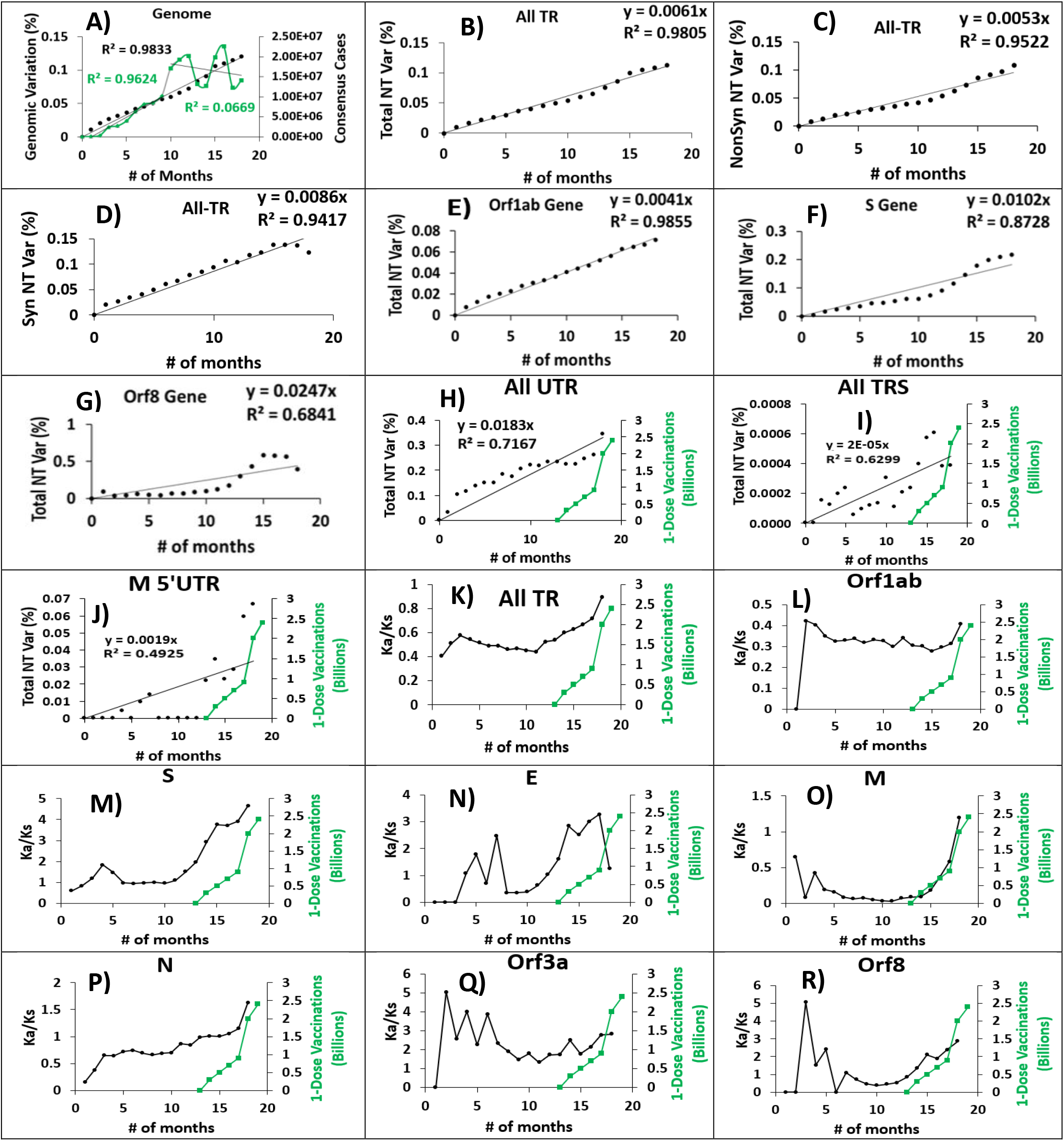
Timelines for the Genome, All-UTR, All-TRS, M 5’UTR, All-TR, the major and accessory genes from the combined sets**. A**: the monthly total NT sequence variations of the genome along the evolution time and the census population (green line). Green font represents vaccination data, black font represents genomic variation. Show Slope **B-D**: the monthly sequence variations in All-TR including the total NT substitutions (**B**), non-synonymous NTsubstitutions (**C**) and synonymous NT sequence (**D**). Non-synonymous and synonymous NT substitutions were calculated using NG method. **E-G**: the monthly total NT sequence variations along the evolution time for Orf1ab, S and Orf8. **H-R**: the Ka/Ks ratio using NG method (black closed circle) and 1-dose vaccinations (grey closed squares) for the major and accessory genes. See **Figure S11** for individual UTR and TRS.

**Figure 11.**
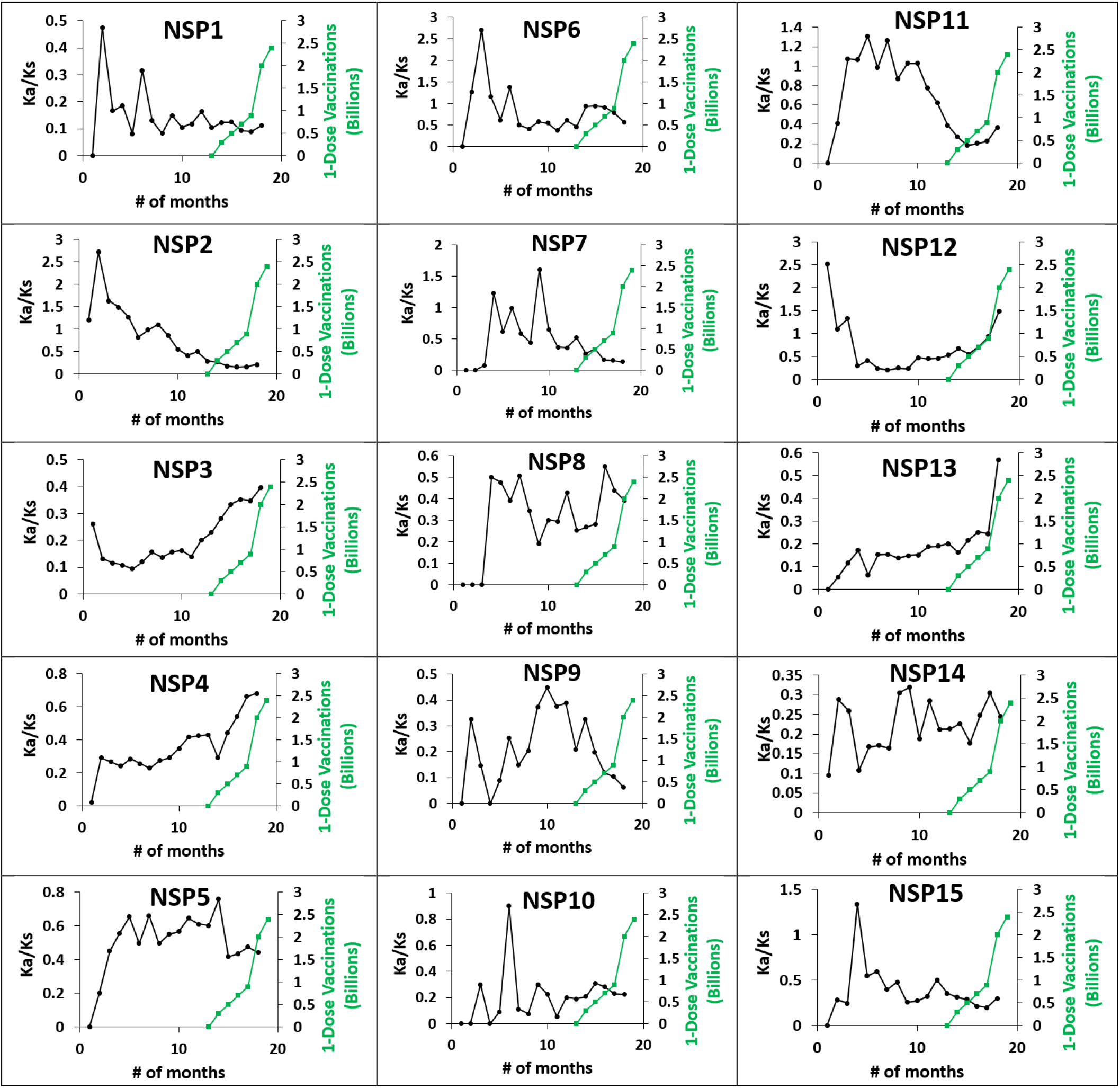
Timelines for Nsp 1-15 gene from the combined sets. Each cell: the Ka/Ks ratio using NG method (black closed circle) and 1-dose vaccinations (green closed squares).

We compared the percentage of the selection type from our *c/μ* test with that from Ka/Ks analysis for All-TR, each coding gene and Nsp1-15 (**Table S10**). Because the percentage of NT sites undergoing negative or positive selection were consistent across the four Ka/Ks methods (**Table S10A and S10B**), Ka/Ks was calculated for each NT site using the NG method. Encouragingly, results from the *c/μ* test and Ka/Ks analysis were generally consistent with each other for each coding gene, however *c/μ* slightly overestimated negative selection and slightly underestimated positive selection for some genes (**Figure 8**). This again confirms our theoretical proof that our *c/μ* classification scheme is consistent with the conventional Ka/Ks scheme under near-neutral selection.

### Time-independent GSR and Ka/Ks~0.7 observed in genome and All-TR, suggesting near neutral selection type, but their independence from the census population is consistent with NNBST rather than ONNT

Up until now, the substitution rates and selection type assignment for the compiled SARS-CoV-2 sequences were determined without explicit temporal information. However, temporal information is critical for elucidating the genomic evolution of SARS-CoV-2. In fact, a significant increase in the total NT sequence variation was observed in the omicron variants as opposed to earlier variants (i.e., beta, delta) (**Figure 1C**). Consistent with the total NT mutation timeline from the Nextstrain team between Dec. 2019-July 2021 (**Figure 1C**), the total NT sequence variation for the SARS-CoV-2 genome exhibited a very good linear correlation coefficient against time (*R*^2^ = 0.9833, **Figure 10A**) leading to a time-independent substitution rate (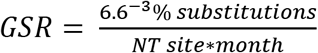 *or* 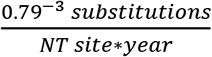) that was generally consistent with the other studies (**Table S2**) [42, 44–46, 80, 81], especially that reported by Callaway et al. (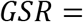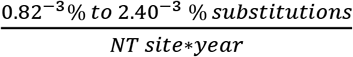). This time-independent GSR is obviously independent from the observed census virus population (10^0^ to 10^7^ infection cases), which varied significantly in the latter ten months (*R*^2^ = 0.0669) compared to the first nine months (*R*^2^ = 0.9624). This phenomenon is analogous to the zero-order kinetics of chemical reactions, in which the reaction rate is constant and does not dependent on the reactant concentration. This suggests that SARS-CoV-2 does not follow ONNT; If it did, we would expect that because the effective population would increase at least several magnitudes as the census population fluctuated, thus leading to a decrease in the substitution rate.

For All-TR and each coding gene, however, its correlation coefficient (*R*^2^ = 0.9805) and calculated total NT substitution rate (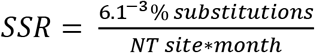) suggests it undergoes near-neutral selection types (**Figure 10B**). Its total NT sequence variation was decomposed into non-synonymous sequence variation (**Figure 10C**) and synonymous sequence variation (**Figure 10D**). Both timelines exhibited very linear correlation coefficients (*R*_*Ka*_ = 0.9522 *and R*_*Ks*_ = 0.9417) and their substitution rates were also time-independent constants (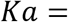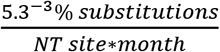 *and* 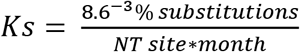), leading to a time-independent Ka/Ks of ~0.7.

Alternatively, the monthly non-synonymous and synonymous NT sequence variation of All-TR can be used to calculate out its monthly Ka/Ks timeline (**Figure 10K**), which varied around ~0.6. These time-independent rates and the ratio of ~1 are thus consistent with NNBST.

### Time-dependent SSRs suggest non-neutral selections, consistent with NNBST

The monthly percent total NT variation for some genes (Orf1ab, S and Orf8, **Figure 10E-G**) was also calculated and their correlation plots demonstrate that the near-constant GSR and SSR for the genome and All-TR, respectively, are not quite followed in each gene. While the Orf1ab-gene showed quite linear SSR, the S-gene and Orf8-gene clearly show fluctuations in their overall SSRs. Intriguingly, the fluctuations reflect an increase in SSR after the 10^th^ month, which is at the same time as the SARS-CoV-2 vaccine distribution. To assess the time-dependent features of translated proteins, the non-synonymous protein substitution rates for S, E, M and Orf8 were calculated over 19 months as examples and their correlation distributions plots were compared to the total NT substitution rate correlation plots in **Figure 10** (data not shown). Indeed, the two plots were nearly identical for S, E and Orf8, indicating the time-dependent features are also exhibited in the translated proteins. The two correlation plots were different for M, which showed a relatively constant protein substitution rate until a spike in the rate occurred in the 19^th^ month. This could also be represented as a time-dependent feature.

### Time-dependent Ka/Ks observed in most coding genes suggests non-neutral selection types at gene level, and thus supporting NNBST rather than KNT and ONNT

The time-independent Ka/Ks for All-TR does not guarantee time-independent Ka/Ks for each gene. The monthly Ka/Ks using the NG method was calculated for each coding segment over the combined three datasets (**Figures 10-11**). Intriguingly, the major genes, accessory genes and Nsp1-15 exhibited increasing or decreasing time-dependent (non-linear) Ka/Ks and did not follow the GSRM. For the major genes, Orf1ab had the least time-dependence, whereas N, S, M and E had increasing time-dependence. For the accessory genes, Orf3a had the least time-dependence, whereas Orf8, Orf7a, Orf10 and Orf6 had increasing time-dependence. For Nsp1-15, Nsp1,3-5,8-10,13-14 had the least time-dependence, whereas Nsp2,6-7,11-12,15 had increasing time-dependence. These results suggest stronger selection types are likely responsible for the variations in Ka/Ks of genes, in which time-dependent substitution rates of genes under non-neutral selection (positive and negative) were balanced out to generate time-independent GSR, following NNBST.

### Positive correlation observed between vaccinations and Ka/Ks in All-TR over time

We observed an increasing Ka/Ks as early as the 11^th^ month in the pandemic (November 2020) for All-TR, Orf1ab, N, S, M, E, Orf7a, Orf8, Nsp3, Nsp4, Nsp12 and Nsp13. Intriguingly, we also noticed an increase in 1-dose global vaccinations after December 2020 (**Figure 1B**). We were hence motivated to determine if a positive correlation existed between 1-dose vaccinations and Ka/Ks for All-TR and each coding gene. If so, this could help explain that the vaccine might drive the emergence of SARS-CoV-2 mutants. Intriguingly, good correlation was observed between increasing Ka/Ks values and increasing 1-dose vaccinations for All-TR and some genes (**Figures 10-11**). Correlation plots modeling 1-dose vaccinations against NG Ka/Ks were generated (**Figure S15**) and regression parameters (correlation coefficient and slope) were tabulated (**Table 3**). We chose 1-dose vaccinations as this is generally sufficient to induce an immune response. We specify a correlation coefficient cutoff (R^2^>0.6000) for segments showing good correlation between increasing 1-dose vaccinations and increasing Ka/Ks. Indeed, All-TR (0.9012) showed good correlation, as did most major genes, all accessory genes and some Nsps. For the major genes, M (0.8780), N (0.7959) and S (0.7435) showed good correlation, whereas Orf1ab (0.328) and E (0.029) showed low correlation. For the accessory genes, Orf8 (0.8339), Orf3a (0.7311), Orf7a (0.687), Orf6 (0.6638) and Orf10 (0.6539) all showed good correlation. For the Nsps, Nsp3-4,9,12-13 (0.6322 - 0.8642) showed good correlation, whereas Nsp1-2,5-8,10-11,14-15 (0.0014 – 0.5845) showed low correlation. This suggests that M, N, S, Orf3a, Orf6, Orf7a, Orf8 and Orf10 genes may play some role in combating the immune response and serve as potential drug targets in reducing COVID-19 infection.

### Positive correlation observed between vaccinations and total NT variation in UTR/TRS over time

Correlation plots modeling increasing 1-dose vaccinations and percent total NT variation of All-UTR, All-TRS and each UTR/TRS were also generated (**Figure 10 and S11**) to investigate if the vaccine increased the substitution rate of the non-coding regions. The slope and correlation coefficient values have been tabulated (**Table 4**). Surprisingly, good correlation was observed for All-UTR, some UTR and TRS. From R^2^, M 5’UTR (0.9038), All-UTR (0.7230), Orf1ab 5’UTR (0.6885) and Orf8 5’UTR (0.6359) showed good correlation, whereas Orf10 5’UTR (0.4600), Orf7a (0.4226), N 5’UTR (0.4103), Orf10 3’UTR (0.3962), S 5’UTR (0.2967), Orf6a 5’UTR (0.0233), E 5’UTR (0.0092) and Orf3a (0) showed low or no correlation. For TRS, Orf6 TRS-B (0.8942) and N TRS-B (0.7415) showed good correlation, whereas Orf1ab TRS-L (0.4543), Orf7a TRS-B (0.3856), Orf8 TRS-B (0.2908), S TRS-B (0.1450), All-TRS (0.0607), E TRS-B (0), M TRS-B (0) and Orf3a TRS-B (0) showed low or no correlation. The good correlation observed in Orf1ab 5’UTR is worth noting, as it showed a high relative NT substitution rate (**Table 4**, Rel^1^ = 5.3) compared to the genomic substitution rate. This may also warrant it as a potential drug target.

### Top NT/AA sites under strong positive selection identified in UTR and proteome using both *c/μ* and Ka/Ks

To determine regions of critical functionality, we identified the top non-synonymous NT and AA sites undergoing strong positive selection in the UTR and All-TR of the proteome. Identifying such sites could be potential drug targets in treating COVID-19. NT sites were sorted by decreasing *c/μ* values with cutoffs (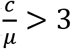 *with c* > 30) in the supporting Excel sheets. Conserved sites were designated as (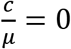). A total of 54 NT sites from and Orf10 3’UTR (23), Orf1ab 5’UTR (14), Orf8 5’UTR (6), Orf10 5’UTR (4), N 5’UTR (3), M 5’UTR (2), Orf7a 5’UTR (2) exhibited strong positive selection. S 5’UTR, E 5’UTR, Orf3a 5’UTR and Orf6 5’UTR did not have any sites under strong positive selection (see supporting Excel book file). The top NT sites were mapped to secondary structure diagrams for each UTR to determine the impact on their functionality (**Figure S9**). Sites under strong positive selection seemed to occur in NTs that may be involved in intra-UTR base-pairing, such as in Orf1ab 5’UTR (first four NTs at the 5’ terminus, SL3, SL4 and SL5), Orf8 5’UTR, Orf10 5’UTR and Orf10 3’UTR (BSL and S2M). The location of these NT substitution sites may enhance UTR structural stability, improving viral replication or aiding in escaping recognition of host immune response elements (i.e., host miRNAs).

The top AA sites under strong positive selection in All-TR for the proteome and each protein were sorted by *c/μ* and Ka/Ks (SP method) with cutoffs (See supporting Excel book file). The identified top 247 AA sites in the proteome under positive selection cutoffs were manually set (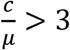 *and* 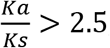, *with c* > 30 *and Ka* > 30). The top AA sites were mapped to protein structure plots for visualization (**Figure 9** for S, N, Orf8 proteins; **Figure S10** for all proteins). Briefly, **t**he S protein contained most sites under strong positive selection (63), followed by Nsp3 (31), N (31), Orf3a (18), Nsp12 (15), Nsp6 (12), Orf8 (12), Nsp4 (11), Nsp2 (10), Nsp11 (7), Orf7a (6), Nsp13 (5), Nsp5 (4), Nsp8 (3), Nsp14 (3), Nsp15 (3), M (3), Orf10 (3), Nsp7 (2), E (2), Nsp9 (1) and Orf6 (1). Nsp10 did not contain any sites undergoing strong positive selection. Secondary structure type was assigned to the top 247 AA substitutions in the proteome and the percentage of these substitutions occurring on four secondary structure types (alpha helix, beta sheet, beta turn and random coil) were calculated out (**Table S11**). For All-TR, the secondary structure composition is 31% alpha helix, 19% beta sheet, 27% beta turn and 23% random coil. Interestingly, we observed a similar percent substitution on beta sheets (18%), but markedly lower value on alpha helices (22%) and higher values on beta turns (47%) and on random coils (32%) for the top 247 mutations. There appears to be more positive substitutions at beta turns and random coils than at ordered alpha helical and beta sheet regions that increases the fitness of this virus.

## Discussion

A rigorous assessment of the five theories (ST, GSRM, KNT, ONNT and NNBST) for SARS-CoV-2 at different levels (i.e., NT/AA, gene/segment and genomic level) was conducted in the results section. Regardless of the large fluctuating factors (i.e., infected human population and vaccination numbers in **Figure 1**), so far the SARS-CoV-2 genome exhibited a near-constant substitution rate from linear fitting (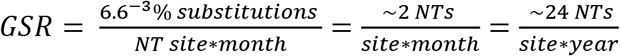) (**Figure 10**), falling within the typical bounds of genomic substitution rates of other RNA viruses (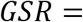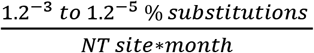) [82]. In fact, a subset of RNA viruses (i.e., HIV-1) seems to assume time-independent GSR [83]. The fact that the GSR of SARS-CoV-2 showed a lack of dependence on the effective virus population size that should be proportional to the infected human population that fluctuates between 10^0^ to ~10^7^, suggesting ONNT is not applicable for this virus. Although the time-independent GSR is strongly indicative of the molecular clock feature, which is a hallmark of the three neutral theories (GSRM, KNT or ONNT), the Poisson distribution with a single constant substitution rate as implied by the GSRM and KNT was not observed at NT/codon level (**Figure 3**), segment level (**Figure 6 and 7**) or the genome level (**Figure 4**). Instead, we observed an L-shaped probability distribution for *c/μ* of the genome and each gene (**Figure 4**), supporting NNBST in which the lower substitution rate of sites under negative selection (*c/μ*<1) is balanced with the higher substitution rate of sites under positive selection (*c/μ*>1), leading to a constant substitution rate that is close to the mutation rate (*μ*). Our NNBST is also very different to the ONNT in which nearly neutral sites are slightly deleterious and thus their substitution rate is inversely proportional to the effective population size. In contrast, our NNBST is unlikely modulated by the effective population size due to the opposite effects of the effective population size on the substitution rate between weakly negative sites and weakly positive sites: the small effective population size speeds up the substitution rate of the weakly negative sites but slows down the substitution rate of the weakly positive sites. In contrast, the large effective population size slows down the substitution rate of the weakly negative sites but speeds up the substitution of the weakly positive sites. Interestingly, this appears to support the data that the constant substitution rate did not depend on the census population change (**Figure 1**). Although many researchers think that KNT is unlikely true as more and more genomic data become available, invalidation of KNT appears to be impossible because Kimura’s original statement of KNT was qualitative rather quantitative. Our relative substitution rate analysis provides a straightforward tool to use the genomic data to assess if KNT is true or not. Many researchers also believe that ONNT, considered a refinement of KNT in that it has addressed many of its weaknesses, could be true. Thus, it is striking that our analysis has showed ONNT is not true for this virus.

While the relative abundance of different selection types in a genome have been qualitatively characterized for ST, KNT, and ONNT [4] (**Figure 12A-C**), the percentage of genomic sites under these selection types have not been quantified to determine which theory is true for this virus. To the best of our knowledge, our *c/μ* tool allowed us to quantify the percentage of NT sites in the genome including UTR under different selection types for the very first time. The pie charts (**Figure 5**) revealed that while most NT sites are conserved or undergo strong negative selection, near-neutral selection (both slightly deleterious/advantageous) and strong positive selection were also observed throughout the genome, thus SSRM results confirmed that SARS-CoV-2 followed NNBST rather than ST, KNT and ONNT. NNBST differs ST in two folds: 1). While NNBST allows some NT sites under weakly negative and weakly positive selection, ST does not allow many such NT sites. 2). the NT sites under negative selection is balanced with the sites under positive selection (**Figure 5**). For example, the GSR of this virus appears to a time-independent constant leading to a linear plot of sequence variation against time, suggesting effective “neutral” evolution. However, in fact some of genes are actually under (strong) negative selection and some are actually under (strong) positive selection, all showing non-linear curves of sequence variation leading to time-dependent SSR. When averaging over all them, the lower SSR from the genes under the negative selection is balanced out with the higher SSR of the genes under positive selection, leading to a time-independent SSR (**Figure 13**). To the best of our knowledge, for the very first time, we showed that NNBST can be consistent with the molecular clock feature, which was considered as the hallmark of the neutral theories only so far.

**Figure 12.**
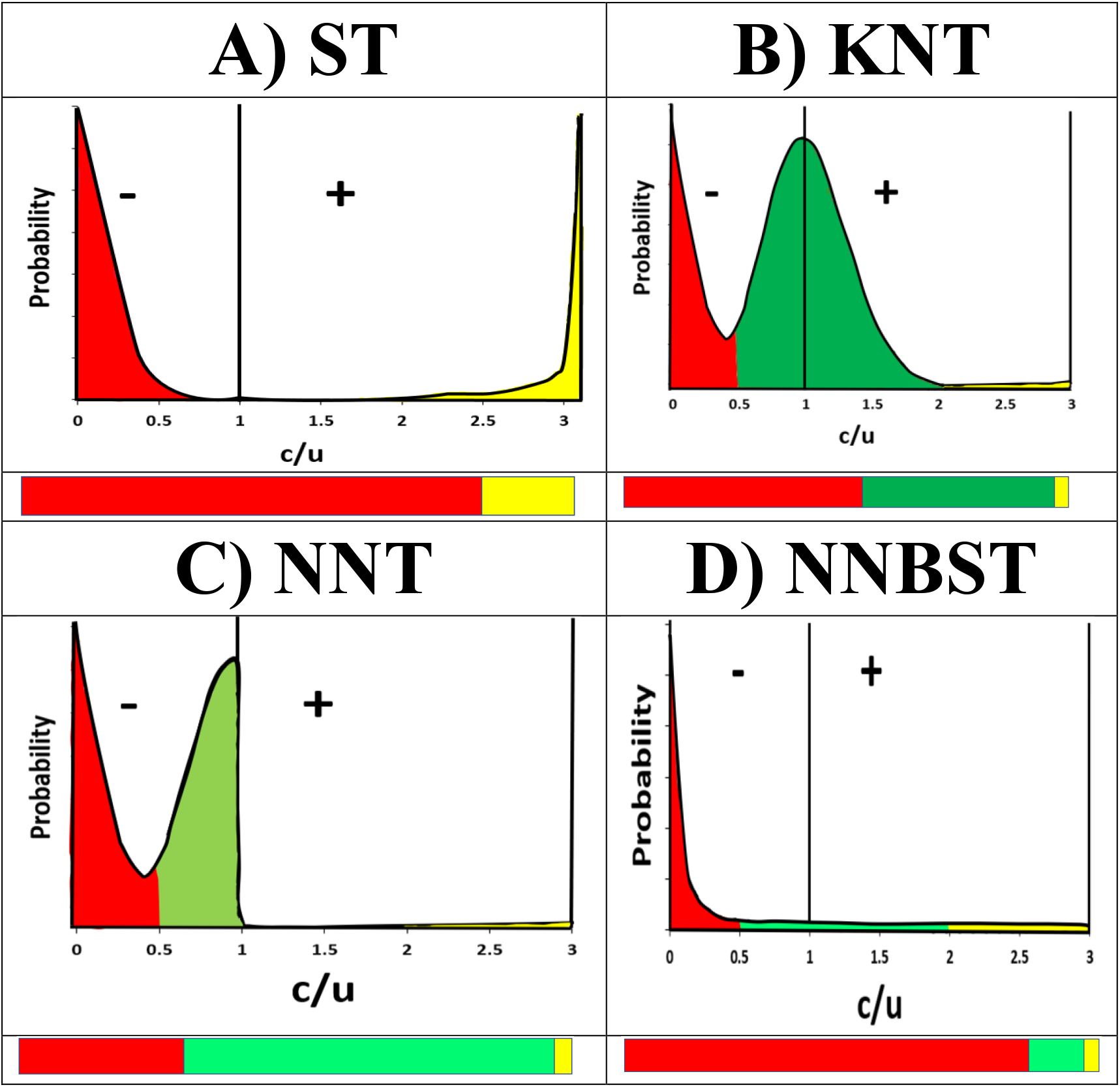
Diagrams of the four evolution theories **A**: ST; **B**: Poisson probability distribution implied by KNT/GSRM; **C**: Asymmetric distribution of the nearly neutral site implied by ONNT. **D**: L-Distribution theory implied by the NNBST. **Top**: Probability distribution showing negative (−), positive (+) and neutral selection (straight line). **Bottom**: Selection type distributions for the four theories of molecular evolution. Colors are assigned for strong negative/conserved selection (red), neutral selection (dark green), weak positive/weak negative selection (light green) and strong positive selection (yellow). Regions enclosed in the dashed lines in **C)** indicate weak positive selection (left) and weak negative selection (right).

**Figure 13.**
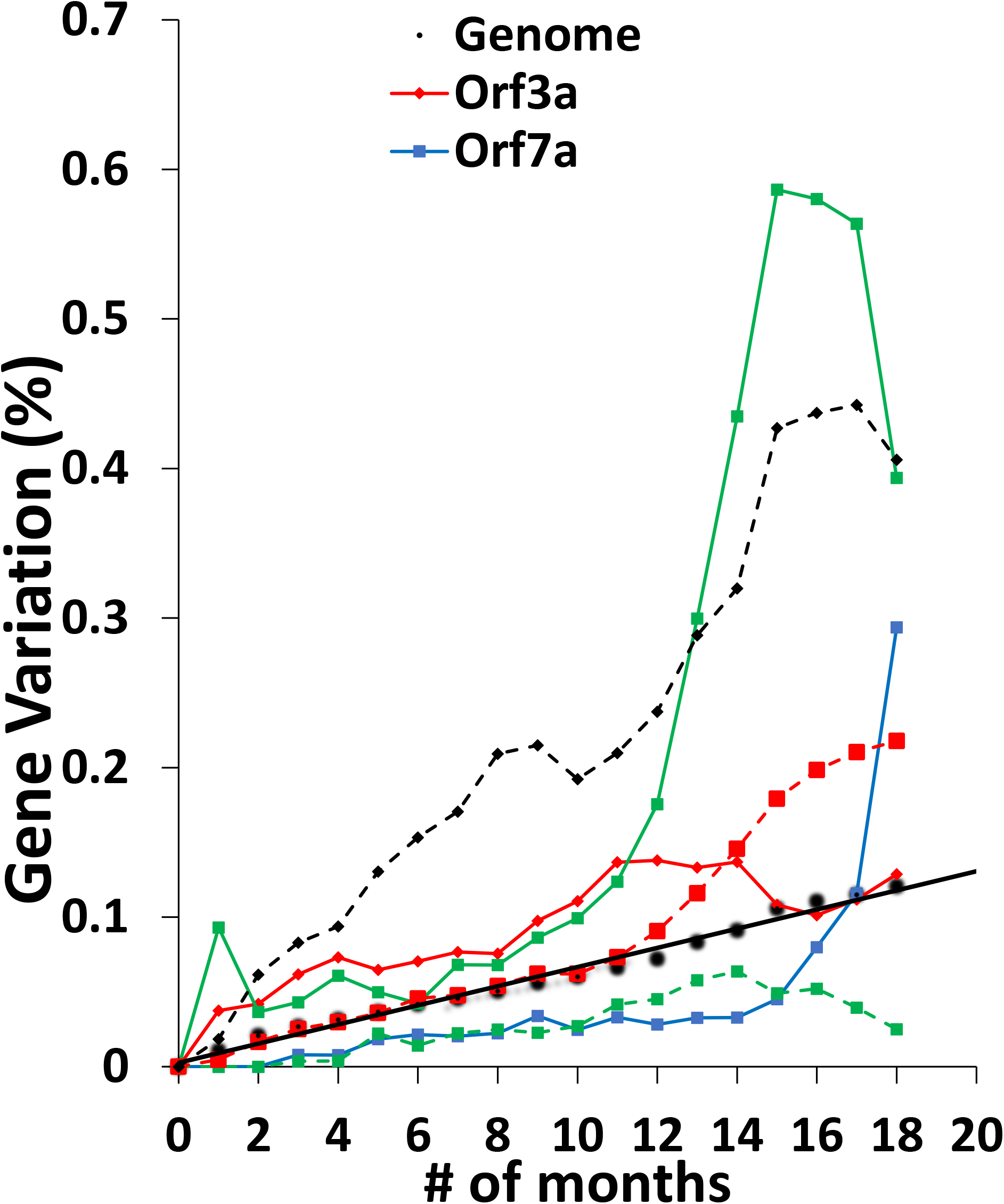
Monthly time-independent substitution rates of the genome (solid black) and time-dependent features of several major genes (E: dotted green; S: dotted red; N: dotted black) and accessory genes (Orf3a: solid red; Orf7a: solid blue; Orf8: solid green).

In the same logic, our results also suggest that the molecular clock feature of a gene or protein is not sufficient to support KNT or ONNT [5, 6] on the gene or protein, the actual mechanism could be NNBST at protein domain level. Because while some domains or motifs of the gene and the protein, small self-folding regions (between 50 to 200 AAs in length) containing various secondary structure elements (i.e., α-helix, β-sheet, β-turn, random coil), could be under negative selections showing time-dependent lower substitution rate, some of them could be under positive selection showing time-dependent higher substitution rate, but the averaging effect of all domains or motifs under both selection types could show time-independent substitution rate (the molecular clock feature) that seems to be caused by neutral selection. Therefore, a more detailed analysis on each domain or motif of a gene or a protein is required to decipher the exact relative role of genetic drift, environmental selection and other factors in the evolution of the gene or the protein. Gojobori, Moriyama and Kimura reviewed synonymous (Ks) and non-synonymous (Ka) substitution rates of viral oncogenes from HIV-1, HBV and influenza A viruses to determine if they followed the KNT (e.g., Ka<Ks exhibits a molecular clock feature). They revealed that the synonymous substitution rates were significantly higher than the non-synonymous substitution rates for *gag* oncogene in HIV-1 (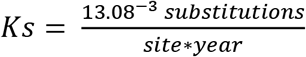 *and* 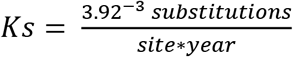), *P* oncogene in HBV (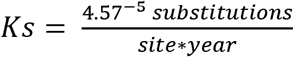 *and* 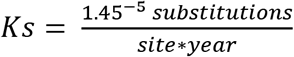) and *Hemagglutinin* oncogene in influenza A (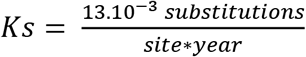 *and* 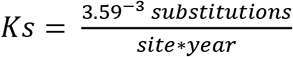) [5]. A molecular clock feature was also described for *Hemagglutinin* in influenza A virus, suggesting these viruses nicely follow KNT. Indeed, recent literature reports explain KNT is followed in other viruses, including those in the human virome [84], siphoviridae phage [85] and human cytomegalovirus [86]. However, these three viruses may exhibit balancing negative and positive selection that may explain their molecular clock feature. Therefore, a re-examination of the molecular evolution of these and other RNA viruses should be conducted to determine if they truly follow KNT as previously proposed. Additionally, the substitution rates of proteins from various organisms (e.g., fibrinopeptides, cytochrome c and hemogloblin) [87] exhibiting molecular clock features were also said to follow KNT. However, are balanced selections actually behind these molecular clock? The molecular evolution of these proteins should also be re-examined to domain and motif level to see if they actually follow the NNBST! If true, NNBST could enable our better understanding of molecular evolution and biology.

Like the GSR of the genome, the SSR of the S-gene showed nearly-neutral features **(Figure 10F**). However, the individual protein domains of the S-protein (S1: N-terminal domain (NTD), receptor-binding domain (RBD), furin cleavage site and S1/S2 cleavage region; S2: Heptad repeat 1, transmembrane domain (TMD) and carboxy-terminal domain (CTD)) may also exhibit either higher substitution rates under positive selection or lower substitution rate under negative selection and thus show time-dependent features (**Figure 4H**). A spectrum of the Ka/Ks across each AA residue was generated and high substitution sites indicative of strong positive selection (Ka/Ks > 2.0) were mapped to the S protein monomer structure model (PDB: 6VSB) for ease of visualization. (**Figure 14)**. Indeed, 18 peaks corresponding to strong positive selection were identified across the protein (S1: 12; S2: 6). In S1, the NTD, RBD and the rest of S1 exhibited four peaks each, with the RBD (e.g., A520S) and the latter portion of S1 (e.g., N658D) exhibiting the highest Ka/Ks substitution sites. Because RBD enables its binding to the host ACE2 cell surface receptor, mediating cell entry, it was targeted by antibody and vaccine therapies can neutralize and prevent cell infection [88–90]. The mutations in the RBD in response to these therapies has enabled antibody resistance and vaccine escape, as well as enhanced binding affinity to host ACE2 [91, 92]. A520S might be such mutation under this positive selection. While not mapped, high substitution sites were observed in the furin cleavage sites and S1/S2 cleavage regions, which might explain the increased infectivity and transmission profiles of later SARS-CoV-2 variants due to vaccine and host immune response effects [93]. In S2, high peaks in heptad repeat 1 (D985D and A1025G) and near the CTD (D1118H, S1161S and V1177L) were also observed. Conversely, the CTD and TMD showed no peaks, indicating they are likely functionally important. These observations clearly suggest that the higher/lower time-dependent SSRs of different domains balance out to produce the overall time-independent SSR of the S-protein, which is consistent with NNBST. Additionally, domains exhibiting strong positive selection may be potential drug targets. In the future, detailed analysis of domains in other SARS-CoV-2 proteins will be analyzed.

**Figure 14.**
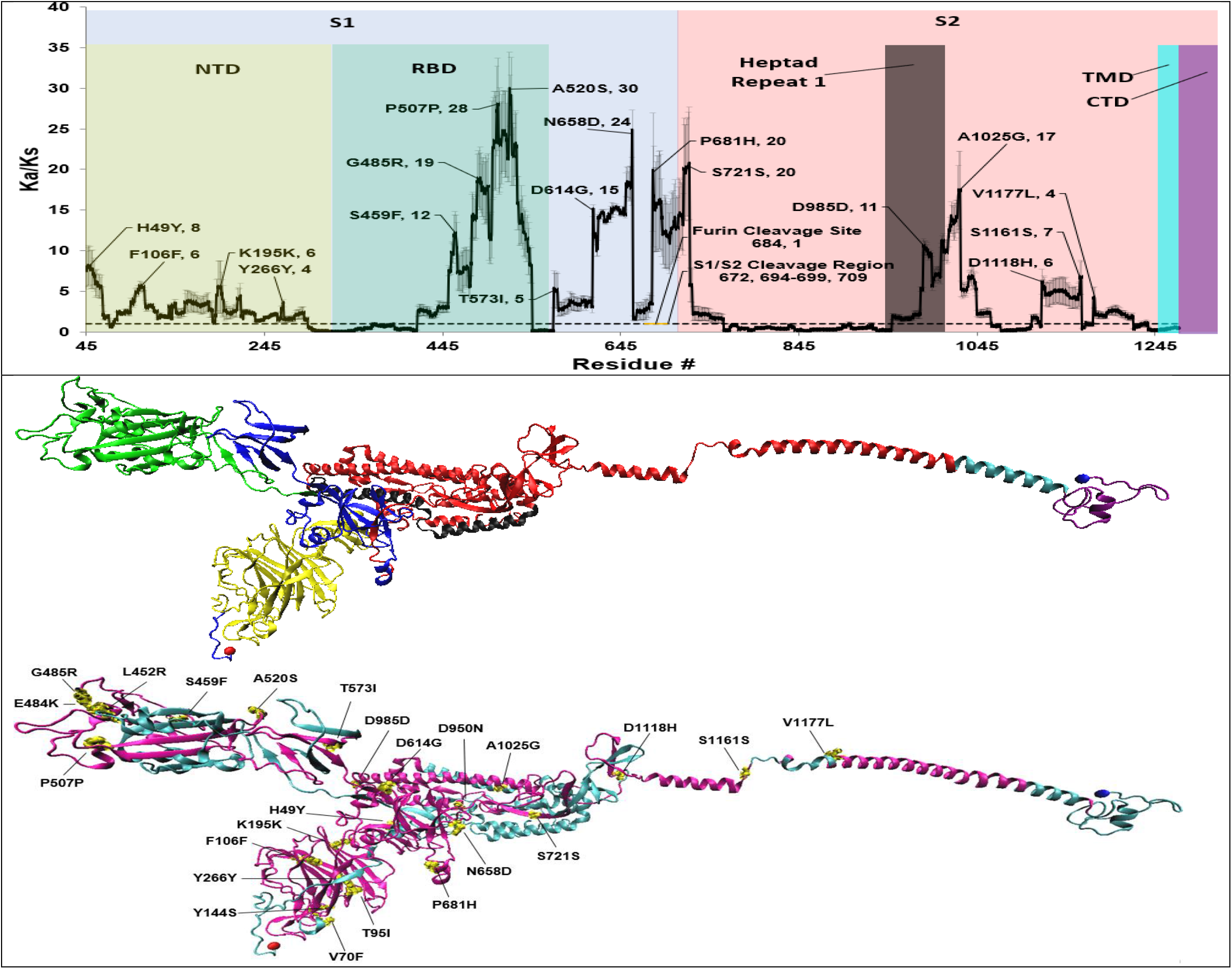
Average Ka/Ks ratio calculated over the S protein sequences using NG method, starting from the 45^th^ AA residue. (Ka/Ks plot). The structure is color-coded to highlight different S protein domains (S1: NTD and RBD; S1/S2 cleavage region: Furin cleavage site; S2: Heptad Repeat 1). (Protein structure models) S protein structure in the open-state (i.e., ready for binding hACE2 receptor), with the viral receptor bonding motif (RBD) facing front (**PDB ID**: 6VSB). The first structure model is color-coded by domain and the second model is color-coded by Ka/Ks<1 (blue) and Ka/Ks>1 (purple). Ka/Ks peaks corresponding to an AA are represented as yellow bonds. N- and C-termini are colored in red and blue spheres, respectively.

In the field of molecular phylogenetics, relaxing the assumption that all NT sites have the same substitution rate in GSRM originated from two prior assumptions: A prior non-uniform distribution of rates such as a gamma distribution described by Nei et al (1976) or a lognormal distribution by Olsen [94] (1987). To reduce the computation cost, Yang [95] (1994) used a discrete approximation, replacing a gamma distribution of rates by a discrete distribution with four well-chosen classes. Similarly, Felsenstein et al [63] (1996) used Hidden Markov Model (HMM) methods with a modest number of discrete rates to model unequal and unknown evolutionary rates at different sites in molecular sequences. On one hand, our observed L-shaped distribution supports the use of a non-uniform distribution. On the other hand, please note that the L-shaped distribution could not have been fitted using a Gamma distribution curve, which is unable to handle cases where x = 0 (i.e., *c/μ* = 0 conserved sites under extreme negative selection).

In genetics, Ka/Ks analysis is commonly used to estimate the selection type for each coding gene in TR [10–16]. The concept of Ka/Ks arose from the McDonald-Krietman test, which sought to determine the ratio (Ka/Ks) of fixed non-synonymous substitution rate (Ka) to fixed synonymous substitution rate (Ks) in protein-coding gene regions with reference to the ratio (P_n_/P_s_) of non-synonymous polymorphisms (P_n_) to synonymous polymorphisms (P_s_) in protein-coding gene regions [96]. Because P_n_/P_s_ ≈ 1, its selection type assignment is like Ka/Ks for positive selection (Ka/Ks>1), neutral selection (Ka/Ks≈1) and negative selection (Ka/Ks<1) [97]. However, Ka/Ks is unable to probe the non-coding UTR and TRS regions. In this study, we formulated a *c/μ* analysis and applied it to systematically analyze the selection type of not only 10 genes, but also 11 UTRs/9 TRSs. In the methods section, we further provided a theoretical proof to show our *c/μ* classification scheme for a gene in TR is consistent with the conventional Ka/Ks scheme under near-neutral selection for the whole genome. The two critical conditions leading to the consistency proof are **1**). K_a_ = K_s_ that the non-synonymous mutation rate (Ka) is approximately equal to the synonymous mutation rate (Ks) for All-TR under nearly neutral selection. **2**). K_s_^i^ = K_s_ under near-neutral conditions that the synonymous mutation rate for each coding gene (K_s_^i^) is approximately equal to the synonymous mutation rate for All-TR (K_s_). Indeed, the Ka/Ks^b^ is ~0.8 for All-TR, thus the first condition was met. For the second condition, we calculated the relative synonymous mutation rate of each coding gene to the synonymous mutation rate of All-TR (**Table S7**). Indeed, all the coding genes except N (3.2) have a value that is close to 1, thus the second condition was also met. Encouragingly, selection type assignment by *c/μ* was mostly consistent with that by Ka/Ks for the 10 coding genes (**Figure 6**). Furthermore, percent NT/codon sites undergoing different selection types for All-TR are also consistent between *c/μ* and Ka/Ks.

Given the success of *c/μ* classification in the TR, we are confident that our *c/μ* classification for the UTRs and TRSs are reliable: All-UTR may undergo positive selection; All-TRS likely undergoes strong negative selection. Furthermore, more UTRs from accessory genes than UTRs from major genes likely undergo positive selection. SSRs of TRS from accessory genes were significantly higher than TRS from major genes, suggesting more critical functional roles of the major genes than the accessory genes. This finding is also consistent with the lower level of conservation in accessory genes versus major genes across different betacoronaviruses, as changes in TRS could prevent translation of their associated genes. This may be the case for Orf10 (more will be discussed later).

UTRs which exhibited negative selection might be explained by their roles in regulating the discontinuous transcription of sgRNAs. Indeed, altering a UTR sequence might prevent its recognition by RdRp, thus its downstream sequence may not get translated into protein. Consequently, UTRs which exhibited positive selection might be to escape recognition by host cell microRNAs (miRNAs) [98, 99]. miRNAs are small (~22 NT), non-coding ssRNAs which bind to complementary RNA sequences within the leader sequence of 5’UTR of corresponding sgRNAs and 3’UTR and effectively inhibit viral replication [98]. Thus, the 5’UTR and their leader sequences might exhibit non-synonymous substitutions sourced from slight positive selection to evade recognition by host miRNAs. Additionally, the Orf1ab 5’-UTR and the 3’-UTR contain cis-acting secondary RNA structures essential for RNA synthesis [100, 101]. In Orf1ab 5’UTR, five stem loop secondary structures (SL1, SL2, SL3-TRS-L, SL4 and SL5) are present. Maintaining the conformation of SL1 ensures efficient viral replication and escape from Nsp1-mediated translation suppression, which downregulates host cell mRNA translation. SL2 is essential for viral viability. SL3 contains the TRS-L and is thus required for sgRNA transcription. SL4 directs sgRNA synthesis during viral synthesis. SL5 helps package viral RNA and aids in translation of Orf1ab [22, 102, 103]. Thus, maintaining the structures of the five stem loops is critical for ensuring efficient viral replication.

It’s not surprising that All-TRS likely undergoes strong negative selection. TRS conservation is critical for correct synthesis of sgRNAs, such that substitutions in TRS could lead to genes not being transcribed and translated. Orf10 does not have a TRS-B and its associated sgRNA and protein have not been observed [23, 104]. Hypothetically, if a conserved TRS-B could be generated for Orf10, transcription and translation of Orf10 sgRNA should be observed. We aligned hypothetical Orf10 TRS-B sequences against the conserved TRS-B sequence across each gene; we assumed the hypothetical Orf10 TRS-B sequence is located within the last seven NTs in the Orf10 5’UTR based on the positions of the known TRS. We observed at least three non-synonymous NT substitutions in the hypothetical Orf10 TRS-B sequences compared with the conserved TRS-B sequence (**Figure 7**). It would be interesting to know if inducing single-point mutations in hypothetical Orf10 TRS-B to match the conserved TRS-B would generate a functional Orf10 protein, demystifying the current belief that Orf10 is not critical for maintaining viral transmission or replication [105].

Mutation analysis of 8 SARS-CoV-2 proteins (S, E, M, N, Nsp3, Nsp5, Nsp11 and Nsp14) was performed by Wang et al. [36], where they identified 8,309 substitution mutations in 15,140 genomic samples (January 5, 2020-June 1, 2020). They listed the top 10 AA substitutions for these proteins ranked by their total mutation frequency, including both synonymous and non-synonymous substitutions. Interestingly, 11 non-synonymous AA sites in their analysis are also in our top 247 AA list (see data file, highlighted in red). A few things to note: 1). Their analysis included a genomic dataset relatively early in the pandemic (6 months), such that the virus did not undergo extensive change compared to our dataset (19 months); 2). Their analysis included a total of 8 proteins (each major protein and 3 out of 15 Nsp proteins) but did not include the accessory proteins, each UTR or TRS; we investigated all 25 coding proteins (each major and accessory protein and each Nsp) as well as the 11 UTRs and 9 TRSs in our study due to the increasing relevance the accessory proteins and non-coding regions likely play in the virus’s viability and infectivity; 3). Their top 10 mutations are based on total substitution count and includes both non-synonymous and synonymous substitutions, which varies between each analyzed protein. For instance, most synonymous substitutions were observed in the M protein (8 out of 10), followed by E (7 out of 10), S (5 out of 10), Nsp11 (4 out of 10), Nsp14 (4 out of 10), Nsp5 (3 out of 10), Nsp3 (3 out of 10) and N (2 out of 10). In contrast, our study uses a *c/μ* and Ka/Ks cutoff to identify AA and NT sites exhibiting strong positive selection, which are likely key functional regions of importance (i.e., vaccine escape, adaptivity). The top 54 NT sites and top 200 AA sites which exhibited strong positive selection in All-UTR and All-TR could serve as good drug targets in stopping COVID-19 infection. Mapping the top sites to secondary structures of UTR or 3D-protein structures can provide spatial insight that predict their functional impact. For instance, NT mutations in UTR (i.e., Orf1ab 5’UTR) may enhance their stability and may allow for easier and faster viral replication processes. Similarly, AA mutations can also stabilize certain protein structural conformations (i.e., “opened” state of S protein) that enhance virus infectivity profiles.

We were hence motivated to determine if a positive correlation existed between 1-dose vaccinations and Ka/Ks for All-TR and each coding gene. The higher correlation coefficient observed in most accessory genes (Orf3a, Orf6, Orf7a and Orf8) and Nsp3 may not be surprising, as these genes likely contain immunoregulatory function (**Table 3**). The Orf3a protein forms tetrameric ion channels and facilitates virus release [106] [107]. The Orf6 protein inhibits the expression of interferon (IFN)-stimulated genes and stunts import trafficking of biomolecules into the nucleus [108]. The Orf7a protein antagonizes the IFN-I response to reduce antiviral activity [109]. The Orf8 protein also disrupts IFN-I signaling and downregulates major histocompatibility complex one (MHC-I) to evade the cellular immune response [110] [111]. The vaccine serves as a selection type against the virus. In response, the viral substitution rates of these genes are likely to increase to adapt and evade the host cell immune response, leading to the increasing Ka/Ks of these genes.

The c/μ framework might resolve a long-standing “neutralist-selectionist” controversy on how to interpret a high substitution rate for a gene or a NT site of a species [112]. For a gene or a NT site under zero or a low substitution rate, both the KNT and the ST converge on these genes to be “critical” for a species’ life cycle. For example, non-synonymous alterations to the NT sequence (i.e., UTR/TRS) or AA codon (i.e., Nsp1-15) can potentially disrupt the replication cycle of SARS-CoV-2 or produce non-functional proteins. However, for a gene and a NT site under high substitution rate, the predicted selection type on these substitutions is mutually exclusive using these two basic theories. The KNT interprets these substitutions as neutral selection type that doesn’t impact species’ fitness for survival, and thus the gene and the NT site are functionally less important. In contrast, the ST interprets these substitutions as both non-neutral and positive selection that are adaptive, and thus the gene and the NT site are functionally more important. For example, the SARS-CoV-2 S protein has undergone several mutations resulting from the mRNA vaccine treatment that is “responsive” to changing environmental conditions that threatens the species’ survival. Therefore, to resolve controversy, an objective test to decide whether new substitutions tend to neutral, nearly neutral, purifying or advantageous is required. Our c/μ framework just like Ka/Ks test provides such a test in which if *c/μ*>1 for a NT site or a gene, then it is assigned to positive selection type rather than neutral section type. Because *c/μ*, like Ka/Ks, can be determined from the experimental sequence data, thus the selection type assignment from them is operational and objective.

FGMR (*μ*) of the genomic replication machinery (RdRp with ExoN) is independent from the virus genome sequence change and thus the replication error rate does not change significantly over a short period of time. In-vitro assays or in-vivo cell lines [63] might be used in the future to experimentally determine FGMR. To understand this phenomenon, we proposed a hypothetical model for viral genomic replication and mutation under no selection type (i.e., *Nc = Ne* in **Figure 2**). It might be helpful to view this as an in-vitro experiment: A test tube containing a constant, high concentration supply of viral RNA genomic sequences and viral RdRp in nutrient media and without selection types (i.e., RNase, vaccine and antibody-free conditions; no cells to avoid death and host immune response). Multiple cycles of replications are conducted, the mutation rate without selection can be measured by sequencing. From this, we could obtain the experimental fundamental genomic mutation rate (*μ*) for any virus (i.e., SARS-CoV-2, Influenza A and others) to validate our calculated substitution rate (*c=μ*) from the virus census population. It’s critical to note that polymerase architecture and replication fidelity can vary significantly within and between lower organisms (i.e., viruses) and higher organisms (i.e., mammals), thus the FGMR can also change significantly. Amongst viruses, the mutation rates of DNA viruses per cell infection (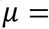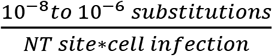) are much lower than that of RNA viruses (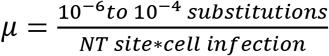), due to the higher error rate of RNA virus polymerases versus DNA polymerases [82]. SARS-CoV-2 RdRp, coronaviruses and DNA viral polymerases, typically contain proofreading mechanisms (3’-ExoN) and exhibit a lower replication error rate, however most RNA viral polymerases lack proofreading mechanisms and exhibit a higher replication error rate. It seems apparent that KNT and ONNT did not consider the effects of different polymerases on an organism’s FGMR, possibly due to a lack of biochemical and sequence data at their development. If the FGMR is assumed to be constant across different organisms, this will likely affect the calculation of relative substitution rates within the genome and thus assessment and quantification of their different selections. In knowing the FGMR for different organisms, the relative substitution rates at different sequence levels (i.e., NT/AA, segment) can be accurately probed.

Our NNBST could aid in resolving Lewontin’s Paradox on KNT, considered as a long-standing issue in evolutionary biology. Assuming the genetic variation balance between genetic drift (decrease of variation) and random mutation under the neutral selection only (increase of variation), the expected pairwise sequence diversity (Π) at neutral sites in a panmictic population of ***N_c_*** can be calculated via the equation:

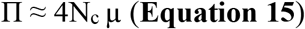

Where *μ* is the mutation rate per base pair per generation. It’s expected that Π should increase linearly with ***N_c_***. However, in 1974, Lewontin observed in metazoans that Π only increased two-order of magnitude, whereas Nc increased more than several-order magnitude [97]. Numerous efforts were made to resolve this paradox [98–100]. Recently, Buffalo predicted sequence divergence using **Equation 15** and also observed this paradox amongst different species and taxa (i.e., humans, metazoans, plants, etc.) and concluded that it is unlikely for linked selection, which strong and abundant selection are considered to explain this paradox. If the L-shaped distribution of relative mutation rate (*c/μ*) is general for all these species, then the KNT and the ONNT which has dominated the landscape of evolutionary theory for half a century will be proved to be false. If the KNT is false, then the foundation for Lewontin’s paradox on the KNT is removed, the paradox will be no longer exist. Our NNBST might be able to explain the lack of proportionality between pair-wise sequence variation π and ***N_c_*** as that the sequence variation is independent from the census population size observed in **Equation 15**.

Some limitations should be noted for our *c/μ* test and analysis. Concerning the methodology, we defined a viral genome sequence of finite length (*N*) which undergoes genomic substitution rate (GSR) without considering recombination events [113]. For SARS-CoV-2, when a mutant sequence becomes 100% divergent from the first virus sequence, we denote this term as the “Saturation point”. It is very likely for mutated NTs to be reverted to their initial NTs via point-substitution, decreasing sequence divergence. To avoid reaching saturation, the genomic sequence length has been previously defined as infinity (*N*∞) [114] although this is not a realistic parameter. Recombination as a rare event might be another mechanism to increase sequence variation (e.g., long insertion/deletion) in the genome, increasing the complexity of molecular evolution. Viral recombination occurs between two virus species within the same host cell, producing a new viral hybrid. Although we did not observe recombination events in our datasets and the chance for recombination is low, recently, SARS-CoV-2 strains containing chimeric S proteins Delta (AY.119.2) and Omicron (BA.1.1) were reported (December 31, 2021 to February 12, 2022) [115]. It would be interesting to investigate recombined viral sequences using our *c/μ* test.

In terms of analysis, despite our datasets spanning 19 months, this is considered a very short evolutionary time. As can be seen, some segments of SARS-CoV-2 (i.e., S-gene and S-protein) exhibited a rapidly increasing SSR 19 months after the first reported human infection, suggesting the virus has not fully adapted to the global human population; considering the immense genetic diversity of humans, significant time must likely pass before SARS-CoV-2 demonstrates adaption. Following adaption, we might expect a noticeably decreasing GSR and SSR for the genome and each gene segment. However, it was not observed in this study.

Lastly, if the identified top 54 NT and 247 AA codon substitutions in the UTR/TR under strong positive selection truly improve SARS-CoV-2 viral fitness, biological data is critical in supporting this claim. While we earlier inferred the effects of these substitutions on viral fitness from the current literature, their exact effect remains to be elusive. Addressing the first and second concerns, our *c/μ* method should be applied to the sequence dataset of SARS-CoV-2’s ancestors spanning hundreds of years in evolutionary time, which has likely undergone recombination events. Addressing the third concern, point-mutation analysis can be performed in the SARS-CoV-2 UTR and TR to determine their effects on viral fitness.

## Conclusion

While the relative abundance of different selection types in a genome have been qualitatively characterized for the four evolution theories (GSRM, KNT, ONNT and ST) [4] and while numerous studies assigned Ka/Ks selection types for the SARS-CoV-2 All-TR region (**Table S3**), the percentage of genomic sites under these selection types have not been quantified to determine if any of these theories is true for this virus. To the best of our knowledge, our *c/μ* test is the first to quantify the percentage of sites in the genome undergoing neutral and non-neutral selection types, which was made possible after establishment of the fundamental genomic mutation rate (*μ*, FGMR) in our framework. Our relative substitution rate (*c/μ*) method was used to assign selection type to each gene, UTR and TRS of SARS-CoV-2 using the selected genome sequences by the Nextstrain team. As expected from our theoretical proof, *c/μ* selection type assignment was generally consistent with that of Ka/Ks for the TR. Therefore, the *c/μ* test can likely reliably assign selection type for the UTR and TRS, which are critical for viral replication and generation of coding sgRNA. The top 54 NT sites in UTR and 247 codon sites in TR which exhibited positive selection were also identified via our *c/μ* analysis, which are consistent with literature reports on their function and could be potential sites for drug intervention. Incredibly, *c/μ* exhibited an L-shaped probability distribution instead of the Poisson probability distribution for the genome and each gene as suggested by GSRM and KNT or the asymmetric distribution predicted by ONNT, both distribution should be centered at 1 (neutral selection) or the distribution lack of nearly neutral sites predicted by ST, thus this data does not support these three theories but instead supports the NNBST. In this novel hybrid mechanism, the lower substitution rate of the negative selection sites is balanced out with the higher substitution rate of the positive selection sites, and the effective population dependent effects on the weakly negative sites is also balanced that on the weakly positive sites size, thus resulting in the observed constant substitution rate. This sharply differs to ONNT in which nearly neutral sites are only slightly deleterious and thus their substitution rates are modulated by the effective population size. Analysis of the NT substitution variation over time showed that while the substitution rate (*c*) for the genome was time-independent regardless of the census population of infected persons and vaccinations, the *c for* some genes, UTRs and TRSs were time-dependent and followed strong negative selection (E, Orf6, Orf1ab 5’UTR) or strong positive selection (N, Orf8). This again indicated that the overall genome was under the NNBST: while particular genes or NT sites followed weak or strong adaptive selection, the others followed weak or strong purifying selection. Our analysis suggests that the molecular clock feature is not sufficient to support KNT or ONNT. Lastly, positive correlation between 1-dose vaccinations and Ka/Ks in most TR genes suggests these genes might respond to the vaccination by being involved in the host immune response. In future projects, our *c/μ* method will investigate the substitution rates of other RNA viruses in infected humans (i.e., Influenza, HIV) as well as coronaviruses in infected non-human host species (i.e., camels, bats) in both intraspecies and cross-species infections. To the best of our knowledge, our *c/μ* analysis was the first to quantify the percentage of NT sites in a genome undergoing neutral and non-neutral selection types, which provides a tool to resolve the long standing “neutralist-selectionist” controversy. Our NNBST in which the sequence variation is independent from the census population size might offer an explanation to Lewontin’s paradox in which pair-wise sequence variation is lack of proportionality to the census population.

## Acknowledgements

We thank Dhrumi C. Patel, Emily Dean, Lucas Bennett, Dylan J. Brunt, Michael J. Pino Jr., Katherine R. Hausman, Annie Tran, Mursalin Singh, Meeraj Amin, Justin D. Carbone and Julia Gabriel for compiling meta data and manual counting. Thank Brian Chen for writing a C-code program for early analysis. Thank Dr. Yong Chen for insightful discussions and comments on our manuscript. C.W acknowledges the support by the New Jersey Health Foundation (PC 76-21) and the National Science Foundation under Grants NSF ACI-1429467/RUI-1904797, and XSEDE MCB 170088. The Anton2 machine at the Pittsburgh Supercomputing Center (PSCA170090P) was generously made available by D. E. Shaw Research.

## Author Contributions

Conceptualization and Equation Derivation, C.W.; MatLab scripts, C.W. and N.J.P.; Data preprocessing, N.J.P. and P.M.L.; Plots and Tables, N.J.P. and P.M.L. and M.H.; writing—original draft preparation, N.J.P. and C.W.; writing—review and editing, C.W.; funding acquisition, C.W. All authors have read and agreed to the published version of the manuscript.

---

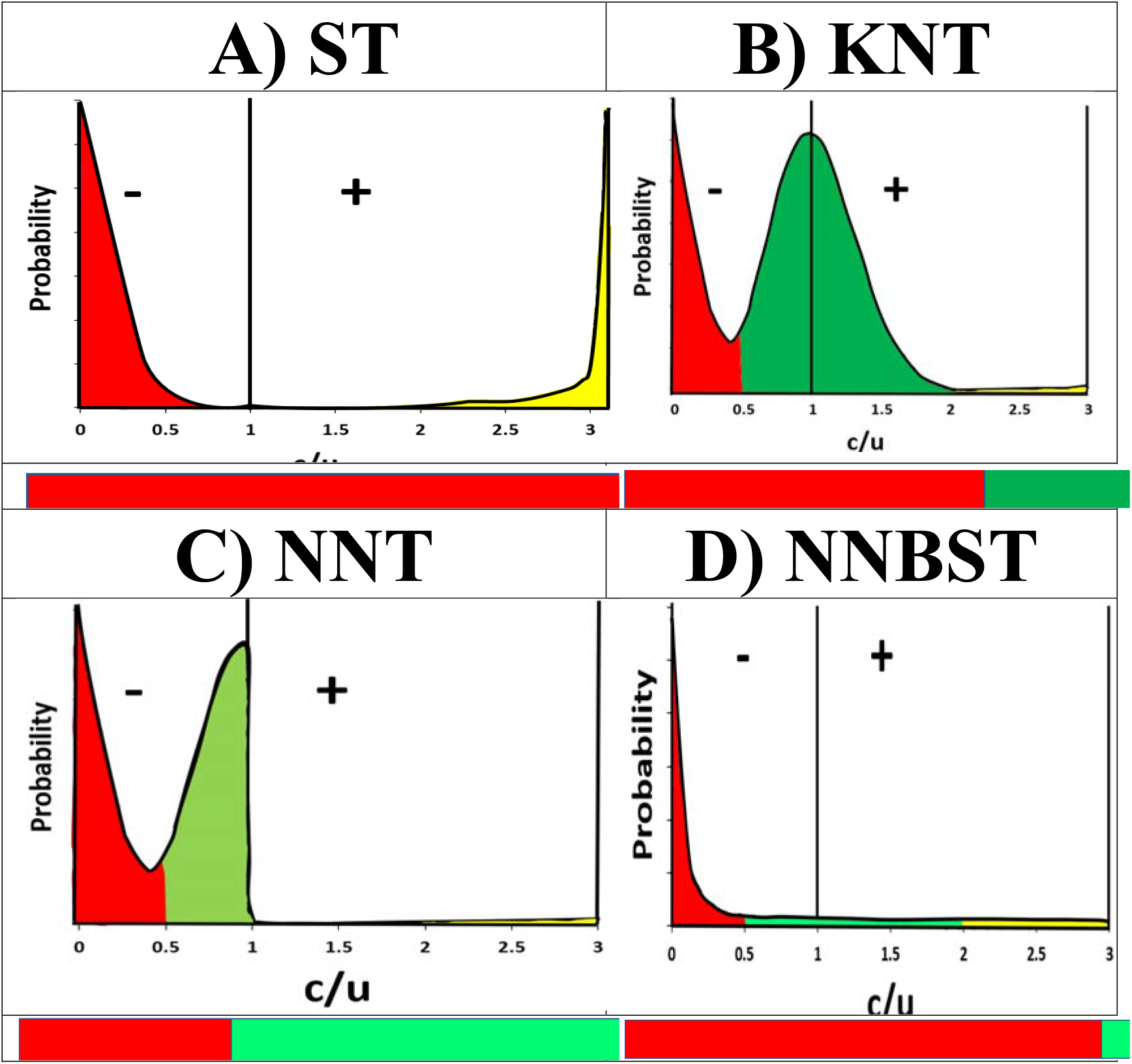
Diagrams of the four evolution theories **A**: ST; **B**: Poisson probability distribution implied by KNT/GSRM; **C**: Asymmetric distribution of the nearly neutral site implied by ONNT. **D**: L-Distribution theory implied by the NNBST. **Top**: Probability distribution showing negative (−), positive (+) and neutral selection (straight line). **Bottom**: Selection type distributions for the four theories of molecular evolution. Colors are assigned for strong negative/conserved selection (red), neutral selection (dark green), weak positive/weak negative selection (light green) and strong positive selection (yellow). Regions enclosed in the dashed lines in **C)** indicate weak positive selection (left) and weak negative selection (right).

## References

[1] W.S. Lee, A.K. Wheatley, S.J. Kent, B.J. DeKosky, Antibody-dependent enhancement and SARS-CoV-2 vaccines and therapies, Nat. Microbiol, 5 (2020) 1185–1191.

[2] V.K. Bhardwaj, R. Singh, P. Das, R. Purohit, Evaluation of acridinedione analogs as potential SARS-CoV-2 main protease inhibitors and their comparison with repurposed anti-viral drugs, Comput. Biol. Med., 128 (2021) 1–12.

[3] R. Singh, V.K. Bhardwaj, R. Purohit, Potential of turmeric-derived compounds against RNA-dependent RNA polymerase of SARS-CoV-2: An in-silico approach, Comput. Biol. Med., 139 (2021) 1–6.

[4] L. Bromham, D. Penny, The modern molecular clock, Nat. Rev. Genet., 4 (2003) 216–224.

[5] T. Gojobori, E.N. Moriyama, M. Kimura, Molecular clock of viral evolution, and the neutral theory, Proc. Natl. Acad. Sci. U. S. A., 87 (1990) 10015–10018.

[6] M. Kimura, Evolutionary rate at the molecular level, Nature, 217 (1968) 624–626.

[7] M. Kimura, The neutral theory of molecular evolution, Cambridge University Press, Cambridge, 1983.

[8] T. Ohta, Near-neutrality in evolution of genes and gene regulation, Proc. Natl. Acad. Sci. U. S. A., 99 (2002) 16134–16137.

[9] T. Ohta, Slightly deleterious mutant substitutions in evolution, Nature, 246 (1973) 96–98.

[10] X.L. Tang, C.C. Wu, X. Li, Y.H. Song, X.M. Yao, X.K. Wu, Y.G. Duan, H. Zhang, Y.R. Wang, Z.H. Qian, J. Cui, J. Lu, On the origin and continuing evolution of SARS-CoV-2, Natl. Sci. Rev., 7 (2020) 1012–1023.

[11] C. Roy, S.M. Mandal, S.K. Mondal, S. Mukherjee, T. Mapder, W. Ghosh, R. Chakraborty, Trends of mutation accumulation across global SARS-CoV-2 genomes: Implications for the evolution of the novel coronavirus, Genomics, 112 (2020) 5331–5342.

[12] S.M. Chaw, J.H. Tai, S.L. Chen, C.H. Hsieh, S.Y. Chang, S.H. Yeh, W.S. Yang, P.J. Chen, H.Y. Wang, The origin and underlying driving forces of the SARS-CoV-2 outbreak, J. Biomed. Sci., 27 (2020) 1–12.

[13] B. Dearlove, E. Lewitus, H.J. Bai, Y.F. Li, D.B. Reeves, M.G. Joyce, P.T. Scott, M.F. Amare, S. Vasan, N.L. Michael, K. Modjarrad, M. Rolland, A SARS-CoV-2 vaccine candidate would likely match all currently circulating variants, Proc. Natl. Acad. Sci. U. S. A., 117 (2020) 23652–23662.

[14] M.R. Garvin, E.T. Prates, M. Pavicic, P. Jones, B.K. Amos, A. Geiger, M.B. Shah, J. Streich, J. Gazolla, D. Kainer, A. Cliff, J. Romero, N. Keith, J.B. Brown, D. Jacobson, Potentially adaptive SARS-CoV-2 mutations discovered with novel spatiotemporal and explainable AI models, Genome Biol., 21 (2020) 1–26.

[15] G. Tonkin-Hill, I. Martincorena, R. Amato, A.R.J. Lawson, M. Gerstrung, I. Johnston, D.K. Jackson, N. Park, S.V. Lensing, M.A. Quail, S. Goncalves, C. Ariani, M.S. Chapman, W.L. Hamilton, L.W. Meredith, G. Hall, A.S. Jahun, Y. Chaudhry, M. Hosmillo, M.L. Pinckert, I. Georgana, A. Yakovleva, L.G. Caller, S.L. Caddy, T. Feltwell, F.A. Khokhar, C.J. Houldcroft, M.D. Curran, S. Parmar, A. Alderton, R. Nelson, E.M. Harrison, J. Sillitoe, S.D. Bentley, J.C. Barrett, M.E. Torok, I.G. Goodfellow, C. Langford, D.P. Kwiatowski, C.-G.U.C.-U. Consor, C.-S. Wellcome Sanger Inst, Patterns of within-host genetic diversity in SARS-CoV-2, eLife, 10 (2021) 1–25.

[16] K.J. Yi, S.Y. Kim, T. Bleazard, T. Kim, J. Youk, Y.S. Ju, Mutational spectrum of SARS-CoV-2 during the global pandemic, Exp. Mol. Med., 53 (2021) 1229–1237.

[17] L.F. Wang, B.T. Eaton, Bats, civets and the emergence of SARS, in: J.E. Childs, J.S. Mackenzie, J.A. Richt (Eds.) Wildlife and Emerging Zoonotic Diseases: The Biology, Circumstances and Consequences of Cross-Species Transmission 2007, pp. 325–344.

[18] L. Yang, Z.Q. Wu, X.W. Ren, F. Yang, G.M. He, J.P. Zhang, J. Dong, L.L. Sun, Y.F. Zhu, J. Du, S. Zhang, Q. Jin, Novel SARS-like betacoronaviruses in bats, China, 2011, Emerg Infect Dis., 19 (2013) 989–991.

[19] A. Chafekar, B.C. Fielding, MERS-CoV: understanding the latest human coronavirus threat, Viruses-Basel, 10 (2018) 1–22.

[20] A. Mubarak, W. Alturaiki, M.G. Hemida, Middle East Respiratory Syndrome Coronavirus (MERS-CoV): infection, immunological response, and vaccine development, J Immunol. Res., 2019 (2019) 1–11.

[21] C. Cao, Z. Cai, X. Xiao, J. Rao, J. Chen, N. Hu, M. Yang, X. Xing, Y. Wang, M. Li, B. Zhou, X. Wang, J. Wang, Y. Xue, The architecture of the SARS-CoV-2 RNA genome inside virion, Nat. Commun., 12 (2021) 3917–3931.

[22] D. Yang, J.L. Leibowitz, The structure and functions of coronavirus genomic 3’ and 5’ ends, Virus Res., 206 (2015) 120–133.

[23] D. Kim, J.Y. Lee, J.S. Yang, J.W. Kim, V.N. Kim, H. Chang, The architecture of SARS-CoV-2 transcriptome, Cell, 181 (2020) 914–921.

[24] J.S. Pita, J.R. de Miranda, W.L. Schneider, M.J. Roossinck, Environment determines fidelity for an RNA virus replicase, J Virol., 81 (2007) 9072–9077.

[25] V.K. Pathak, H.M. Temin, 5-Azacytidine and RNA Secondary Structure Increase the Retrovirus Mutation Rate, J. Virol., 66 (1992) 3093–3100.

[26] J.W. Drake, Too many mutants with multiple mutations, Crit. Rev. Biochem. Mol. Biol., 42 (2007) 247–258.

[27] V.K. Pathak, H.M. Temin, 5-Azacytidine and RNA secondary structure increase the retrovirus mutation-rate, J. Virol., 66 (1992) 3093–3100.

[28] S.H. Kim, N. Elango, C. Warden, E. Vigoda, S.V. Yi, Heterogeneous genomic molecular clocks in primates, PLoS Genet., 2 (2006) 1527–1534.

[29] J. Meunier, A. Khelifi, V. Navratil, L. Duret, Homologly-dependent methylation in primate repetitive DNA, Proc Natl Acad Sci U. S. A., 102 (2005) 5471–5476.

[30] L. Duret, P.F. Arndt, The impact of recombination on nucleotide substitutions in the human genome, PLoS Genet., 4 (2008) 1–19.

[31] N. Galtier, L. Duret, Adaptation or biased gene conversion? Extending the null hypothesis of molecular evolution, Trends Genet., 23 (2007) 273–277.

[32] M. Lynch, The origins of eukaryotic gene structure, Mol Biol Evol., 23 (2006) 450–468.

[33] S. Kumar, Molecular clocks: four decades of evolution, Nat. Rev. Genet., 6 (2005) 654–662.

[34] A.J. Drummond, S.Y.W. Ho, M.J. Phillips, A. Rambaut, Relaxed phylogenetics and dating with confidence, PLoS. Biol., 4 (2006) 699–710.

[35] J.H. Chen, R. Wang, N.B. Gilby, G.W. Wei, Omicron variant (B.1.1.529): infectivity, vaccine breakthrough, and antibody resistance, Journal of Chemical Information and Modeling, 62 (2022) 412–422.

[36] R. Wang, Y. Hozumi, C.C. Yin, G.W. Wei, Decoding SARS-CoV-2 transmission and evolution and ramifications for COVID-19 diagnosis, Vaccine, and medicine, Journal of Chemical Information and Modeling, 60 (2020) 5853–5865.

[37] R. Singh, V.K. Bhardwaj, P. Das, R. Purohit, A computational approach for rational discovery of inhibitors for non-structural protein 1 of SARS-CoV-2, Comput. Biol. Med., 135 (2021) 1–9.

[38] P. Kashyap, V.K. Bhardwaj, M. Chauhan, V. Chauhan, A. Kumar, R. Purohit, A. Kumar, S. Kumar, A ricin-based peptide BRIP from Hordeum vulgare inhibits M-pro of SARS-CoV-2, Sci Rep, 12 (2022) 1–11.

[39] J. Sharma, V.K. Bhardwaj, R. Singh, V. Rajendran, R. Purohit, S. Kumar, An in-silico evaluation of different bioactive molecules of tea for their inhibition potency against non structural protein-15 of SARS-CoV-2, Food Chem., 346 (2021) 1–8.

[40] J. Hadfield, C. Megill, S.M. Bell, J. Huddleston, B. Potter, C. Callender, P. Sagulenko, T. Bedford, R.A. Neher, Nextstrain: real-time tracking of pathogen evolution, Bioinformatics, 34 (2018) 4121–4123.

[41] P. Sagulenko, V. Puller, R.A. Neher, TreeTime: Maximum-likelihood phylodynamic analysis, Virus Evol., 4 (2018) 1–9.

[42] S. Duchene, L. Featherstone, M. Haritopoulou-Sinanidou, A. Rambaut, P. Lemey, G. Baele, Temporal signal and the phylodynamic threshold of SARS-CoV-2, Virus Evol., 6 (2020) 1–8.

[43] E. Calloway, The coronavirus is mutating-does it matter?, Nature, 585 (2020) 174–177.

[44] T. Koyama, D. Platt, L. Parida, Variant analysis of SARS-CoV-2 genomes, Bull. World Health Organ., 98 (2020) 495–504.

[45] D.S. Candido, I.M. Claro, J.G. de Jesus, W.M. Souza, F.R.R. Moreira, S. Dellicour, T.A. Mellan, L. du Plessis, R.H.M. Pereira, F.C.S. Sales, E.R. Manuli, J. Theze, L. Almeida, M.T. Menezes, C.M. Voloch, M.J. Fumagalli, T.M. Coletti, C.A.M. Silva, M.S. Ramundo, M.R. Amorim, H.H. Hoeltgebaum, S. Mishra, M.S. Gill, L.M. Carvalho, L.F. Buss, C.A. Prete, J. Ashworth, H.I. Nakaya, P.S. Peixoto, O.J. Brady, S.M. Nicholls, A. Tanuri, T.D. Rossi, C.K.V. Braga, A.L. Gerber, A.P.D. Guimaraes, N. Gaburo, C.S. Alencar, A.C.S. Ferreira, C.X. Lima, J.E. Levi, C. Granato, G.M. Ferreira, R.S. Francisco, F. Granja, M.T. Garcia, M.L. Moretti, M.W. Perroud, T. Castineiras, C.S. Lazari, S.C. Hill, A.A.D. Santos, C.L. Simeoni, J. Forato, A.C. Sposito, A.Z. Schreiber, M.N.N. Santos, C.Z. de Sa, R.P. Souza, L.C. Resende-Moreira, M.M. Teixeira, J. Hubner, P.A.F. Leme, R.G. Moreira, M.L. Nogueira, N.M. Ferguson, S.F. Costa, J.L. Proenca-Modena, A.T.R. Vasconcelos, S. Bhatt, P. Lemey, C.H. Wu, A. Rambaut, N.J. Loman, R.S. Aguiar, O.G. Pybus, E.C. Sabino, N.R. Faria, U.K.C.A.D. Brazil, Evolution and epidemic spread of SARS-CoV-2 in Brazil, Science, 369 (2020) 1255–1260.

[46] X.G. Li, W. Wang, X.F. Zhao, J.J. Zai, Q. Zhao, Y. Li, A. Chaillon, Transmission dynamics and evolutionary history of 2019-nCoV, J. Med. Virol., 92 (2020) 501–511.

[47] X.G. Li, J.J. Zai, Q. Zhao, Q. Nie, Y. Li, B.T. Foley, A. Chaillon, Evolutionary history, potential intermediate animal host, and cross-species analyses of SARS-CoV-2, J. Med. Virol., 92 (2020) 602–611.

[48] A. Eyre-Walker, P.D. Keightley, The distribution of fitness effects of new mutations, Nat. Rev. Genet., 8 (2007) 610–618.

[49] L. Kozlovskaya, A. Piniaeva, G. Ignatyev, A. Selivanov, A. Shishova, A. Kovpak, I. Gordeychuk, Y. Ivin, A. Berestovskaya, E. Prokhortchouk, D. Protsenko, M. Rychev, A. Ishmukhametov, Isolation and phylogenetic analysis of SARS-CoV-2 variants collected in Russia during the COVID-19 outbreak, Int. J. Infect. Dis., 99 (2020) 40–46.

[50] M. Mukherjee, S. Goswami, Global cataloguing of variations in untranslated regions of viral genome and prediction of key host RNA binding protein-microRNA interactions modulating genome stability in SARS-CoV-2, PLoS One, 15 (2020) 1–20.

[51] C.C. Yin, Genotyping coronavirus SARS-CoV-2: methods and implications, Genomics, 112 (2020) 3588–3596.

[52] R. Wang, J.H. Chen, K.F. Gao, Y. Hozumi, C.C. Yin, G.W. Wei, Analysis of SARS-CoV-2 mutations in the United States suggests presence of four substrains and novel variants, Commun. Biol., 4 (2021) 1–14.

[53] T.A.T. Soratto, H. Darban, A. Bjerkner, M. Coorens, J. Albert, T. Allander, B. Andersson, Four SARS-CoV-2 genome sequences from late april in Stockholm, Sweden, reveal a rare mutation in the spike protein, Microbiol. Resour. Ann., 9 (2020) 1–3.

[54] I. Saha, N. Ghosh, D. Maity, N. Sharma, J.P. Sarkar, K. Mitra, Genome-wide analysis of Indian SARS-CoV-2 genomes for the identification of genetic mutation and SNP, Infect. Genet. Evol., 85 (2020) 1–9.

[55] N.K. Ghanchi, A. Nasir, K.I. Masood, S.H. Abidi, S.F. Mahmood, A. Kanji, S. Razzak, W. Khan, S. Shahid, M. Yameen, A. Raza, J. Ashraf, Z. Ansar, M.B. Dharejo, N. Islam, Z. Hasan, R. Hasan, Higher entropy observed in SARS-CoV-2 genomes from the first COVID-19 wave in Pakistan, PLoS One, 16 (2021) e0256451.

[56] S.P. Ryder, B.R. Morgan, P. Coskun, K. Antkowiak, F. Massi, Analysis of emerging variants in structured regions of the SARS-CoV-2 genome, Evol. Bioinf., 17 (2021) 1–18.

[57] Z.C. Miao, A. Tidu, G. Eriani, F. Martin, Secondary structure of the SARS-CoV-2 5’-UTR, RNA Biol., 18 (2021) 447–456.

[58] J.X. Zhao, J.M. Qiu, S. Aryal, J.L. Hackett, J.X. Wang, The RNA architecture of the SARS-CoV-2 3 ’-untranslated region, Viruses-Basel, 12 (2020) 1–13.

[59] S. Elbe, G. Buckland-Merrett, Data, disease and diplomacy: GISAID’s innovative contribution to global health, Glob. Chall., 1 (2017) 33–46.

[60] T.H. Jukes, C.R. Cantor, Evolution of protein molecules, in: H.N. Munro (Ed.) Mammalian Protein Metabolism, Academic Press, New York, 1969, pp. 21–132.

[61] J.P. Huelsenbeck, B. Larget, D. Swofford, A compound Poisson process for relaxing the molecular clock, Genetics, 154 (2000) 1879–1892.

[62] L. Chao, C.U. Rang, L.E. Wong, Distribution of spontaneous mutants and inferences about the replication mode of the RNA bacteriophage phi 6, J. Virol., 76 (2002) 3276–3281.

[63] J. Felsenstein, Taking variation of evolutionary rates between sites into account in inferring phylogenies, J. Mol. Evol., 53 (2001) 447–455.

[64] S. Duffy, L.A. Shackelton, E.C. Holmes, Rates of evolutionary change in viruses: patterns and determinants, Nat. Rev. Genet., 9 (2008) 267–276.

[65] K. Katoh, J. Rozewicki, K.D. Yamada, MAFFT online service: multiple sequence alignment, interactive sequence choice and visualization, Brief. Bioinform., 20 (2019) 1160–1166.

[66] F. Corpet, Multiple sequence alignment with hierarchical-clustering, Nucleic Acids Res., 16 (1988) 10881–10890.

[67] M. Nei, T. Gojobori, Simple methods for estimating the numbers of synonymous and nonsynonymous nucleotide substitutions, Mol. Biol. Evol., 3 (1986) 418–426.

[68] M. Nei, L. Jin, Variances of the average numbers of nucleotide substitutions within and between populations, Mol. Biol. Evol., 6 (1989) 290–300.

[69] W.-H. Li, C.-I. Wu, C.-C. Luo, A new method for estimating synonymous and nonsynonymous rates of nucleotide substitution considering the relative likelihood of nucleotide and codon changes, Mol. Biol. Evol., 2 (1985) 150–174.

[70] P. Pamilo, N. Bianchi, Evolution of the Zfx and Zfy genes: rates and interdependence between the genes, Mol. Biol. Evol., 10 (1993) 271–281.

[71] Z. Yang, R. Nielsen, Estimating synonymous and nonsynonymous substitution rates under realistic evolutionary models, Mol Biol Cell, 17 (2000) 32–43.

[72] M. Kimura, A simple method for estimating evolutionary rates of base substitutions through comparative studies of nucleotide sequences J. Mol. Evol., 16 (1980) 111–120.

[73] N. Goldman, Z. Yang, A codon-based model of nucleotide substitution for protein-coding DNA sequences, Mol Biol Cell, 11 (1994) 725–736.

[74] W. Zheng, C. Zhang, Y. Li, R. Pearce, E.W. Bell, Y. Zhang, Folding non-homologous proteins by coupling deep-learning contact maps with I-TASSER assembly simulations, Cell Rep Methods, 1 (2021) 1–14,e11-e10.

[75] B. Berkhout, F. van Hemert, On the biased nucleotide composition of the human coronavirus RNA genome, Virus Research, 202 (2015) 41–47.

[76] N. Schmidt, C.A. Lareau, H. Keshishian, S. Ganskih, C. Schneider, T. Hennig, R. Melanson, S. Werner, Y. Wei, M. Zimmer, J. Ade, L. Kirschner, S. Zielinski, L. Dölken, E.S. Lander, N. Caliskan, U. Fischer, J. Vogel, S.A. Carr, J. Bodem, M. Munschauer, The SARS-CoV-2 RNA&#protein interactome in infected human cells, Nat. Microbiol., 6 (2021) 339–353.

[77] T. Mourier, M. Sadykov, M.J. Carr, G. Gonzalez, W.W. Hall, A. Pain, Host-directed editing of the SARS-CoV-2 genome, Biochem. Biophys. Res. Commun., 538 (2021) 35–39.

[78] R.D. Knight, S.J. Freeland, L.F. Landweber, A simple model based on mutation and selection explains trends in codon and amino-acid usage and GC composition within and across genomes Genome Biol., 2 (2001) 1–13.

[79] L. Bofkin, N. Goldman, Variation in evolutionary processes at different codon positions, Molecular Biology and Evolution, 24 (2007) 513–521.

[80] R. Sender, Y.M. Bar-On, S. Gleizer, B. Bernshtein, A. Flamholz, R. Phillips, R. Milo, The total number and mass of SARS-CoV-2 virions, Proc. Natl. Acad. Sci. U. S. A., 118 (2021) 1–9.

[81] M. Worobey, J. Pekar, B.B. Larsen, M.I. Nelson, V. Hill, J.B. Joy, A. Rambaut, M.A. Suchard, J.O. Wertheim, P. Lemey, The emergence of SARS-CoV-2 in Europe and North America, Science, 370 (2020) 564–570.

[82] K.M. Peck, A.S. Lauring, Complexities of viral mutation rates, J. Virol., 92 (2018) 1–8.

[83] G.M. Jenkins, A. Rambaut, O.G. Pybus, E.C. Holmes, Rates of molecular evolution in RNA viruses: A quantitative phylogenetic analysis, J. Mol. Evol., 54 (2002) 156–165.

[84] Z.S. Ma, J.D. Mei, Stochastic neutral drifts seem prevalent in driving human virome assembly: Neutral, near-neutral and non-neutral theoretic analyses, Comp. Struct. Biotechnol. J.. 20 (2022) 2029–2041.

[85] A. Kupczok, H. Neve, K.D. Huang, M.P. Hoeppner, K.J. Heller, C. Franz, T. Dagan, Rates of mutation and recombination in siphoviridae phage genome evolution over three decades, Molecular Biology and Evolution, 35 (2018) 1147–1159.

[86] S. Pignatelli, P. Dal Monte, G. Rossini, S. Chou, T. Gojobori, K. Hanada, J.J. Guo, W. Rawlinson, W. Britt, M. Mach, M.P. Landini, Human cytomegalovirus glycoprotein N (gpUL73-gN) genomic variants: identification of a novel subgroup, geographical distribution and evidence of positive selective pressure, J. Gen. Virol., 84 (2003) 647–655.

[87] S. Yi, Neutrality and molecular clocks, Nature Education, 4 (2013) 1–7.

[88] T.N. Starr, A.J. Greaney, S.K. Hilton, D. Ellis, K.H.D. Crawford, A.S. Dingens, M.J. Navarro, J.E. Bowen, M.A. Tortorici, A.C. Walls, N.P. King, D. Veesler, J.D. Bloom, Deep mutational scanning of SARS-CoV-2 receptor binding domain reveals constraints on folding and ACE2 binding, Cell, 182 (2020) 1295–1310.

[89] A.E. Chaouat, H. Achdout, I.B. Kol, O. Berhani, G. Roi, E. Vitner, S. Melamed, B. Politi, E. Zahavy, I. Brizic, T.L. Rovis, O. Alfi, D. Wolf, S. Jonjic, T. Israely, O. Mandelboim, K. Subbarao, SARS-CoV-2 receptor binding domain fusion protein efficiently neutralizes virus infection, PLoS Pathog., 17 (2021) 1–12.

[90] L. Min, Q. Sun, Antibodies and Vaccines Target RBD of SARS-CoV-2, Front. Mol. Biosci., 8 (2021) 9.

[91] N.G. Davies, S. Abbott, R.C. Barnard, C.I. Jarvis, A.J. Kucharski, J.D. Munday, C.A.B. Pearson, T.W. Russell, D.C. Tully, A.D. Washburne, T. Wenseleers, A. Gimma, W. Waites, K.L.M. Wong, K. van Zandvoort, J.D. Silverman, K. Diaz-Ordaz, R. Keogh, R.M. Eggo, S. Funk, M. Jit, K.E. Atkins, W.J. Edmunds, Estimated transmissibility and impact of SARS-CoV-2 lineage B.1.1.7 in England, Science, 372 (2021) 1–9.

[92] E. Volz, V. Hill, J.T. McCrone, A. Price, D. Jorgensen, A. O’Toole, J. Southgate, R. Johnson, B. Jackson, F.F. Nascimento, S.M. Rey, S.M. Nicholls, R.M. Colquhoun, A.D. Filipe, J. Shepherd, D.J. Pascall, R. Shah, N. Jesudason, K. Li, R. Jarrett, N. Pacchiarini, M. Bull, L. Geidelberg, I. Siveroni, I. Goodfellow, N.J. Loman, O.G. Pybus, D.L. Robertson, E.C. Thomson, A. Rambaut, T.R. Connor, C.-U. Consortium, Evaluating the effects of SARS-CoV-2 spike mutation D614G on transmissibility and pathogenicity, Cell, 184 (2021) 64–75.

[93] M.G. Hossain, Y.D. Tang, S. Akter, C.F. Zheng, Roles of the polybasic furin cleavage site of spike protein in SARS-CoV-2 replication, pathogenesis, and host immune responses and vaccination, J. Med. Virol., 94 (2022) 1815–1820.

[94] G.J. Olsen, Earliest phylogenetic branchings - comparing rRNA-based evolutionary trees inferred with various techniques, Cold Spring Harbor Symp. Quant. Biol., 52 (1987) 825–837.

[95] Z.H. Yang, Maximum-likelihood phylogenetic estimation from DNA-sequences with variable rates over sites - approximate methods, J. Mol. Evol., 39 (1994) 306–314.

[96] J.H. McDonald, M. Kreitman, Adaptive protein evolution at the adh locus in Drosophila, Nature, 351 (1991) 652–654.

[97] V.L. Cannataro, S.G. Gaffney, J.P. Townsend, Effect Sizes of Somatic Mutations in Cancer, Jnci-Journal of the National Cancer Institute, 110 (2018).

[98] M. Mohammadi-Dehcheshmeh, S.M. Moghbeli, S. Rahimirad, I.O. Alanazi, Z.S. Al Shehri, E. Ebrahimie, A transcription regulatory sequence in the 5 ’ untranslated region of SARS-CoV-2 is vital for virus replication with an altered evolutionary pattern against human inhibitory microRNAs, Cells, 10 (2021) 1–17.

[99] S.M. Hammond, An overview of microRNAs, Adv. Drug Deliv. Rev., 87 (2015) 3–14.

[100] S. Perlman, J. Netland, Coronaviruses post-SARS: update on replication and pathogenesis, Nat. Rev. Microbiol., 7 (2009) 439–450.

[101] P.S. Masters, The molecular biology of coronaviruses, in: K. Maramorosch, A.J. Shatkin (Eds.) Advances in Virus Research, Vol 66, Elsevier Academic Press Inc, San Diego, 2006, pp. 193–292.

[102] S.M. Vora, P. Fontana, T. Mao, V. Leger, Y. Zhang, T.-M. Fu, J. Lieberman, L. Gehrke, M. Shi, L. Wang, A. Iwasaki, H. Wu, Targeting stem-loop 1 of the SARS-CoV-2 5’ UTR to suppress viral translation and Nsp1 evasion, Proceedings of the National Academy of Sciences, 119 (2022) 1–10.

[103] V. Rohit, S. Sandhini, K. Shiv, M. Shailendra, K.M. Tushar, S. Milan, M.G. Marta, RNA-protein interaction analysis of SARS-CoV-2 5’ and 3’ untranslated regions reveals a role of lysosome-associated membrane protein-2a during viral infection, mSystems, 6 (2021) 1–24.

[104] A.D. Davidson, M.K. Williamson, S. Lewis, D. Shoemark, M.W. Carroll, K.J. Heesom, M. Zambon, J. Ellis, P.A. Lewis, J.A. Hiscox, D.A. Matthews, Characterisation of the transcriptome and proteome of SARS-CoV-2 reveals a cell passage induced in-frame deletion of the furin-like cleavage site from the spike glycoprotein, Genome Med., 12 (2020) 1–15.

[105] K. Pancer, A. Milewska, K. Owczarek, A. Dabrowska, M. Kowalski, P. Labaj, W. Branicki, M. Sanak, K. Pyrc, The SARS-CoV-2 ORF10 is not essential in vitro or in vivo in humans, PLoS Pathog., 16 (2020) 1–8.

[106] C. Castano-Rodriguez, J.M. Honrubia, J. Gutierrez-Alvarez, M.L. DeDiego, J.L. Nieto-Torres, J.M. Jimenez-Guardeno, J.A. Regla-Nava, R. Fernandez-Delgado, C. Verdia-Baguena, M. Queralt-Martin, G. Kochan, S. Perlman, V.M. Aguilella, I. Sola, L. Enjuanes, Role of Severe Acute Respiratory Syndrome Coronavirus viroporins E, 3a, and 8a in replication and pathogenesis, mBio, 9 (2018) 1–23.

[107] Y.J. Ren, T. Shu, D. Wu, J.F. Mu, C. Wang, M.H. Huang, Y. Han, X.Y. Zhang, W. Zhou, Y. Qiu, X. Zhou, The ORF3a protein of SARS-CoV-2 induces apoptosis in cells, Cell. Mol. Immunol., 17 (2020) 881–883.

[108] J.Y. Li, C.H. Liao, Q. Wang, Y.J. Tan, R. Luo, Y. Qiu, X.Y. Ge, The ORF6, ORF8 and nucleocapsid proteins of SARS-CoV-2 inhibit type I interferon signaling pathway, Virus Research, 286 (2020) 198074–198079.

[109] Z.G. Cao, H.J. Xia, R. Rajsbaum, X.Z. Xia, H.L. Wang, P.Y. Shi, Ubiquitination of SARS-CoV-2 ORF7a promotes antagonism of interferon response, Cell. Mol. Immunol., 18 (2021) 746–748.

[110] H.H. Wong, T.S. Fung, S.G. Fang, M. Huang, M.T. Le, D.X. Liu, Accessory proteins 8b and 8ab of severe acute respiratory syndrome coronavirus suppress the interferon signaling pathway by mediating ubiquitin-dependent rapid degradation of interferon regulatory factor 3, Virology, 515 (2018) 165–175.

[111] Y.W. Zhang, Y.S. Chen, Y.Z. Li, F. Huang, B.H. Luo, Y.C. Yuan, B.J. Xia, X.C. Ma, T. Yang, F. Yu, J. Liu, B.F. Liu, Z. Song, J.L. Chen, S.M. Yan, L.Y. Wu, T. Pan, X. Zhang, R. Li, W.J. Huang, X. He, F. Xiao, J.S. Zhang, H. Zhang, The ORF8 protein of SARS-CoV-2 mediates immune evasion through down-regulating MHC-I, Proc. Natl. Acad. Sci. U. S. A., 118 (2021) 1–12.

[112] L. Duret, Neutral theory: The null hypothesis of molecular evolution, Nature Education 1(2008) 218.

[113] S. Clancy, Genetic recombination, Nature Education, 1 (2008) 40.

[114] M. Kimura, J.F. Crow, The number of alleles that can be maintained in a finite population, Genetics, 49 (1964) 725–738.

[115] K.A. Lacek, B.L. Rambo-Martin, D. Batra, X.Y. Zheng, N. Hassell, H. Sakaguchi, T. Peacock, N. Groves, M. Keller, M.M. Wilson, M. Sheth, M.L. Davis, M. Borroughs, J. Gerhart, S.S. Shepard, P.W. Cook, J. Lee, D.E. Wentworth, J.R. Barnes, R. Kondor, C.R. Paden, SARS-CoV-2 delta-omicron recombinant viruses, United States, Emerg. Infect.Dis, 28 (2022) 1442–1445.

