## Supporting Document for "L-shape distribution of the relative substitution rate (c/μ) observed for SARS-COV-2’s genome, inconsistent with the selectionist theory, the neutral theory and the nearly neutral theory but a near-neutral balanced selection theory: implication on “neutralist-selectionist” debate"

**Table S1.** The primary structure of the SARS-CoV-2 reference genome (Wuhan-Hu-1/ MN908947.3), including nucleotide (NT)/amino acid (AA) lengths (**bolded**) and their residue positions within the genome (**parentheses**).

**A**). Lengths of Genome, All-TR and coding genes and proteins.

| Name | Genome | Orf1ab | S | Orf3a | E | M |
|---|---|---|---|---|---|---|
| #N.T. | **29,903** (1-29,903) | **21,291** (266-21,555) | **3,822** (21,563-25,384) | **828** (25,393-26,220) | **228** (26,245-26,472) | **669** (26,523-27,191) |
| #A.A | N/A | **7,097** (1-7,097) | **1,274** (7,098-8,371) | **276** (8,372-8,647) | **76** (8,648-8,723) | **223** (8,724-8,946) |
| **Name** | **All-TR** | **Orf6** | **Orf7a** | **Orf8** | **N** | **Orf10** |
| #N.T. | **29,133** (N/A) | **186** (27,202-27,387) | **366** (27,394-27,759) | **366** (27,894-28,259) | **1,260** (28,274-29,533) | **117** (29,559-29,674) |
| #A.A | **9,711** (1-9,711) | **62** (8,947-9,008) | **122** (9,009-9,130) | **122** (9,131-9,252) | **420** (9,253-9,672) | **39** (9,673-9,711) |

*Protein length includes the stop codon.
**All-TR consists of all protein-coding major and accessory genes
***Major genes/proteins: Orf1ab, S, E, M, N; Accessory genes/proteins: Orf3a, Orf6, Orf7a, Orf8, Orf10.

**B**). Lengths of non-structural proteins 1-15 (Nsp 1-15) coded in Orf1ab.

| Name | Nsp1 | Nsp2 | Nsp3/PL-PRO | Nsp4 | Nsp5/3CL-PRO |
|---|---|---|---|---|---|
| #N.T. | **540** (266-805) | **1,914** (806-2,719) | **5,835** (2,720-8,554) | **1,500** (8,555-10,054) | **918** (10,055-10,972) |
| #A.A | **180** (1-180) | **638** (181-818) | **1,945** (819-2,763) | **500** (2,764-3,263) | **306** (3,264-3,569) |
| **Name** | **Nsp6** | **Nsp7** | **Nsp8** | **Nsp9** | **Nsp10** |
| #N.T. | **870** (10,973-11,842) | **249** (11,843-12,091) | **594** (12,092-12,685) | **339** (12,686-13,024) | **417** (13,025-13,441) |
| #A.A | **290** (3,570-3,859) | **83** (3,860-3,942) | **198** (3,943-4,140) | **113** (4,141-4,253) | **139** (4,254-4,392) |
| **Name** | **Nsp11/RdRp** | **Nsp12/Helicase** | **Nsp13/ExoN** | **Nsp14/EndoRNase** | **Nsp15/OMT** |
| #N.T. | **2,796** (13,442-16,236) | **1,803** (16,237-18,039) | **1,581** (18,040-19,620) | **1,038** (19,621-20,658) | **894** (20,659-21,552) |
| #A.A | **932** (4,393-5,324) | **601** (5,325-5,925) | **527** (5,926-6,452) | **346** (6,453-6,798) | **298** (6,799-7,096) |

***PL-PRO**: Papain-like proteinase; **3CL-PRO**: 3C-like proteinase; **RdRp:** RNA-dependent RNA polymerase; **ExoN**: 3'-to-5' Exonuclease; **EndoRNase:** 5' Endoribonuclease; **OMT:** O-methyltransferase.

**Total length of Nsp 1-15s combined does not include Orf1ab STOP codon.

**C**). Lengths of All-UTR and UTRs.

| Name | All-UTR | Orf1ab 5'UTR | S 5'UTR | Orf3a 5'UTR | E 5'UTR | M 5'UTR |
|---|---|---|---|---|---|---|
| #N.T. | **771** (N/A) | **265** (1-265) | **7** (21,556-21,562) | **8** (25,385-25,392) | **24** (26,221-26,244) | **50** (26,473-26,522) |
| **Name** | **Orf6 5'UTR** | **Orf7a 5'UTR** | **Orf8 5'UTR** | **N 5'UTR** | **Orf10 5'UTR** | **Orf10 3'UTR** |
| #N.T. | **10** (27,192-27,201) | **6** (27,388-27,393) | **134** (27,760-27,893) | **14** (28,260-28,273) | **24** (29,534-29,558) | **229** (29,675-29,903) |

*All-UTR consists of each individual UTR sequence.
**5'-UTR and 3'-UTR are non-coding sequences and thus only have NT lengths.

**D).** Length of TRS-L and TRS-Bs.

| Name | All-TRS | Orf1ab TRS-L | S TRS-B | Orf3a TRS-B | E TRS-B | |
|---|---|---|---|---|---|---|
| #N.T. | **61** (N/A) | 7 (69-75) aacgaac | 7 (21,555-21,561) aacgaac | 7 (25,384-25,390) aacgaac | **6** (26,237-26,242) aacgaa | |
| **Name** | **M TRS-B** | **Orf6 TRS-B** | **Orf7a TRS-B** | **Orf8 TRS-B** | **N TRS-B** | **Orf10 TRS-B*** |
| #N.T. | 7 (26,472-26,478) aacgaac | **6** (27,041-27,046) aacgaa | 7 (27,387-27,393) aacgaac | 7 (27,887-27,893) aacgaac | 7 (28,259-28,265) aacgaac | 7 (29,551-29,557) aaggcag |

*All-TRS consists of each individual TRS-L and TRS-B sequence.
**TRS-L and TRS-Bs are non-coding sequences and thus only have NT lengths.
***Contains conserved 6-7 NT sequence below position within genome.
****Hypothetical Orf10 TRS-B, derived from MSA of the last 7 NTs in the Orf10 5' UTR to the reference sequence. Red and black colors represent non-conserved and conserved NTs, respectively.

**Table S2.** Substitution rates from the literature compared with our study (top).

| Substitution rate ( E-03 per NT site per year) | # of Genome | Date range studied | Method | NT sub mod | Clock type | Ref |
|---|---|---|---|---|---|---|
| 0.79±0.03 | 11,198 | Dec 2019-Jul 2021 | Linear Regression | JC69 | strict clock | This Study |
| 0.82 | 29,903 | Dec 2019-Sep 2020 | Linear Regression | JC69 | strict clock | Callaway |
| 1.1 | 606 | Jan 2020-Feb 2020 | BEAST v1.10 | HKY85 | strict clock | Duchene et al [42] |
| 1.12 | 10,022 | Feb 2020-May 2020 | BEAST v2.5 | HKY85 | strict clock | Koyama et al [44] |
| 1.13 (1.03-1.23) | 1,182 | Dec 2019-Apr 2020 | BEAST v.1.10.4 | HKY85 | strict clock | Candido et al [45] |
| 1.79-1.81 | 32 | Dec 2019-Jan 2020 | BEAST v1.8.4 | HKY85 | strict clock | Li et al [46] |
| 1.19 | 70 | Dec 2019-Feb 2020 | BEAST v1.8.4 | HKY85 | strict clock | Li et al [47] |
| 2.4 (1.5-3.3) | 137 | Dec 2019-Feb 2020 | BEAST 1.10.4 | HKY85 | uncorrected relaxed clock | Chaw et al [12] |

*JC69: Jukes TH, Cantor CR (1969). Evolution of Protein Molecules. New York: Academic Press. pp. 21–132.

**HKC85: Hasegawa M, Kishino H, Yano T (1985). "Dating of the human-ape splitting by a molecular clock of mitochondrial DNA". Journal of Molecular Evolution. 22 (2): 160–74.

***Methods are grouped based on linear regression or statistical models (e.g., BEAST vx.x.x).

**Table S3.** Ka/Ks and the selection pressures of the genome and each coding gene from available literature.

| Ref | This study | Yi et al [16] | Roy et al [11] | Tang et al [10] | Garvin et al [14] | Chaw et al [12] | Dearlove et al [13] | Tonkin-Hill et al [15] |
|---|---|---|---|---|---|---|---|---|
| **# of Genomes** | 11,779 | 351,525 | 71,703 | 103 | 15,789 | 137 | 18,514 | 1,181 |
| **Host** | Human | Human | Human | Human, Bat, Pangolin | Human | Human, Bat, Pangolin | Human | Human |
| **Collection Date** | Dec-19 -Jul 2021 | Dec 2019 -Jan 2021 | Dec 2019 -Aug 2020 | Dec 2019 -Jan 2021 | Dec 2019 -Jun 2020 | Dec 2019 -Feb 2020 | Dec 2019 -May 2020 | Mar 2020 -Apr 2020 |
| **Method** | NG LWL PBL ML | the 192-context model | HyPhy program | Free-ratio model CODEML in the PAML ZC45 as reference | Ka/Ks calculated from 385 haplotypes | LWL | Mixed-effect likelihood method | dNdScv |
| **Genome** | 0.8(-) | 0.5(-) | (-) | 0.081(-) | n/a | 0.32(-) | n/a | ~0.6(-) |
| **Orf1ab** | 0.4(-) | ~0.2-~0.7(-) | (-) | 0.058(-) | n/a | 1a/0.14, 1b/0.05(-) | ~0.6 (-) | n/a |
| **S** | 3.3(+) | 0.6(-) | 0.6193(-) | 0.13(-) | (+) | 0.013(-) | ~0.3-~0.8 (-) | n/a |
| **E** | 2.0(+) | ~1.0(0) | 1.0206(+) | 0(-) | n/a | 0(-) | ~0.2-~0.8 (-) | n/a |
| **M** | 0.5(-) | ~0.2(-) | 0.6548(-) | 0.055(-) | (-) | 0.15(-) | ~0.3 (-) | n/a |
| **N** | 1.2(+) | 0.7(-) | 1.2633(+) | 0.108(-) | (+) | 0.40(-) | ~0.8(-) | n/a |
| **Orf3a** | 2.4(+) | ~1.2(+) | 1.5013(+) | 0.102(-) | (+) | 1.83(+) | ~0.9-~1.7 (+) | n/a |
| **Orf6** | 0.5(-) | ~0.8(-) | 1.3944(+) | 0.209(-) | n/a | 0(-) | ~0.5-~0.8(-) | n/a |
| **Orf8** | 1.9(+) | ~0.9(-) | 1.4522(+) | 0.05(-) | n/a | 8.51(+) | ~0.8-~1.3(+) | n/a |
| **Orf10** | 1.2(+) | ~1.2(+) | 1.2981(+) | n/a | n/a | n/a | ~1.0-~1.9(+) | n/a |
| **Nsp 1** | 0.1(-) | ~0.4(-) | 0.7398(-) | n/a | (-) | n/a | n/a | n/a |
| **Nsp 2** | 0.3(-) | ~0.6(-) | 0.9479(-) | n/a | (+) | n/a | n/a | n/a |
| **Nsp 3** | 0.3(-) | ~0.5(-) | 0.5803(-) | n/a | (+) | n/a | n/a | n/a |
| **Nsp 4** | 0.6(-) | ~0.4(-) | 0.5126(-) | n/a | (-) | n/a | n/a | n/a |
| **Nsp 5** | 0.5(-) | ~0.3(-) | 0.6417(-) | n/a | n/a | n/a | n/a | n/a |
| **Nsp 6** | 0.7(-) | ~0.6(-) | 0.7000(-) | n/a | (+) | n/a | n/a | n/a |
| **Nsp 7** | 0.3(-) | ~0.2(-) | 0.4999(-) | n/a | (+) | n/a | n/a | n/a |
| **Nsp 8** | 0.5(-) | ~0.4(-) | 0.4892(-) | n/a | n/a | n/a | n/a | n/a |
| **Nsp 9** | 0.1(-) | ~0.3(-) | 0.4933(-) | n/a | n/a | n/a | n/a | n/a |
| **Nsp 10** | 0.2(-) | ~0.2(-) | 0.4187(-) | n/a | n/a | n/a | n/a | n/a |
| **Nsp 11** | 0.5(-) | ~0.4(-) | 0.6057(-) | n/a | (+) | n/a | n/a | n/a |
| **Nsp 12** | 0.8(-) | ~0.4(-) | 0.4500(-) | n/a | (+) | n/a | n/a | n/a |
| **Nsp 13** | 0.4(-) | ~0.5(-) | 0.4024(-) | n/a | (-) | n/a | n/a | n/a |
| **Nsp 14** | 0.3(-) | ~0.6(-) | 0.3937(-) | n/a | (+) | n/a | n/a | n/a |
| **Nsp 15** | 0.3(-) | ~0.5(-) | 0.4554(-) | n/a | n/a | n/a | n/a | n/a |

*Ka/Ks[b] results from **Table S7** from our study.

**Table S4**. Transition-Transversion NT substitutions counted for the genome and each gene in TR and UTR.

**A).** Percentage of each NT type in the reference genome sequence Wuhan-Hu-1.

| NT | A | G | U | C |
|---|---|---|---|---|
| **Rel. Abundance (%)** | 30 | 20 | 32 | 18 |

**B).** Transition (TS)-Transversion (TV) NT substitutions. Significant abundances are bolded.

| | | | | TS | | | | TV | | | | | | | |
|---|---|---|---|---|---|---|---|---|---|---|---|---|---|---|---|
| **Seg.** | **Tot TS TV** | **Tot TS** | **Tot TV** | **A-G** | **G-A** | **C-U** | **U-C** | **A-U** | **U-A** | **C-A** | **A-C** | **C-G** | **G-C** | **G-U** | **U-G** |
| **Genome** | 317828 | 218417 | 99411 | 32128 | 26588 | **136701** | 23000 | 11728 | 4578 | 14465 | 3624 | 5315 | 14112 | **39241** | 6348 |
| **All-TR** | 295011 | 203094 | 91917 | 31558 | 26088 | **122829** | 22619 | 10786 | 4088 | 14251 | 3412 | 5276 | 13951 | **33882** | 6271 |
| **All-UTR** | 22817 | 15323 | 7494 | 570 | 500 | **13872** | 381 | 942 | 490 | 214 | 212 | 39 | 161 | **5359** | 77 |
| **All-TRS** | 262 | 151 | 111 | 22 | 15 | 114 | 0 | 0 | 0 | 8 | 3 | 0 | 3 | 97 | 0 |
| **Orf1ab** | 139479 | 118682 | 20797 | 12942 | 6641 | **85535** | 13564 | 1239 | 699 | 4051 | 1389 | 88 | 344 | **12173** | 814 |
| **S** | 68052 | 33513 | 34539 | **12897** | 5607 | **13142** | 1867 | 4363 | 479 | **9666** | 1599 | 4144 | 3598 | **5502** | 5188 |
| **E** | 963 | 839 | 124 | 33 | 22 | **744** | 40 | 1 | 2 | 0 | 0 | 4 | 2 | **104** | 11 |
| **M** | 5073 | 4427 | 646 | 159 | 64 | **1650** | **2554** | 36 | 9 | 28 | 6 | 170 | 48 | 307 | 42 |
| **N** | 48627 | 24078 | 24549 | 2308 | **12409** | **8738** | 623 | 3642 | 2814 | 194 | 50 | 807 | **9695** | **7296** | 51 |
| **Orf3a** | 10883 | 6183 | 4700 | 251 | 250 | **4702** | 980 | 80 | 26 | 88 | 59 | 35 | 156 | **4210** | 46 |
| **Orf6** | 756 | 566 | 190 | 40 | 8 | **180** | **338** | 74 | 6 | 6 | 12 | 1 | 22 | 67 | 2 |
| **Orf7a** | 5447 | 4994 | 453 | 79 | 154 | **2784** | **1977** | 38 | 32 | 41 | 54 | 16 | 19 | 240 | 13 |
| **Orf8** | 14633 | 9238 | 5395 | **2823** | 869 | **4907** | 639 | 1304 | 12 | 165 | 238 | 9 | 46 | **3518** | 103 |
| **Orf10** | 720 | 420 | 300 | 24 | 21 | **338** | 37 | 6 | 9 | 4 | 5 | 1 | 2 | **272** | 1 |
| **Nsp1** | 2552 | 2243 | 309 | 166 | 172 | **951** | **954** | 6 | 28 | 31 | 20 | 1 | 10 | 199 | 14 |
| **Nsp2** | 12332 | 10729 | 1603 | 763 | 820 | **8609** | 537 | 534 | 87 | **114** | 100 | 6 | 17 | 695 | 50 |
| **Nsp3** | 48217 | 39609 | 8608 | 3304 | 1241 | **30787** | 4277 | 298 | 170 | 3129 | 865 | 24 | 40 | 3757 | 325 |
| **Nsp4** | 9908 | 7927 | 1981 | 436 | 319 | **6366** | 806 | 25 | 134 | 28 | 108 | 4 | 33 | **1617** | 32 |
| **Nsp5** | 3812 | 3420 | 392 | **977** | **574** | **1721** | 148 | 13 | 20 | 29 | 5 | 1 | 17 | 238 | 69 |
| **Nsp6** | 7413 | 5905 | 1508 | **3200** | 167 | **1718** | **820** | 22 | 41 | 215 | 13 | 3 | 12 | **1128** | 74 |
| **Nsp7** | 807 | 690 | 117 | 62 | 16 | **588** | 24 | 3 | 1 | 8 | 3 | 1 | 7 | 89 | 5 |
| **Nsp8** | 1251 | 1109 | 142 | **210** | 57 | **759** | 83 | 21 | 14 | 26 | 5 | 0 | 0 | 67 | 9 |
| **Nsp9** | 2600 | 2473 | 127 | 222 | 52 | **1909** | **290** | 5 | 4 | **14** | 3 | 2 | 0 | 87 | 12 |
| **Nsp10** | 738 | 526 | 212 | 27 | 65 | **332** | **102** | **104** | 13 | 17 | 5 | 1 | 0 | 67 | 5 |
| **Nsp11** | 28997 | 27790 | 1207 | 621 | 2296 | **20954** | **3919** | 45 | 51 | **111** | 36 | 7 | 10 | 860 | 87 |
| **Nsp12** | 8165 | 5931 | 2234 | **907** | 243 | **4372** | 409 | 41 | 45 | 57 | 133 | 19 | 4 | **1903** | 32 |
| **Nsp13** | 6348 | 5490 | 858 | 555 | 172 | **4328** | 435 | 17 | 41 | **149** | 27 | 5 | 8 | 574 | 37 |
| **Nsp14** | 3610 | 2955 | 655 | **1087** | 281 | **992** | 595 | 36 | 29 | 42 | 35 | 11 | 10 | 440 | 52 |
| **Nsp15** | 2729 | 1885 | 844 | **405** | 166 | **1149** | 165 | 69 | 21 | 81 | 31 | 3 | 176 | **452** | 11 |
| **Orf1ab 5'UTR** | 14862 | 11900 | 2962 | 125 | 85 | **11512** | 178 | 112 | 65 | 39 | 61 | 23 | 26 | **2582** | 54 |
| **S 5'UTR** | 5 | 4 | 1 | **1** | 0 | **3** | 0 | **1** | 0 | 0 | 0 | 0 | 0 | 0 | 0 |
| **E 5'UTR** | 63 | 41 | 22 | 1 | 1 | 38 | 1 | 1 | 1 | 4 | 0 | 0 | 0 | 16 | 0 |
| **M 5'UTR** | 182 | 28 | 154 | 3 | 0 | 7 | 18 | 103 | 1 | 0 | 0 | 0 | 1 | 48 | 1 |
| **N 5'UTR** | 727 | 181 | 546 | 156 | 10 | 2 | 13 | 540 | 0 | 5 | 1 | 0 | 0 | 0 | 0 |
| **Orf3a 5'UTR** | 1 | 1 | 0 | 1 | 0 | 0 | 0 | 0 | 0 | 0 | 0 | 0 | 0 | 0 | 0 |
| **Orf6 5'UTR** | 18 | 16 | 2 | 0 | 1 | 1 | 14 | 0 | 0 | 0 | 0 | 0 | 0 | 2 | 0 |
| **Orf7a 5'UTR** | 117 | 81 | 36 | 2 | 4 | 75 | 0 | 0 | 0 | 4 | 1 | 0 | 3 | 28 | 0 |
| **Orf8 5'UTR** | 1824 | 1524 | 300 | 27 | 19 | 1430 | 48 | 3 | 8 | 63 | 5 | 2 | 3 | 210 | 6 |
| **Orf10 3'UTR** | 5018 | 1547 | 3471 | 254 | 380 | 804 | 109 | 182 | 415 | 99 | 144 | 14 | 128 | 2473 | 16 |
| **Orf1ab TRS-L** | 3 | 3 | 0 | 2 | 0 | 1 | 0 | 0 | 0 | 0 | 0 | 0 | 0 | 0 | 0 |
| **S TRS-B** | 3 | 3 | 0 | 0 | 0 | 3 | 0 | 0 | 0 | 0 | 0 | 0 | 0 | 0 | 0 |
| **Orf3a TRS-B** | 3 | 3 | 0 | 3 | 0 | 0 | 0 | 0 | 0 | 0 | 0 | 0 | 0 | 0 | 0 |
| **E TRS-B** | 0 | 0 | 0 | 0 | 0 | 0 | 0 | 0 | 0 | 0 | 0 | 0 | 0 | 0 | 0 |
| **M TRS-B** | 0 | 0 | 0 | 0 | 0 | 0 | 0 | 0 | 0 | 0 | 0 | 0 | 0 | 0 | 0 |
| **Orf6 TRS-B** | 11 | 11 | 0 | 0 | 0 | 11 | 0 | 0 | 0 | 0 | 0 | 0 | 0 | 0 | 0 |
| **Orf7a TRS-B** | 117 | 81 | 36 | 2 | 4 | 75 | 0 | 0 | 0 | 4 | 1 | 0 | 3 | 28 | 0 |
| **Orf8 TRS-B** | 111 | 39 | 72 | 14 | 1 | 24 | 0 | 0 | 0 | 1 | 2 | 0 | 0 | 69 | 0 |
| **N TRS-B** | 14 | 11 | 3 | 1 | 10 | 0 | 0 | 0 | 0 | 3 | 0 | 0 | 0 | 0 | 0 |

**C).** Percentage of NT substitutions occurring in the first, second and third positions of each AA codon (P1, P2 and P3) for All-TR and each coding gene and Nsp (1-15).

**C1).** All-TR and each coding major/accessory gene.

| Codon | All-TR | Orf1ab | Non-Orf1ab | S | Orf3a | E | M | Orf6 | Orf7a | Orf8 | N | Orf10 | Sum |
|---|---|---|---|---|---|---|---|---|---|---|---|---|---|
| P1(%) | 28 | 12 | 16 | 7 | 1 | 0 | 0 | 0 | 0 | 2 | 5 | 0 | 16 |
| P2(%) | 44 | 17 | 27 | 13 | 2 | 0 | 1 | 0 | 1 | 2 | 7 | 0 | 27 |
| P3(%) | 28 | 18 | 10 | 3 | 1 | 0 | 1 | 0 | 0 | 1 | 4 | 0 | 10 |

Decomposition of All-TR into Orf1ab and non-Orf1ab. Decomposition of non-Orf1ab into genes.

**C2).** Decomposition of 12% of Orf1ab into Nsp1-15.

| Codon | Nsp1 | Nsp2 | Nsp3 | Nsp4 | Nsp5 | Nsp6 | Nsp7 | Nsp8 | Nsp9 | Nsp10 | Nsp11 | Nsp12 | Nsp13 | Nsp14 | Nsp15 | Sum |
|---|---|---|---|---|---|---|---|---|---|---|---|---|---|---|---|---|
| P1(%) | 0 | 1 | 2 | 1 | 0 | 1 | 0 | 0 | 0 | 0 | 4 | 1 | 1 | 0 | 0 | 12 |
| P2(%) | 0 | 1 | 6 | 1 | 0 | 1 | 0 | 0 | 0 | 0 | 4 | 1 | 1 | 1 | 1 | 17 |
| P3(%) | 1 | 2 | 8 | 1 | 0 | 1 | 0 | 0 | 1 | 0 | 1 | 0 | 1 | 0 | 0 | 18 |

**Table S5.** Percentage of each AA in the reference proteome sequence Wuhan-Hu-1 and AA substitution matrix for All-TR. Diagonal elements (yellow) represent synonymous substitutions. Off-diagonal elements represent non-synonymous substitutions. Total mutation count (bottom left cell), mutation count and percent mutation for each column (bottom two rows) are reported for each AA.

**A**). Percentage of each AA in the reference proteome sequence Wuhan-Hu-1.

| AA | A | C | D | E | F | G | H | I | K | L |
|---|---|---|---|---|---|---|---|---|---|---|
| Rel. Abundance (%) | 7 | 3 | 5 | 5 | 5 | 6 | 2 | 5 | 6 | 9 |
| **AA** | **M** | **N** | **P** | **Q** | **R** | **S** | **T** | **V** | **W** | **Y** |
| Rel. Abundance (%) | 2 | 5 | 4 | 4 | 4 | 7 | 7 | 8 | 1 | 5 |

**B).** Substitution in the Proteome.

| All-TR | A | R | N | D | C | Q | E | G | H | I | L | K | M | F | P | S | T | W | Y | V |
|---|---|---|---|---|---|---|---|---|---|---|---|---|---|---|---|---|---|---|---|---|
| A | 2566 | 0 | 0 | 5538 | 0 | 0 | 5 | 141 | 0 | 2 | 4 | 0 | 0 | 18 | 28 | 2877 | 542 | 0 | 5 | 6671 |
| R | 0 | 1167 | 114 | 2 | 563 | 59 | 3 | 84 | 123 | 2921 | 107 | 11858 | 2013 | 0 | 5 | 853 | 25 | 19 | 0 | 2 |
| N | 0 | 0 | 5389 | 376 | 0 | 0 | 0 | 2 | 39 | 35 | 4 | 365 | 0 | 0 | 0 | 321 | 92 | 0 | 4072 | 2 |
| D | 591 | 0 | 2482 | 6832 | 2 | 14 | 179 | 14114 | 3227 | 22 | 8251 | 2 | 0 | 12 | 0 | 4 | 6 | 0 | 3828 | 25 |
| C | 0 | 40 | 0 | 0 | 1012 | 0 | 0 | 4 | 0 | 2 | 0 | 0 | 0 | 244 | 0 | 29 | 0 | 6 | 38 | 0 |
| Q | 0 | 617 | 0 | 4 | 0 | 524 | 11 | 0 | 3468 | 0 | 176 | 146 | 0 | 0 | 33 | 0 | 0 | 0 | 8 | 0 |
| E | 104 | 4 | 14 | 1454 | 0 | 247 | 638 | 234 | 6 | 0 | 2 | 2782 | 0 | 0 | 0 | 0 | 0 | 0 | 2 | 201 |
| G | 72 | 5648 | 8 | 1136 | 2082 | 2 | 92 | 1975 | 0 | 0 | 36 | 0 | 0 | 8 | 444 | 2884 | 0 | 29 | 0 | 1097 |
| H | 0 | 64 | 12 | 19 | 0 | 139 | 0 | 0 | 3580 | 0 | 10 | 0 | 0 | 0 | 3 | 0 | 0 | 0 | 2004 | 0 |
| I | 0 | 1 | 5 | 0 | 0 | 0 | 0 | 0 | 0 | 2053 | 272 | 13 | 113 | 138 | 2 | 50 | 5642 | 0 | 2 | 1076 |
| L | 0 | 2256 | 0 | 0 | 0 | 177 | 3 | 0 | 20 | 218 | 7331 | 4 | 24 | 5056 | 392 | 433 | 0 | 11 | 10 | 212 |
| K | 0 | 1902 | 2051 | 0 | 0 | 728 | 202 | 2 | 0 | 9 | 4 | 1034 | 9 | 0 | 0 | 8 | 711 | 0 | 2 | 0 |
| M | 0 | 2 | 0 | 0 | 0 | 0 | 0 | 0 | 0 | 2054 | 60 | 18 | 1 | 0 | 0 | 0 | 147 | 0 | 0 | 190 |
| F | 0 | 0 | 0 | 0 | 31 | 0 | 0 | 0 | 5 | 16 | 837 | 0 | 0 | 17667 | 0 | 366 | 0 | 4 | 127 | 285 |
| P | 10 | 2903 | 0 | 0 | 0 | 79 | 0 | 0 | 3487 | 2 | 18840 | 2 | 0 | 240 | 5047 | 4768 | 111 | 0 | 0 | 0 |
| S | 2871 | 88 | 558 | 0 | 34 | 0 | 2 | 205 | 0 | 426 | 5272 | 5 | 0 | 4134 | 949 | 6260 | 103 | 0 | 114 | 0 |
| T | 2026 | 1899 | 982 | 0 | 0 | 2 | 8 | 0 | 0 | 20095 | 0 | 2326 | 186 | 0 | 19 | 63 | 7002 | 0 | 0 | 8 |
| W | 0 | 114 | 0 | 0 | 415 | 0 | 0 | 1 | 0 | 0 | 229 | 0 | 0 | 0 | 0 | 7 | 0 | 0 | 0 | 0 |
| Y | 0 | 0 | 111 | 2 | 2900 | 0 | 0 | 0 | 177 | 0 | 0 | 0 | 0 | 21 | 0 | 105 | 4 | 0 | 4830 | 0 |
| V | 2787 | 2 | 0 | 4 | 0 | 0 | 6 | 40 | 0 | 937 | 2626 | 0 | 196 | 1940 | 2 | 6 | 2 | 0 | 0 | 5529 |
| 290102 | 11027 | 16707 | 11726 | 15367 | 7039 | 1971 | 1149 | 16802 | 14132 | 28792 | 44061 | 18555 | 2542 | 29478 | 6924 | 19034 | 14387 | 69 | 15042 | 15298 |
| % | 3.8 | 5.8 | 4.0 | 5.3 | 2.4 | 0.7 | 0.4 | 5.8 | 4.9 | 9.9 | 15.2 | 6.4 | 0.9 | 10.2 | 2.4 | 6.6 | 5.0 | 0.0 | 5.2 | 5.3 |

**Table S6.** Decomposition of each NT/codon sequence into different selection types using *c/μ* test. The five categories of selection pressure are: conserved percentage (%C), strong negative percentage (%SN), weak negative percentage (%WN), weak positive percentage (%WP) and strong positive percentage (%SP) **(Figure 7**). **%C:** $c/\mu = 0$, **%SN**: $0 < c/\mu < 0.5$, **%WN:** $0.5 < c/\mu < 1.0$, **%WP:** $1.0 < c/\mu < 2.0$, **%SP:** $c/\mu > 2.0$.

**A).** All-TR, each major/accessory coding gene (Codon sequence).

| PRO | All-TR | Orf1ab | Non-Orf1ab | S | Orf3a | E | M | Orf6 | Orf7a | Orf8 | N | Orf10 |
|---|---|---|---|---|---|---|---|---|---|---|---|---|
| %C | 22 | 23 | 17 | 22 | 8 | 32 | 24 | 11 | 7 | 4 | 11 | 5 |
| %SN | 61 | 63 | 59 | 60 | 54 | 55 | 60 | 74 | 66 | 60 | 56 | 67 |
| %WN | 8 | 7 | 11 | 7 | 22 | 9 | 8 | 8 | 15 | 10 | 14 | 15 |
| %WP | 4 | 4 | 5 | 4 | 8 | 1 | 4 | 2 | 7 | 11 | 7 | 5 |
| %SP | 5 | 3 | 8 | 7 | 9 | 3 | 4 | 5 | 5 | 16 | 12 | 8 |
| Sel(+/-) | - | - | - | - | - | - | - | - | - | - | - | - |

**B).** Nsp1-15 (Codon sequence).

| PRO | Nsp1 | Nsp2 | Nsp3 | Nsp4 | Nsp5 | Nsp6 | Nsp7 | Nsp8 | Nsp9 | Nsp10 | Nsp11 | Nsp12 | Nsp13 | Nsp14 | Nsp15 |
|---|---|---|---|---|---|---|---|---|---|---|---|---|---|---|---|
| %C | 13 | 11 | 21 | 24 | 26 | 21 | 22 | 29 | 23 | 30 | 31 | 29 | 25 | 23 | 29 |
| %SN | 72 | 71 | 64 | 63 | 60 | 62 | 60 | 61 | 58 | 60 | 60 | 58 | 62 | 66 | 61 |
| %WN | 9 | 9 | 7 | 6 | 6 | 6 | 11 | 7 | 10 | 8 | 5 | 6 | 6 | 6 | 4 |
| %WP | 2 | 4 | 4 | 3 | 4 | 4 | 4 | 2 | 4 | 1 | 2 | 4 | 5 | 3 | 3 |
| %SP | 4 | 4 | 3 | 4 | 5 | 7 | 4 | 2 | 6 | 1 | 2 | 4 | 2 | 2 | 3 |
| Sel(+/-) | - | - | - | - | - | - | - | - | - | - | - | - | - | - | - |

*Orf1ab decomposed into Nsp (1-15).

**C).** UTR (NT sequence).

| Gene | All-UTR | Orf1ab 5'UTR | Non-Orf1ab 5'UTR | S 5'UTR | Orf3a 5'UTR | E 5'UTR | M 5'UTR | Orf6 5'UTR | Orf7a 5'UTR | Orf8 5'UTR | N |
|---|---|---|---|---|---|---|---|---|---|---|---|
| %C | 34 | 34 | 33 | 71 | 88 | 54 | 62 | 70 | 0 | 38 | |
| %SN | 63 | 63 | 63 | 29 | 13 | 46 | 36 | 30 | 83 | 60 | |
| %WN | 2 | 1 | 2 | 0 | 0 | 0 | 2 | 0 | 17 | 1 | |
| %WP | <1 | 0 | 1 | 0 | 0 | 0 | 0 | 0 | 0 | 0 | |
| %SP | <1 | 2 | 1 | 0 | 0 | 0 | 0 | 0 | 0 | 1 | |
| Sel (+/-) | - | - | - | - | - | - | - | - | - | - | |

**D).** TRS (NT sequence).

| Gene | All TRS | Orf1ab TRS-L | Non-Orf1ab TRS | S TRS-B | Orf3a TRS-B | E TRS-B | M TRS-B | Orf6 TRS-B | Orf7a T |
|---|---|---|---|---|---|---|---|---|---|
| %C | 67 | 57 | 69 | 86 | 71 | 100 | 100 | 83 | 14 |
| %SN | 30 | 43 | 28 | 14 | 29 | 0 | 0 | 17 | 71 |
| %WN | 3 | 0 | 4 | 0 | 0 | 0 | 0 | 0 | 14 |
| %WP | <1 | 0 | 0 | 0 | 0 | 0 | 0 | 0 | 0 |
| %SP | <1 | 0 | 0 | 0 | 0 | 0 | 0 | 0 | 0 |
| Sel (+/-) | - | - | - | - | - | - | - | - | - |

**Table S7.** Function and decomposition of the total NT substitution rates into Ka (non-synonymous NT) and Ks (synonymous NT) substitution rates using two types of methods (SP and averaged NG, LWL, PBL and ML methods). The rates (%NT site per 19 months) were calculated from the three data sets A1a/A1b/A1c.

| Seg (Length) | Function | Total NT sub rate | SP | | | NG, LWL, PBL and ML | | |
|---|---|---|---|---|---|---|---|---|
| | | | NonSyn | Syn | Ka/Ks[a] | NonSyn | Syn | Ka/Ks[b] (mean) |
| Genome(29,903) | | 0.0902 | n/a | n/a | n/a | n/a | n/a | n/a |
| All-TR(29,133) | **Proteome** | 0.0859 | 0.1873 | 0.0703 | 2.7 | 0.0382 | 0.0477 | 0.8±0.2 |
| All-UTR(771) | | 0.2554 | n/a | n/a | n/a | n/a | n/a | n/a |
| All-TRS(61) | | 0.0365 | n/a | n/a | n/a | n/a | n/a | n/a |
| Orf1ab(21,291) | Viral genome replication | 0.0556 | 0.0903 | 0.0766 | 1.2 | 0.0159 | 0.0397 | 0.4±0.1 |
| S(3,822) | **Viral infection and transmission** | 0.1512 | 0.4147 | 0.0388 | 10.7 | 0.1160 | 0.0352 | 3.3±0.7 |
| E(228) | Viral assembly, pathogenesis | 0.0359 | 0.0895 | 0.0181 | 4.9 | 0.0239 | 0.0120 | 2.0±0.4 |
| M(669) | Viral budding from host cell | 0.0644 | 0.1099 | 0.0833 | 1.3 | 0.0215 | 0.0429 | 0.5±0.1 |
| N(1,260) | **Packaging viral RNA, viral self-assembly** | 0.3277 | 0.8871 | 0.0960 | 9.2 | 0.1787 | 0.1490 | 1.2±0.2 |
| Orf3a(828) | **Viral release, improves transmission** | 0.1116 | 0.2957 | 0.0391 | 7.6 | 0.0788 | 0.0328 | 2.4±0.4 |
| Orf6(186) | Stunts import of biomolecules into host cell nucleus | 0.0345 | 0.0652 | 0.0383 | 1.7 | 0.0115 | 0.0230 | 0.5±0.2 |
| Orf7a(366) | Reduces antiviral activity, improves infection | 0.1264 | 0.3482 | 0.0309 | 11.3 | 0.0995 | 0.0269 | 3.7±0.7 |
| Orf8(366) | **Evade host immune response** | 0.3395 | 0.9216 | 0.0969 | 9.5 | 0.2224 | 0.1171 | 1.9±0.4 |
| Orf10(117) | N/A | 0.0523 | 0.1252 | 0.0318 | 3.9 | 0.0285 | 0.0238 | 1.2±0.3 |
| Nsp1(538) | Induces degradation of host cell mRNA; viral mRNA unaffected | 0.0401 | 0.0325 | 0.0878 | 0.4 | 0.0036 | 0.0365 | 0.1±0.0 |
| Nsp2(1,912) | Protects host cell mitochondria, decreases cell stressors, protects viral RNA | 0.0547 | 0.0754 | 0.0887 | 0.9 | 0.0126 | 0.0421 | 0.3±0.1 |
| Nsp3(5,388) | Cleaves N-terminus of Orf1ab; inhibits INF-I production, evades host cell immune response, improves infection | 0.0702 | 0.1088 | 0.1017 | 1.1 | 0.0162 | 0.0540 | 0.3±0.1 |
| Nsp4(1,498) | Double-membrane vesicle formation of host cells with Nsp3; initiates viral replication | 0.0561 | 0.1087 | 0.0595 | 1.8 | 0.0210 | 0.0351 | 0.6±0.2 |
| Nsp5(916) | Cleaves Orf1ab into 11 NSPs; facilitates NSP maturation | 0.0353 | 0.0656 | 0.0402 | 1.6 | 0.0118 | 0.0235 | 0.5±0.1 |
| Nsp6(868) | Induces/restricts autophagosome production, facilitates Orf1ab assembly from host cell ER | 0.0724 | 0.1487 | 0.0683 | 2.2 | 0.0298 | 0.0426 | 0.7±0.2 |
| Nsp7(247) | Forms hexadecamer with NSP8 as cofactor for NSP11 to drive viral replication | 0.0275 | 0.0404 | 0.0421 | 1.0 | 0.0063 | 0.0212 | 0.3±0.1 |
| Nsp8(592) | Forms hexadecamer with NSP7 as cofactor for NSP11 to drive viral replication | 0.0179 | 0.0311 | 0.0225 | 1.4 | 0.0060 | 0.0119 | 0.5±0.1 |
| Nsp9(337) | ssRNA-binding protein dimer; prevents viral RNA degradation | 0.0651 | 0.0513 | 0.1440 | 0.4 | 0.0059 | 0.0592 | 0.1±0.0 |
| Nsp10(415) | Methylates 5'-cap of viral RNA, increases viral RNA stability | 0.0150 | 0.0187 | 0.0264 | 0.7 | 0.0025 | 0.0125 | 0.2±0.0 |
| Nsp11(2,794) | **RNA-dependent RNA Polymerase (RdRp); replicates viral RNA from an RNA template strand** | 0.0881 | 0.1457 | 0.1185 | 1.2 | 0.0294 | 0.0587 | 0.5±0.1 |
| Nsp12(1,801) | Helicase; unwinds dsRNA, initiates viral replication | 0.0385 | 0.0835 | 0.0320 | 2.6 | 0.0171 | 0.0214 | 0.8±0.2 |
| Nsp13(1,579) | Exoribonuclease (ExoRNase); proofreads duplicate RNA strand, decreases nucleotide substitutions | 0.0341 | 0.0538 | 0.0486 | 1.1 | 0.0097 | 0.0244 | 0.4±0.1 |
| Nsp14(1,036) | Endoribonuclease (EndoRNase); similar role to NSP12 | 0.0295 | 0.04 | 0.0488 | 0.8 | 0.0068 | 0.0227 | 0.3±0.1 |
| Nsp15(892) | 2'-O-Methyltransferase (2'-OMT); methylates 5'-cap of viral RNA, increases viral RNA stability | 0.0259 | 0.04 | 0.0380 | 1.1 | 0.0060 | 0.0199 | 0.3±0.1 |

**Table S8.** The Ka/Ks for each gene calculated from data sets A1a/A1b/A1c averaged over the four methods and for each method.

| Seg (Length) | Ka/Ks NG(P/N) | Ka/Ks LWL(P/N) | Ka/Ks PBL(P/N) | Ka/Ks ML(P/N) | Ka/Ks[b] (mean) |
|---|---|---|---|---|---|
| **Genome(29,903)** | n/a | n/a | n/a | n/a | n/a |
| **All-TR(29,133)** | 0.7±0.0(-) | 0.6±0.0(-) | 1.0±0.1 | 0.7±0.0(-) | 0.8±0.2(-) |
| **All-UTR(771)** | n/a | n/a | n/a | n/a | n/a |
| **Orf1ab(21,291)** | 0.5±0.2(-) | 0.3±0.0(-) | 0.6±0.0(-) | 0.4±0.0(-) | 0.4±0.1(-) |
| **S(3,822)** | 3.2±0.4(+) | 2.9±0.4(+) | 4.2±0.5(+) | 2.7±0.3(+) | 3.3±0.7(+) |
| **E(228)** | 1.7±0.3(+) | 1.6±0.3(+) | 2.4±0.6(+) | 2.1±0.4(+) | 2.0±0.4(+) |
| **M(669)** | 0.4±0.1(-) | 0.4±0.1(-) | 0.6±0.2(-) | 0.6±0.2(-) | 0.5±0.1(-) |
| **N(1,260)** | 1.1±0.1(+) | 1.0±0.1 | 1.4±0.1(+) | 1.3±0.1(+) | 1.2±0.2(+) |
| **Orf3a(828)** | 2.2±0.2(+) | 2.0±0.1(+) | 2.9±0.2(+) | 2.5±0.2(+) | 2.4±0.4(+) |
| **Orf6(186)** | 0.4±0.1(-) | 0.3±0.1(-) | 0.7±0.2(-) | 0.5±0.2(-) | 0.5±0.2(-) |
| **Orf7a(366)** | 3.4±0.5(+) | 3.0±0.5(+) | 4.6±0.8(+) | 3.7±0.6(+) | 3.7±0.7(+) |
| **Orf8(366)** | 1.7±0.2(+) | 1.5±0.2(+) | 2.5±0.4(+) | 1.8±0.3(+) | 1.9±0.4(+) |
| **Orf10(117)** | 1.0±0.2 | 0.9±0.1(-) | 1.6±0.3(+) | 1.4±0.2(+) | 1.2±0.3(+) |
| **Nsp1(538)** | 0.1±0.0(-) | 0.1±0.0(-) | 0.2±0.0(-) | 0.2±0.0(-) | 0.1±0.0(-) |
| **Nsp2(1,912)** | 0.3±0.0(-) | 0.2±0.0(-) | 0.4±0.1(-) | 0.3±0.1(-) | 0.3±0.1(-) |
| **Nsp3(5,388)** | 0.3±0.0(-) | 0.3±0.0(-) | 0.5±0.0(-) | 0.3±0.0(-) | 0.3±0.1(-) |
| **Nsp4(1,498)** | 0.6±0.0(-) | 0.5±0.0(-) | 0.9±0.0(-) | 0.5±0.0(-) | 0.6±0.2(-) |
| **Nsp5(916)** | 0.5±0.0(-) | 0.5±0.0(-) | 0.7±0.1(-) | 0.5±0.0(-) | 0.5±0.1(-) |
| **Nsp6(868)** | 0.6±0.0(-) | 0.6±0.0(-) | 1.0±0.0 | 0.7±0.0(-) | 0.7±0.2(-) |
| **Nsp7(247)** | 0.3±0.1(-) | 0.3±0.0(-) | 0.4±0.1(-) | 0.4±0.1(-) | 0.3±0.1(-) |
| **Nsp8(592)** | 0.4±0.1(-) | 0.4±0.1(-) | 0.6±0.1(-) | 0.4±0.1(-) | 0.5±0.1(-) |
| **Nsp9(337)** | 0.1±0.0(-) | 0.1±0.0(-) | 0.2±0.0(-) | 0.1±0.0(-) | 0.1±0.0(-) |
| **Nsp10(415)** | 0.2±0.0(-) | 0.2±0.0(-) | 0.2±0.0(-) | 0.2±0.0(-) | 0.2±0.0(-) |
| **Nsp11(2,794)** | 0.4±0.1(-) | 0.3±0.1(-) | 0.6±0.1(-) | 0.5±0.1(-) | 0.5±0.1(-) |
| **Nsp12(1,801)** | 0.8±0.1(-) | 0.7±0.1(-) | 1.1±0.1(+) | 0.7±0.1(-) | 0.8±0.2(-) |
| **Nsp13(1,579)** | 0.3±0.0(-) | 0.3±0.0(-) | 0.5±0.0(-) | 0.3±0.0(-) | 0.4±0.1(-) |
| **Nsp14(1,036)** | 0.2±0.0(-) | 0.2±0.0(-) | 0.4±0.0(-) | 0.2±0.0(-) | 0.3±0.1(-) |
| **Nsp15(892)** | 0.3±0.0(-) | 0.3±0.0(-) | 0.3±0.0(-) | 0.2±0.0(-) | 0.3±0.1(-) |

[a] Ka/Ks calculated by dividing the non-synonymous substitution rate by the synonymous substitution rate. See **Table 3**.

[b] Ka/Ks calculated by taking the average of the absolute mean values for each of the four methods (NG, LWL, PBL and ML).

**Table S9**. Ka, Ks and Ka/Ks values calculated for each gene in sets A1a-A1c using the four methods, used to validate the average selection pressure assigned by c/u analysis. See Ka/Ks[b] (mean) values in **Tables 3 and S7-S8.**

**A).** Nei-Gojobori (NG) method.

| | A1a | | | A1b | | | A1c | | |
|---|---|---|---|---|---|---|---|---|---|
| Gene | Ka | Ks | Ka/Ks | Ka | Ks | Ka/Ks | Ka | Ks | Ka/Ks |
| All-TR | 3.1 | 4.4 | 0.7(-) | 3.3 | 4.5 | 0.7(-) | 2.7 | 4.0 | 0.7(-) |
| E | 1.8 | 0.9 | 2.0(+) | 1.7 | 0.9 | 1.9(+) | 1.3 | 1.0 | 1.4(+) |
| S | 7.4 | 2.3 | 3.3(+) | 7.9 | 2.2 | 3.6(+) | 5.9 | 2.1 | 2.8(+) |
| M | 1.8 | 4.1 | 0.4(-) | 2.3 | 4.2 | 0.5(-) | 1.6 | 5.2 | 0.3(-) |
| N | 13.9 | 12.4 | 1.1(+) | 14.5 | 12.0 | 1.2(+) | 11.6 | 10.5 | 1.1(+) |
| Orf1ab | 1.5 | 4.5 | 0.3(-) | 1.6 | 4.6 | 0.4(-) | 1.4 | 4.0 | 0.4(-) |
| Orf3a | 4.8 | 2.3 | 2.1(+) | 5.1 | 2.2 | 2.4(+) | 5.1 | 2.2 | 2.3(+) |
| Orf6 | 0.8 | 3.0 | 0.3(-) | 1.0 | 2.5 | 0.4(-) | 1.3 | 2.6 | 0.5(-) |
| Orf7a | 5.4 | 1.5 | 3.6(+) | 7.4 | 1.9 | 3.8(+) | 5.1 | 1.8 | 2.8(+) |
| Orf8 | 12.1 | 6.4 | 1.9(+) | 11.3 | 6.2 | 1.8(+) | 8.7 | 6.1 | 1.4(+) |
| Orf10 | 1.9 | 2.2 | 0.9(-) | 2.0 | 2.0 | 1.0 | 2.2 | 1.8 | 1.2(+) |
| Nsp1 | 0.5 | 4.5 | 0.1(-) | 0.6 | 5.1 | 0.1(-) | 0.6 | 4.7 | 0.1(-) |
| Nsp2 | 1.3 | 5.5 | 0.2(-) | 1.3 | 5.4 | 0.2(-) | 1.3 | 4.1 | 0.3(-) |
| Nsp3 | 1.9 | 5.9 | 0.3(-) | 2.0 | 6.1 | 0.3(-) | 1.6 | 5.6 | 0.3(-) |
| Nsp4 | 1.7 | 3.3 | 0.5(-) | 2.1 | 3.6 | 0.6(-) | 1.7 | 3.1 | 0.6(-) |
| Nsp5 | 1.0 | 2.3 | 0.4(-) | 1.1 | 2.1 | 0.5(-) | 1.3 | 2.4 | 0.5(-) |
| Nsp6 | 2.4 | 3.7 | 0.7(-) | 2.9 | 4.4 | 0.7(-) | 2.3 | 3.8 | 0.6(-) |
| Nsp7 | 0.6 | 2.6 | 0.2(-) | 0.7 | 2.4 | 0.3(-) | 0.8 | 2.2 | 0.3(-) |
| Nsp8 | 0.6 | 1.3 | 0.4(-) | 0.5 | 1.2 | 0.4(-) | 0.5 | 1.5 | 0.3(-) |
| Nsp9 | 1.0 | 7.7 | 0.1(-) | 0.9 | 9.1 | 0.1(-) | 0.8 | 6.4 | 0.1(-) |
| Nsp10 | 0.4 | 1.5 | 0.2(-) | 0.3 | 1.5 | 0.2(-) | 0.3 | 1.5 | 0.2(-) |
| Nsp11 | 2.4 | 7.9 | 0.3(-) | 2.6 | 7.6 | 0.3(-) | 2.4 | 5.7 | 0.4(-) |
| Nsp12 | 1.4 | 1.7 | 0.8(-) | 1.6 | 1.8 | 0.9(-) | 1.3 | 1.8 | 0.7(-) |
| Nsp13 | 0.8 | 2.9 | 0.3(-) | 0.9 | 2.8 | 0.3(-) | 0.9 | 3.0 | 0.3(-) |
| Nsp14 | 0.7 | 2.8 | 0.2(-) | 0.7 | 3.0 | 0.2(-) | 0.6 | 2.8 | 0.2(-) |
| Nsp15 | 0.6 | 2.3 | 0.3(-) | 0.6 | 2.1 | 0.3(-) | 0.8 | 2.2 | 0.3(-) |

**B).** Li-Wu-Luo (LWL) method.

| | A1a | | | A1b | | | A1c | | |
|---|---|---|---|---|---|---|---|---|---|
| Gene | Ka | Ks | Ka/Ks | Ka | Ks | Ka/Ks | Ka | Ks | Ka/Ks |
| All-TR | 3.0 | 4.8 | 0.6(-) | 3.2 | 4.8 | 0.7(-) | 2.6 | 4.3 | 0.6(-) |
| E | 1.7 | 1.0 | 1.8(+) | 1.7 | 1.0 | 1.7(+) | 1.3 | 1.0 | 1.3(+) |
| S | 7.2 | 2.4 | 3.0(+) | 7.7 | 2.4 | 3.2(+) | 5.8 | 2.3 | 2.5(+) |
| M | 1.8 | 4.5 | 0.4(-) | 2.2 | 4.6 | 0.5(-) | 1.6 | 5.7 | 0.3(-) |
| N | 13.6 | 13.4 | 1.0 | 14.3 | 12.9 | 1.1(+) | 11.4 | 11.2 | 1.0 |
| Orf1ab | 1.5 | 4.9 | 0.3(-) | 1.6 | 5.0 | 0.3(-) | 1.4 | 4.3 | 0.3(-) |
| Orf3a | 0.5 | 4.8 | 0.1(-) | 0.6 | 5.4 | 0.1(-) | 0.6 | 5.0 | 0.1(-) |
| Orf6 | 1.2 | 6.0 | 0.2(-) | 1.2 | 5.9 | 0.2(-) | 1.3 | 4.5 | 0.3(-) |
| Orf7a | 1.9 | 6.4 | 0.3(-) | 2.0 | 6.6 | 0.3(-) | 1.5 | 6.1 | 0.3(-) |
| Orf8 | 1.7 | 3.5 | 0.5(-) | 2.1 | 3.9 | 0.5(-) | 1.7 | 3.4 | 0.5(-) |
| Orf10 | 1.0 | 2.5 | 0.4(-) | 1.1 | 2.3 | 0.5(-) | 1.2 | 2.5 | 0.5(-) |
| Nsp1 | 2.4 | 4.0 | 0.6(-) | 2.8 | 4.7 | 0.6(-) | 2.3 | 4.1 | 0.5(-) |
| Nsp2 | 0.6 | 2.8 | 0.2(-) | 0.7 | 2.6 | 0.3(-) | 0.8 | 2.4 | 0.3(-) |
| Nsp3 | 0.5 | 1.4 | 0.4(-) | 0.5 | 1.3 | 0.4(-) | 0.5 | 1.7 | 0.3(-) |
| Nsp4 | 1.0 | 8.2 | 0.1(-) | 0.9 | 9.7 | 0.1(-) | 0.7 | 6.9 | 0.1(-) |
| Nsp5 | 0.4 | 1.6 | 0.2(-) | 0.3 | 1.6 | 0.2(-) | 0.3 | 1.6 | 0.2(-) |
| Nsp6 | 2.3 | 8.5 | 0.3(-) | 2.5 | 8.2 | 0.3(-) | 2.3 | 6.1 | 0.4(-) |
| Nsp7 | 1.4 | 1.8 | 0.8(-) | 1.6 | 2.0 | 0.8(-) | 1.3 | 1.9 | 0.7(-) |
| Nsp8 | 0.8 | 3.1 | 0.3(-) | 0.9 | 3.0 | 0.3(-) | 0.9 | 3.2 | 0.3(-) |
| Nsp9 | 0.6 | 3.0 | 0.2(-) | 0.7 | 3.3 | 0.2(-) | 0.6 | 3.0 | 0.2(-) |
| Nsp10 | 0.6 | 2.5 | 0.2(-) | 0.6 | 2.3 | 0.3(-) | 0.7 | 2.4 | 0.3(-) |
| Nsp11 | 4.7 | 2.5 | 1.9(+) | 5.0 | 2.4 | 2.1(+) | 5.0 | 2.4 | 2.1(+) |
| Nsp12 | 0.8 | 3.5 | 0.2(-) | 1.0 | 2.9 | 0.3(-) | 1.2 | 3.0 | 0.4(-) |
| Nsp13 | 5.3 | 1.6 | 3.2(+) | 7.2 | 2.1 | 3.4(+) | 5.0 | 2.0 | 2.5(+) |
| Nsp14 | 11.7 | 7.2 | 1.6(+) | 11.0 | 6.9 | 1.6(+) | 8.5 | 6.8 | 1.3(+) |
| Nsp15 | 1.9 | 2.4 | 0.8(-) | 2.0 | 2.1 | 0.9(-) | 2.1 | 2.0 | 1.1(+) |

**C).** Pamilo-Bianchi-Li (PBL) method.

| Gene | A1a | | | A1b | | | A1c | | |
|---|---|---|---|---|---|---|---|---|---|
| | Ka | Ks | Ka/Ks | Ka | Ks | Ka/Ks | Ka | Ks | Ka/Ks |
| All-TR | 3.3 | 3.3 | 1.0 | 3.5 | 3.3 | 1.1(+) | 2.9 | 3.0 | 1.0 |
| E | 2.0 | 0.7 | 2.9(+) | 2.0 | 0.7 | 2.6(+) | 1.5 | 0.8 | 1.8(+) |
| S | 7.6 | 1.8 | 4.2(+) | 8.1 | 1.7 | 4.7(+) | 6.2 | 1.7 | 3.7(+) |
| M | 2.1 | 3.3 | 0.6(-) | 2.6 | 3.4 | 0.8(-) | 1.9 | 4.5 | 0.4(-) |
| N | 14.0 | 10.6 | 1.3(+) | 14.7 | 10.1 | 1.4(+) | 11.9 | 8.6 | 1.4(+) |
| Orf1ab | 1.7 | 3.2 | 0.5(-) | 1.8 | 3.3 | 0.6(-) | 1.6 | 2.9 | 0.6(-) |
| Orf3a | 0.6 | 3.4 | 0.2(-) | 0.6 | 3.8 | 0.2(-) | 0.6 | 3.5 | 0.2(-) |
| Orf6 | 1.4 | 4.1 | 0.3(-) | 1.4 | 4.1 | 0.4(-) | 1.5 | 3.1 | 0.5(-) |
| Orf7a | 2.1 | 4.1 | 0.5(-) | 2.2 | 4.2 | 0.5(-) | 1.7 | 3.8 | 0.4(-) |
| Orf8 | 1.9 | 2.3 | 0.8(-) | 2.4 | 2.6 | 0.9(-) | 1.9 | 2.2 | 0.9(-) |
| Orf10 | 1.2 | 1.8 | 0.7(-) | 1.2 | 1.7 | 0.7(-) | 1.5 | 1.8 | 0.8(-) |
| Nsp1 | 2.7 | 2.8 | 1.0 | 3.2 | 3.2 | 1.0 | 2.5 | 2.7 | 0.9(-) |
| Nsp2 | 0.7 | 2.2 | 0.3(-) | 0.8 | 2.0 | 0.4(-) | 0.9 | 1.7 | 0.5(-) |
| Nsp3 | 0.6 | 0.9 | 0.7(-) | 0.6 | 0.9 | 0.7(-) | 0.6 | 1.2 | 0.5(-) |
| Nsp4 | 1.2 | 5.2 | 0.2(-) | 1.1 | 6.1 | 0.2(-) | 0.9 | 4.4 | 0.2(-) |
| Nsp5 | 0.4 | 1.5 | 0.3(-) | 0.4 | 1.6 | 0.2(-) | 0.3 | 1.5 | 0.2(-) |
| Nsp6 | 2.8 | 5.1 | 0.6(-) | 3.0 | 4.9 | 0.6(-) | 2.8 | 3.7 | 0.8(-) |
| Nsp7 | 1.5 | 1.4 | 1.1(+) | 1.7 | 1.5 | 1.2(+) | 1.4 | 1.6 | 0.9(-) |
| Nsp8 | 1.0 | 2.1 | 0.5(-) | 1.1 | 2.0 | 0.5(-) | 1.0 | 2.1 | 0.5(-) |
| Nsp9 | 0.7 | 1.9 | 0.4(-) | 0.8 | 2.0 | 0.4(-) | 0.7 | 2.0 | 0.3(-) |
| Nsp10 | 0.7 | 2.0 | 0.3(-) | 0.7 | 1.9 | 0.4(-) | 0.8 | 2.3 | 0.4(-) |
| Nsp11 | 5.0 | 1.9 | 2.7(+) | 5.4 | 1.7 | 3.1(+) | 5.3 | 1.8 | 2.9(+) |
| Nsp12 | 0.9 | 1.8 | 0.5(-) | 1.1 | 1.5 | 0.7(-) | 1.4 | 1.6 | 0.9(-) |
| Nsp13 | 6.3 | 1.3 | 5.0(+) | 8.6 | 1.7 | 5.1(+) | 5.9 | 1.6 | 3.7(+) |
| Nsp14 | 12.8 | 4.7 | 2.7(+) | 12.0 | 4.5 | 2.6(+) | 9.4 | 4.7 | 2.0(+) |
| Nsp15 | 2.0 | 1.6 | 1.3(+) | 2.1 | 1.4 | 1.6(+) | 2.2 | 1.2 | 1.8(+) |

**D).** Maximum-Likelihood (ML) method.

| Gene | A1a | | | A1b | | | A1c | | |
|---|---|---|---|---|---|---|---|---|---|
| | Ka | Ks | Ka/Ks | Ka | Ks | Ka/Ks | Ka | Ks | Ka/Ks |
| All-TR | 3.1 | 4.7 | 0.7(-) | 3.3 | 4.8 | 0.7(-) | 2.7 | 4.2 | 0.7(-) |
| E | 1.9 | 0.8 | 2.4(+) | 1.9 | 0.8 | 2.3(+) | 1.4 | 0.9 | 1.6(+) |
| S | 7.2 | 2.6 | 2.7(+) | 7.7 | 2.6 | 3.0(+) | 5.8 | 2.4 | 2.4(+) |
| M | 2.0 | 3.4 | 0.6(-) | 2.6 | 3.5 | 0.7(-) | 1.9 | 4.5 | 0.4(-) |
| N | 14.5 | 11.2 | 1.3(+) | 15.1 | 10.9 | 1.4(+) | 12.1 | 9.5 | 1.3(+) |
| Orf1ab | 1.6 | 4.5 | 0.4(-) | 1.7 | 4.5 | 0.4(-) | 1.5 | 4.0 | 0.4(-) |
| Orf3a | 0.6 | 3.2 | 0.2(-) | 0.6 | 3.6 | 0.2(-) | 0.7 | 3.4 | 0.2(-) |
| Orf6 | 1.4 | 4.9 | 0.3(-) | 1.4 | 4.8 | 0.3(-) | 1.4 | 3.7 | 0.4(-) |
| Orf7a | 1.9 | 6.2 | 0.3(-) | 2.1 | 6.4 | 0.3(-) | 1.6 | 5.8 | 0.3(-) |
| Orf8 | 1.8 | 3.5 | 0.5(-) | 2.2 | 3.9 | 0.6(-) | 1.7 | 3.3 | 0.5(-) |
| Orf10 | 1.0 | 2.6 | 0.4(-) | 1.1 | 2.4 | 0.5(-) | 1.3 | 2.6 | 0.5(-) |
| Nsp1 | 2.6 | 3.4 | 0.8(-) | 3.1 | 4.0 | 0.8(-) | 2.4 | 3.4 | 0.7(-) |
| Nsp2 | 0.7 | 2.2 | 0.3(-) | 0.8 | 2.0 | 0.4(-) | 0.9 | 1.8 | 0.5(-) |
| Nsp3 | 0.6 | 1.2 | 0.5(-) | 0.5 | 1.1 | 0.5(-) | 0.5 | 1.5 | 0.4(-) |
| Nsp4 | 1.0 | 7.6 | 0.1(-) | 1.0 | 8.9 | 0.1(-) | 0.8 | 6.4 | 0.1(-) |
| Nsp5 | 0.4 | 1.8 | 0.2(-) | 0.3 | 1.9 | 0.2(-) | 0.3 | 1.8 | 0.2(-) |
| Nsp6 | 2.6 | 6.3 | 0.4(-) | 2.8 | 6.1 | 0.5(-) | 2.6 | 4.5 | 0.6(-) |
| Nsp7 | 1.4 | 1.9 | 0.7(-) | 1.6 | 2.0 | 0.8(-) | 1.3 | 2.1 | 0.6(-) |
| Nsp8 | 0.9 | 2.8 | 0.3(-) | 1.0 | 2.7 | 0.4(-) | 1.0 | 2.8 | 0.3(-) |
| Nsp9 | 0.7 | 2.9 | 0.2(-) | 0.7 | 3.2 | 0.2(-) | 0.6 | 3.0 | 0.2(-) |
| Nsp10 | 0.6 | 2.9 | 0.2(-) | 0.6 | 2.7 | 0.2(-) | 0.7 | 3.0 | 0.2(-) |
| Nsp11 | 5.0 | 2.2 | 2.3(+) | 5.4 | 2.0 | 2.7(+) | 5.2 | 2.2 | 2.4(+) |
| Nsp12 | 0.9 | 2.3 | 0.4(-) | 1.1 | 1.9 | 0.6(-) | 1.4 | 1.9 | 0.7(-) |
| Nsp13 | 5.6 | 1.4 | 3.9(+) | 7.7 | 1.9 | 4.1(+) | 5.3 | 1.8 | 3.0(+) |
| Nsp14 | 12.3 | 6.2 | 2.0(+) | 11.5 | 6.0 | 1.9(+) | 9.0 | 6.0 | 1.5(+) |
| Nsp15 | 2.0 | 1.8 | 1.1(+) | 2.1 | 1.6 | 1.4(+) | 2.2 | 1.4 | 1.6(+) |

**Table S10.** Decomposition of each codon/NT sequence into different selection type using Ka/Ks values. Two categories (**A-B**): positive selection (P) and negative selection (N). Five categories (**C-D**): conserved percentage (%C), strong negative percentage (%SN), weak negative percentage (%WN), weak positive percentage (%WP) and strong positive percentage (%SP).

#### A). All-TR

| PRO | All-TR | Orf1ab | Non-Orf1ab | S | Orf3a | E | M | Orf6 | Orf7a | Orf8 | N | Orf10 |
|---|---|---|---|---|---|---|---|---|---|---|---|---|
| %P/N, NG | 23/77 | 15/85 | 45/55 | 50/50 | **58/42** | 29/71 | 19/81 | 0/100 | 30/70 | **88/12** | 44/56 | 3/97 |
| %P/N, LWL | 22/78 | 14/86 | 44/56 | 50/50 | **55/45** | 26/74 | 17/83 | 0/100 | 30/70 | **88/12** | 36/64 | 3/97 |
| %P/N, PBL | 28/72 | 19/81 | **52/48** | **53/47** | **82/18** | 37/63 | 29/71 | 19/81 | 45/55 | **93/7** | 42/58 | 10/90 |
| %P/N, Avg | 24/76 | 16/84 | 47/53 | **51/49** | **65/35** | 31/69 | 22/78 | 6/94 | 35/65 | **90/10** | 41/59 | 5/95 |
| Sel (+/-) | - | - | - | + | + | - | - | - | - | + | - | - |

*With reference to the AA codon substitution.

#### B). Nsp 1-15

| PRO | Nsp1 | Nsp2 | Nsp3 | Nsp4 | Nsp5 | Nsp6 | Nsp7 | Nsp8 | Nsp9 | Nsp10 | Nsp11 | Nsp12 | Nsp13 | Nsp14 | Nsp15 |
|---|---|---|---|---|---|---|---|---|---|---|---|---|---|---|---|
| %P/N, NG | 0/100 | 17/83 | 25/75 | 18/82 | 30/70 | 43/57 | 0/100 | 17/83 | 0/100 | 0/100 | 17/83 | 27/73 | 14/86 | 26/74 | 7/93 |
| %P/N, LWL | 0/100 | 15/85 | 23/77 | 14/86 | 29/71 | 42/58 | 0/100 | 12/88 | 0/100 | 0/100 | 16/84 | 22/78 | 12/88 | 21/79 | 7/93 |
| %P/N, PBL | 0/100 | 24/76 | 30/70 | 28/72 | 34/66 | 49/51 | 0/100 | 21/79 | 0/100 | 0/100 | 23/77 | 36/64 | 18/82 | 32/68 | 10/90 |
| %P/N, Avg | 0/100 | 19/81 | 26/74 | 20/80 | 31/69 | 45/55 | 0/100 | 16/84 | 0/100 | 0/100 | 19/81 | 28/72 | 15/85 | 26/74 | 8/92 |
| Sel (+/-) | - | - | - | - | - | - | - | - | - | - | - | - | - | - | - |

A&B) **%N:** Ka=Ks=0 **or** 0 < Ka/Ks < 1.0; **%P:** 1.0 < Ka/Ks

#### C). All-TR

| PRO | All-TR | Orf1ab | Non-Orf1ab | S | Orf3a | E | M | Orf6 | Orf7a | Orf8 | N |
|---|---|---|---|---|---|---|---|---|---|---|---|
| %C | 22 | 23 | 17 | 22 | 8 | 32 | 24 | 11 | 7 | 4 | 11 |
| %SN | 42 | 49 | 23 | 20 | 3 | 16 | 44 | 68 | 16 | 2 | 36 |
| %WN | 13 | 13 | 14 | 8 | 32 | 24 | 13 | 21 | 48 | 6 | 10 |
| %WP | 8 | 7 | 10 | 10 | 8 | 4 | 9 | 0 | 0 | 0 | 22 |
| %SP | 15 | 8 | 35 | 40 | 50 | 25 | 10 | 0 | 30 | 88 | 21 |
| Sel(+/-) | - | - | + | + | + | - | - | - | - | + | - |

#### D). Nsp 1-15

| PRO | Nsp1 | Nsp2 | Nsp3 | Nsp4 | Nsp5 | Nsp6 | Nsp7 | Nsp8 | Nsp9 | Nsp10 | Nsp11 | Nsp12 | Nsp13 | Nsp14 | Nsp15 |
|---|---|---|---|---|---|---|---|---|---|---|---|---|---|---|---|
| %C | 13 | 11 | 21 | 24 | 26 | 21 | 22 | 29 | 23 | 30 | 31 | 29 | 25 | 23 | 29 |
| %SN | 77 | 59 | 41 | 43 | 47 | 39 | 78 | 50 | 68 | 70 | 52 | 40 | 52 | 49 | 53 |
| %WN | 11 | 15 | 18 | 19 | 6 | 6 | 0 | 10 | 9 | 0 | 7 | 14 | 13 | 7 | 14 |
| %WP | 0 | 8 | 6 | 8 | 12 | 10 | 0 | 12 | 0 | 0 | 4 | 10 | 8 | 16 | 4 |
| %SP | 0 | 7 | 14 | 6 | 8 | 25 | 0 | 0 | 0 | 0 | 7 | 7 | 2 | 3 | 0 |
| Sel(+/-) | - | - | - | - | - | - | - | - | - | - | - | - | - | - | - |

C&D) **%C:** Ka/Ks = 0; **%SN**: 0 < Ka/Ks <0.5; **%WN:** 0.5 < Ka/Ks < 1.0; **%WP:** 1.0 < Ka/Ks <2.0; **%SP:** Ka/Ks >2.0

**Table S11**: Percentage of top 247 non-synonymous AA substitutions occurring on the four secondary structure types in each coding protein in All-TR.

| | E | S | M | N | Orf3A | Orf6 | Orf7A | Orf8 | Orf10 | Nsp1 | Nsp2 | Nsp3 | Nsp4 | Nsp5 | Nsp6 | Nsp7 | Nsp8 | Nsp9 | Nsp10 | Nsp11 | Nsp12 | Nsp13 | Nsp14 | Nsp15 |
|---|---|---|---|---|---|---|---|---|---|---|---|---|---|---|---|---|---|---|---|---|---|---|---|---|
| **%A** | 0.00 | 0.13 | **1.00** | **0.36** | **0.27** | **0.50** | 0.20 | 0.00 | **0.50** | 0.00 | 0.29 | 0.31 | **0.33** | 0.25 | **0.80** | **0.50** | **1.00** | 0.00 | 0.00 | 0.00 | 0.25 | 0.00 | **0.67** | **0.33** |
| **%B** | 0.00 | 0.21 | 0.00 | 0.00 | **0.27** | 0.00 | 0.00 | 0.19 | **0.50** | 0.00 | 0.29 | 0.08 | 0.22 | **0.75** | 0.00 | 0.00 | 0.00 | **0.50** | 0.00 | 0.00 | 0.13 | 0.00 | 0.00 | 0.00 |
| **%C** | **1.00** | 0.29 | 0.00 | 0.32 | **0.27** | **0.50** | **0.80** | 0.31 | 0.00 | 0.00 | 0.00 | 0.23 | 0.11 | 0.00 | 0.10 | 0.00 | 0.00 | 0.00 | 0.00 | 0.25 | 0.13 | 0.25 | 0.33 | **0.33** |
| **%T** | 0.00 | **0.37** | 0.00 | 0.25 | 0.18 | 0.00 | 0.00 | **0.50** | 0.00 | 0.00 | **0.43** | **0.38** | **0.33** | 0.00 | 0.10 | **0.50** | 0.00 | **0.50** | 0.00 | **0.75** | **0.50** | **0.75** | 0.00 | **0.33** |

**%A**: Alpha-helices; **%B**: Beta-sheets; **%C**: Random coils; **%T**: Beta-turns

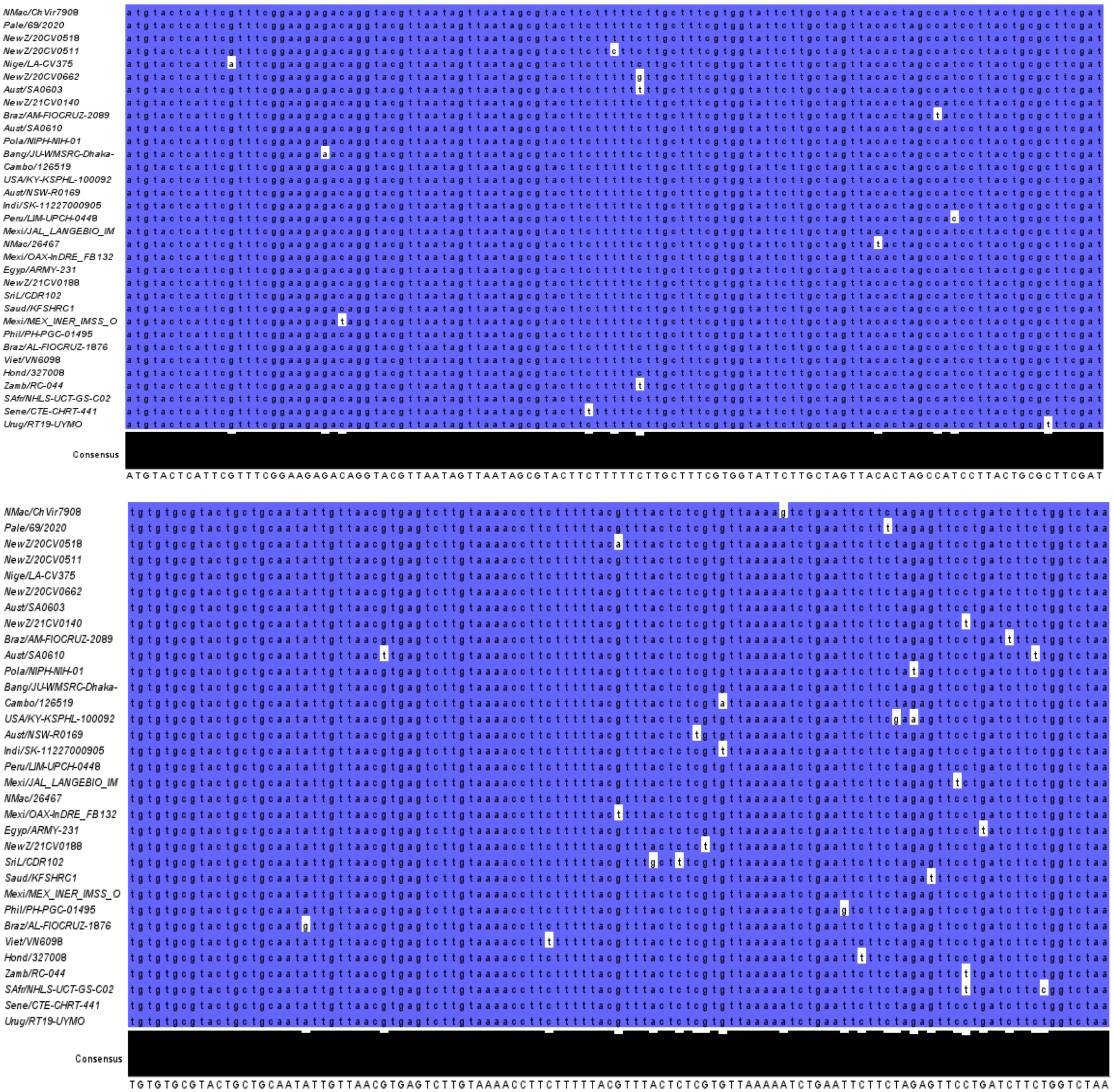

**Figure S2a.** MSA of the selected non-redundant E gene sequences from set A1a against the human reference sequence (Wuhan-Hu-1) using Jalview. The sequences are colored based on “Percentage Identity” using Jalview. With respect to the reference sequence: **Blue:** Conserved; **White**: Non-conserved.

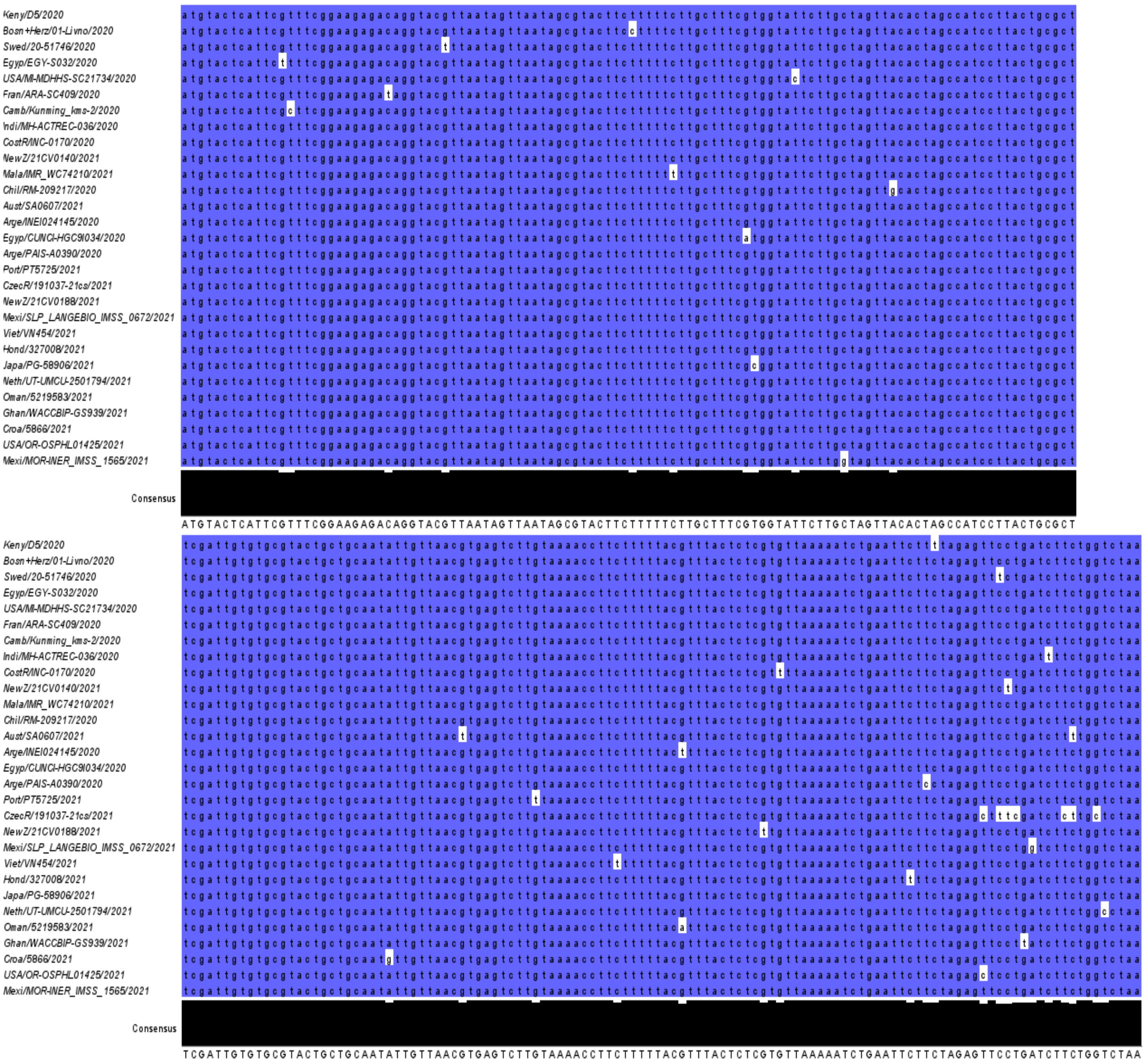


**Figure S2b.** MSA of the selected non-redundant E gene sequences from set A1b against the human reference sequence (Wuhan-Hu-1) using Jalview. The sequences are colored based on “Percentage Identity” using Jalview. With respect to the reference sequence: **Blue:** Conserved; **White**: Non-conserved.

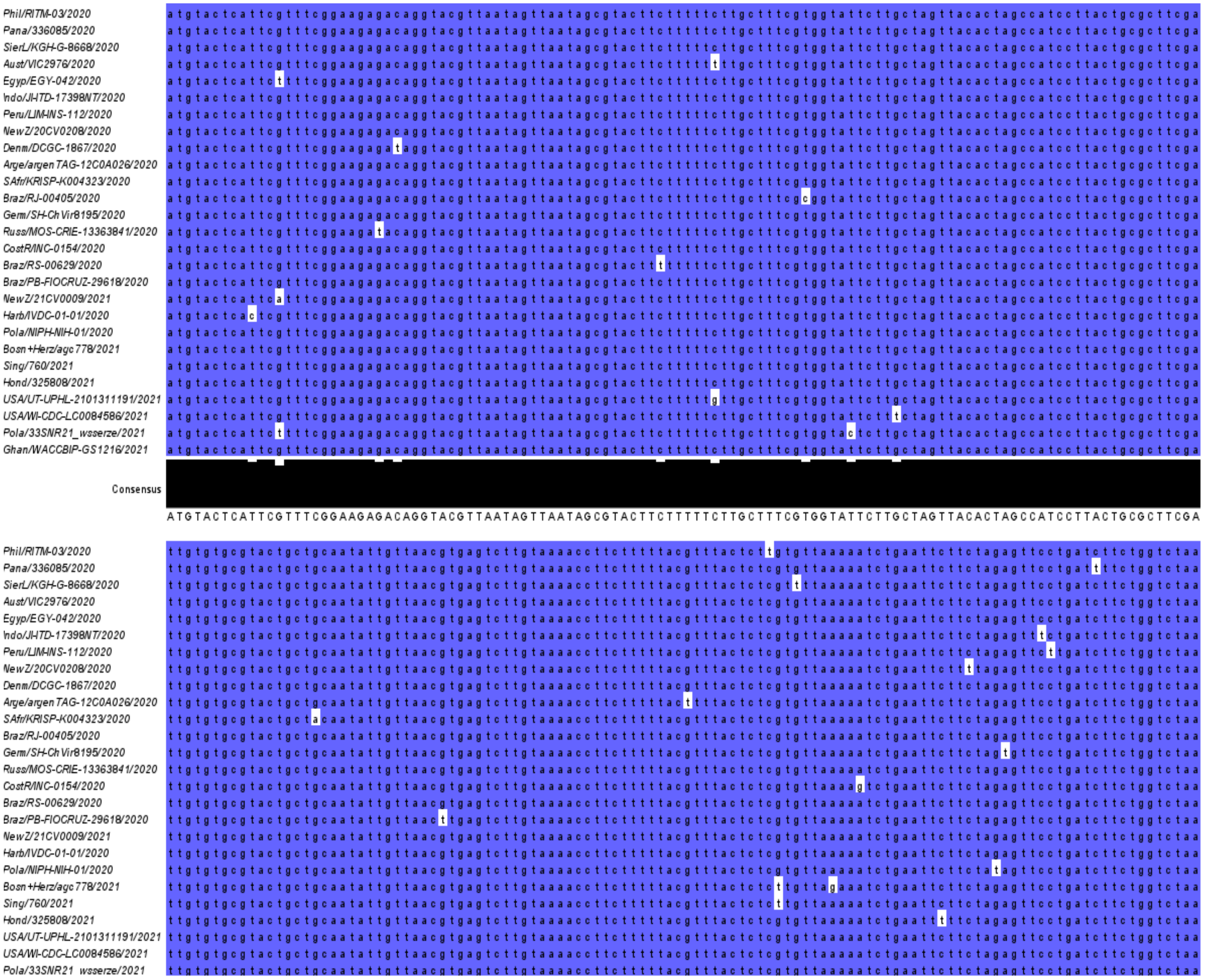

**Figure S2c.** MSA of the selected non-redundant E gene sequences from Set A1c against the human reference sequence (Wuhan-Hu-1) using Jalview. The sequences are colored based on “Percentage Identity” using Jalview. With respect to the reference sequence: **Blue:** Conserved; **White**: Non-conserved.

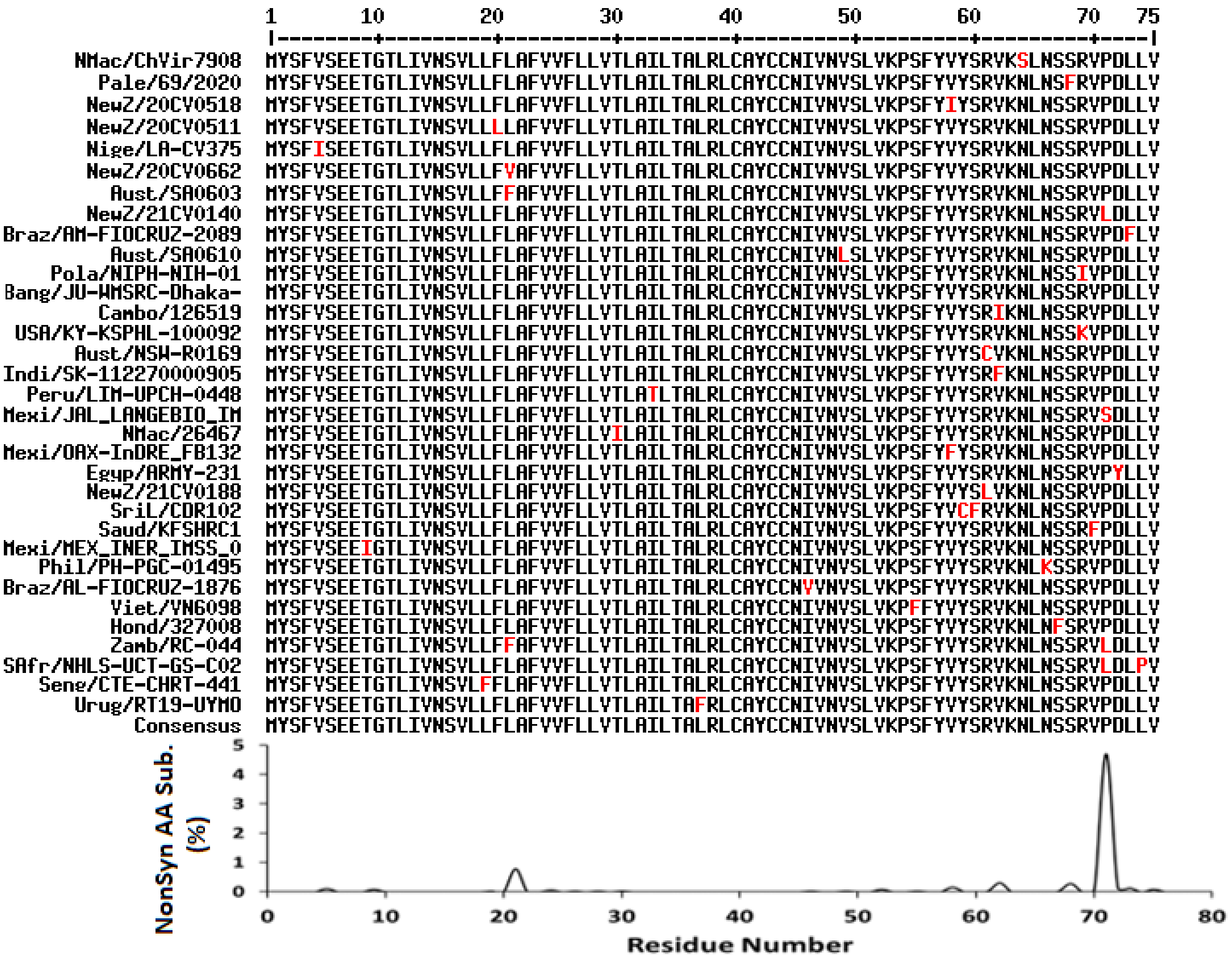

**Figure S3a.** MSA of selected non-redundant E protein sequences from set A1a against the human reference sequence (Wuhan-Hu-1) using the Multialin web server. **Top**: Sequences containing non-synonymous AA substitutions (red). The human reference sequence was removed to highlight the observed substitutions. **Bottom**: Percent non-synonymous AA substitution in each residue within the non-redundant E protein sequences. Percent substitution was calculated against the consensus sequence (bottom of MSA).

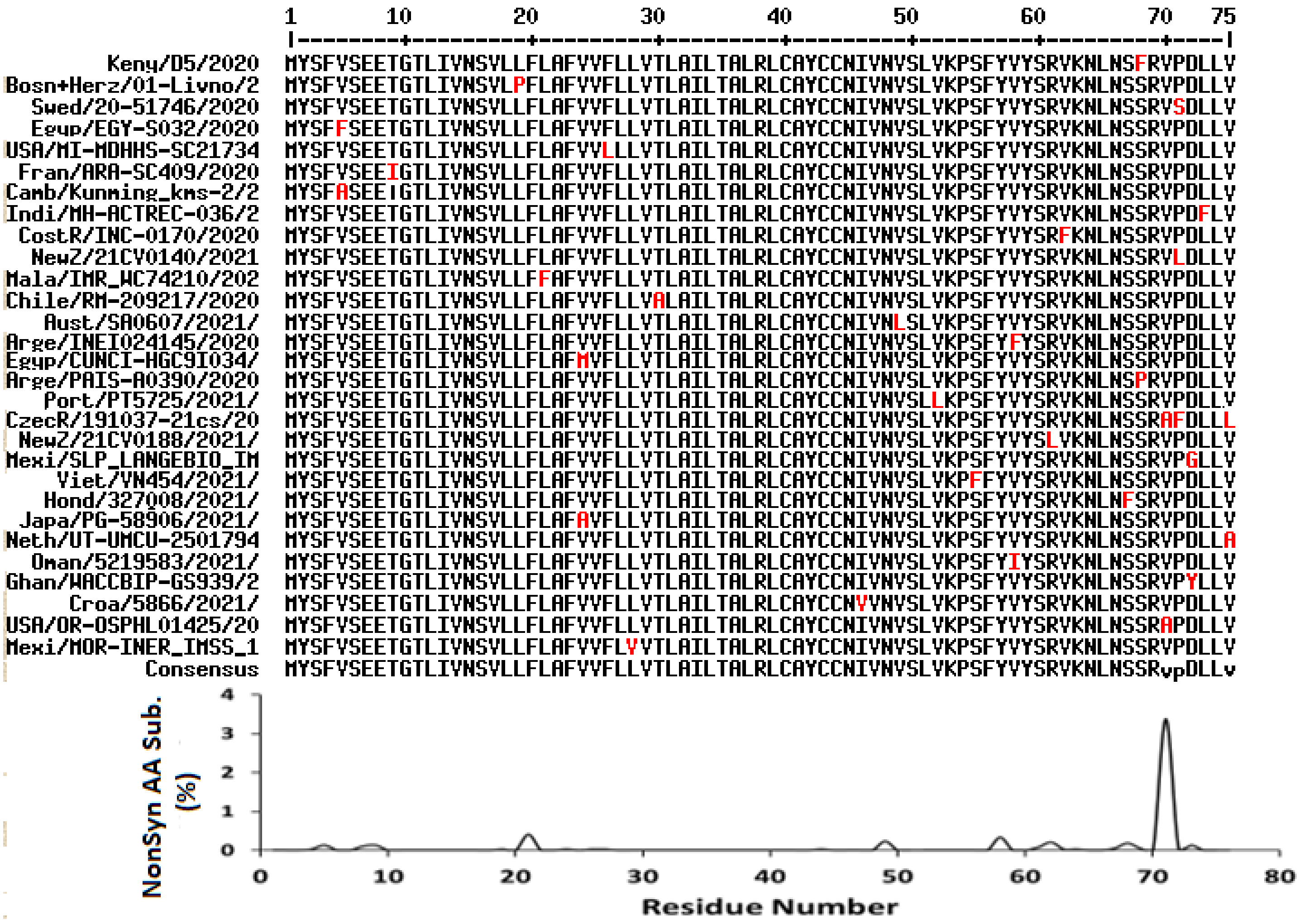


**Figure S3b.** MSA of selected non-redundant E protein sequences from set A1b against the human reference sequence (Wuhan-Hu-1) using the Multialin web server. **Top**: Sequences containing non-synonymous AA substitutions (red). The human reference sequence was removed to highlight the observed substitutions. **Bottom**: Percent non-synonymous AA substitution in each residue within the non-redundant E protein sequences. Percent substitution was calculated against the consensus sequence (bottom of MSA).

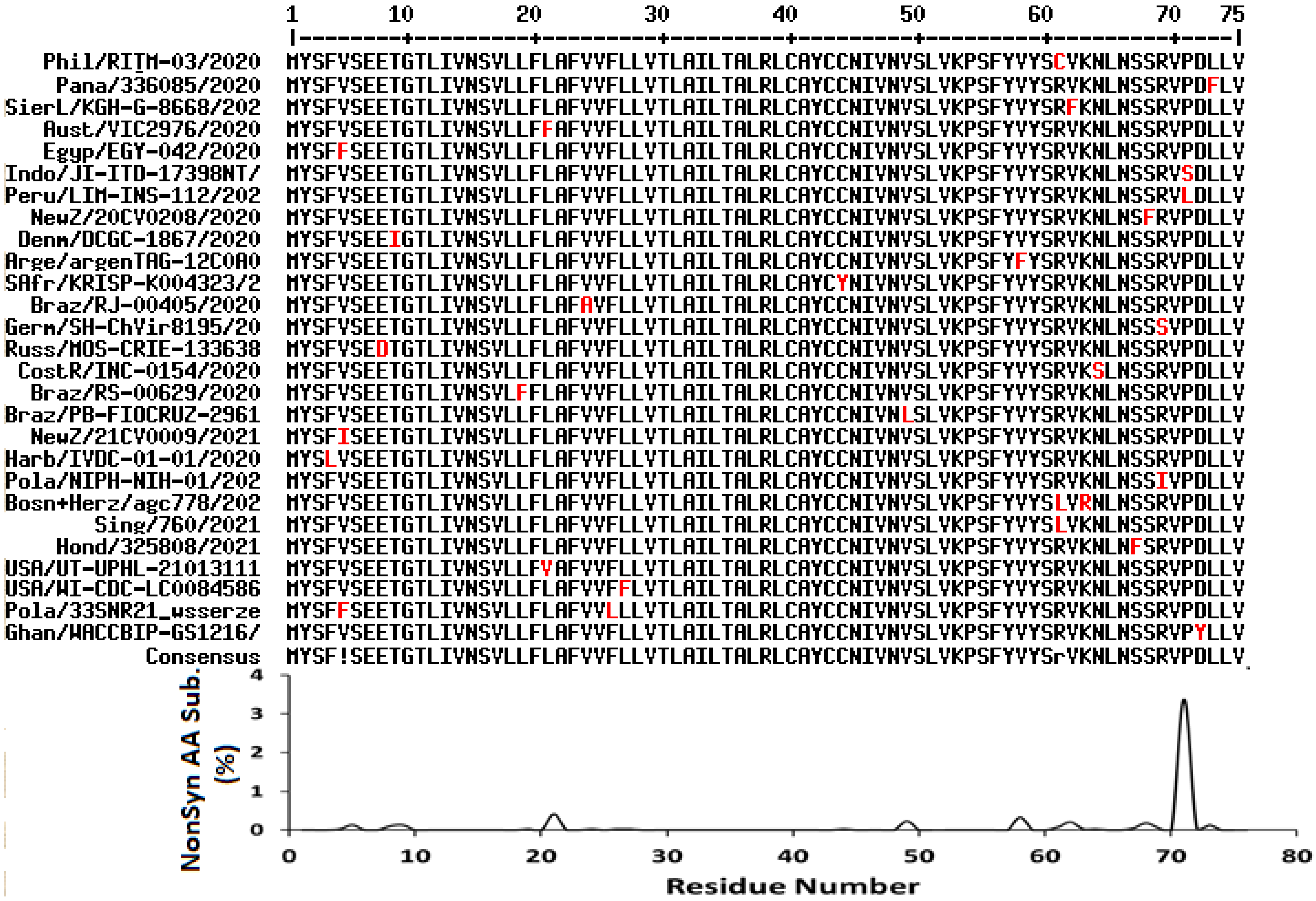

**Figure S3c.** MSA of selected non-redundant E protein sequences from set A1c against the human reference sequence (Wuhan-Hu-1) using the Multialin web server. **Top**: Sequences containing non-synonymous AA substitutions (red). The human reference sequence was removed to highlight the observed substitutions. **Bottom**: Percent non-synonymous AA substitution in each residue within the non-redundant E protein sequences. Percent substitution was calculated against the consensus sequence (bottom of MSA).

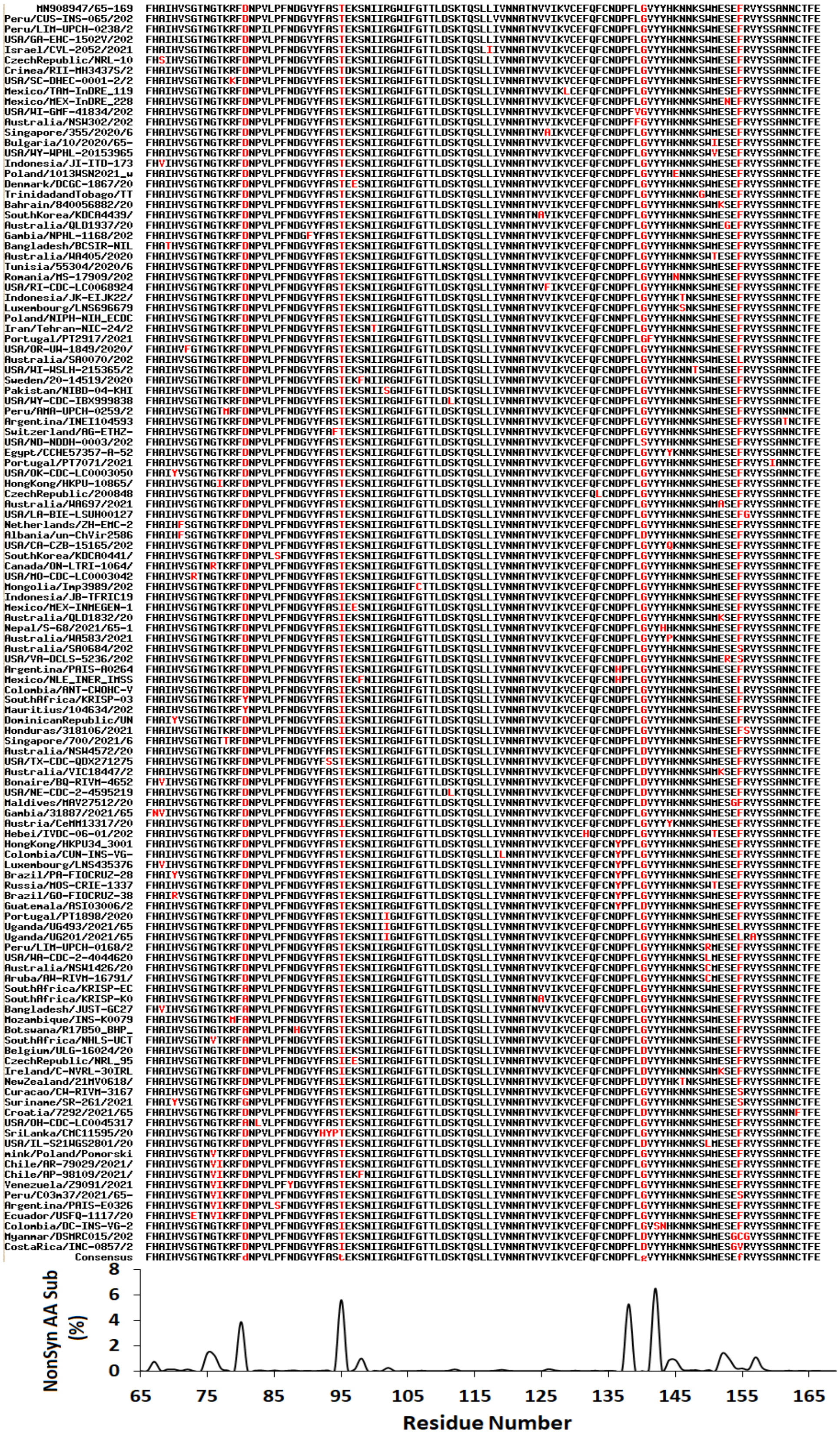


**Figure S4a.** MSA of selected non-redundant S protein sequences (residues 65-169) from set A1c against the human reference sequence (Wuhan-Hu-1) using the Multialin web server. **Top**: Sequences containing non-synonymous AA substitutions (red). The human reference sequence was removed to highlight the observed substitutions. **Bottom**: Percent non-synonymous AA substitution in each residue within the non-redundant E protein sequences. Percent substitution was calculated against the consensus sequence (bottom of MSA).

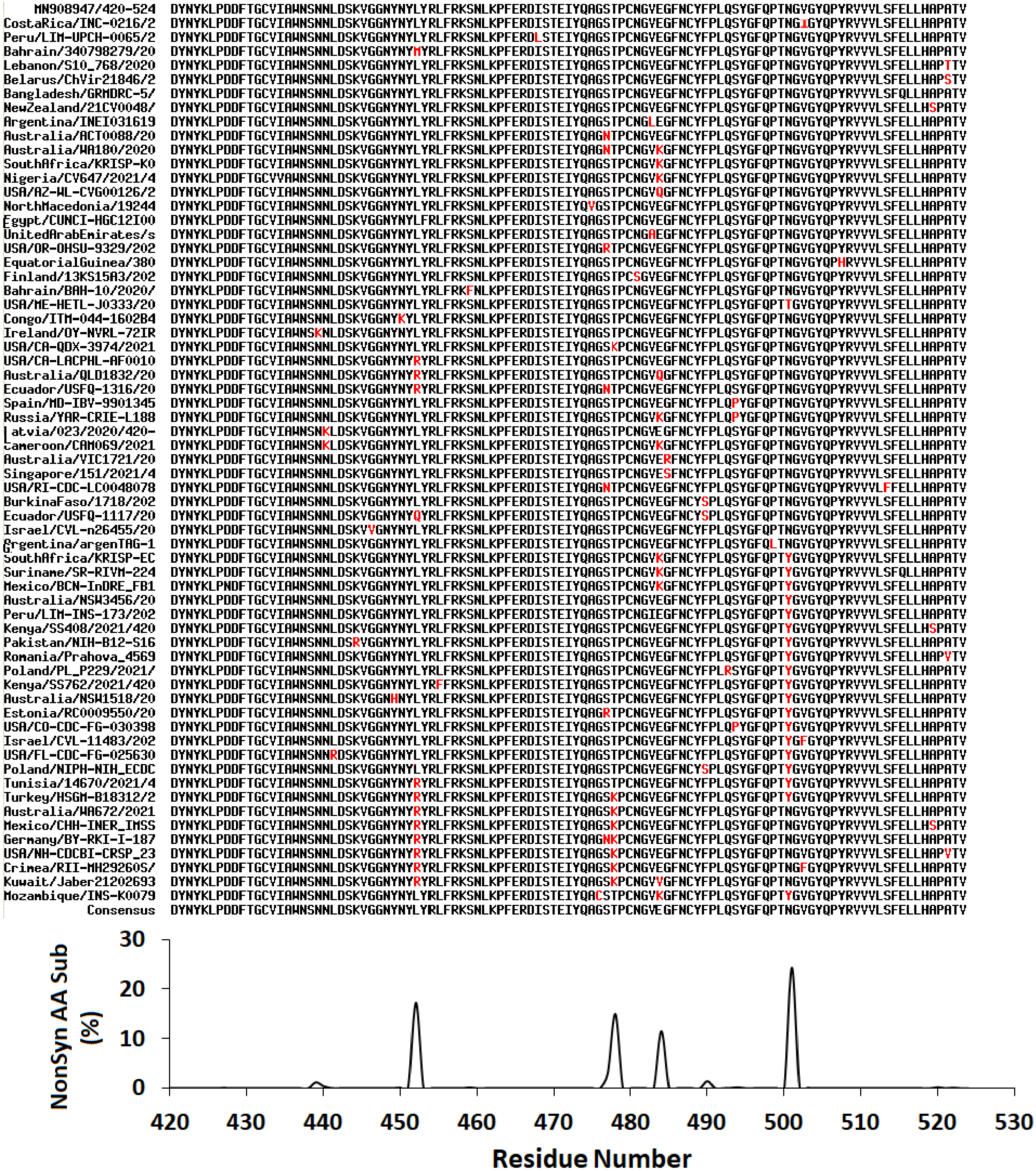


**Figure S4b.** MSA of selected non-redundant S protein sequences (residues 420-524) from set A1c against the human reference sequence (Wuhan-Hu-1) using the Multialin web server. **Top**: Sequences containing non-synonymous AA substitutions (red). The human reference sequence was removed to highlight the observed substitutions. **Bottom**: Percent non-synonymous AA substitution in each residue within the non-redundant E protein sequences. Percent substitution was calculated against the consensus sequence (bottom of MSA).

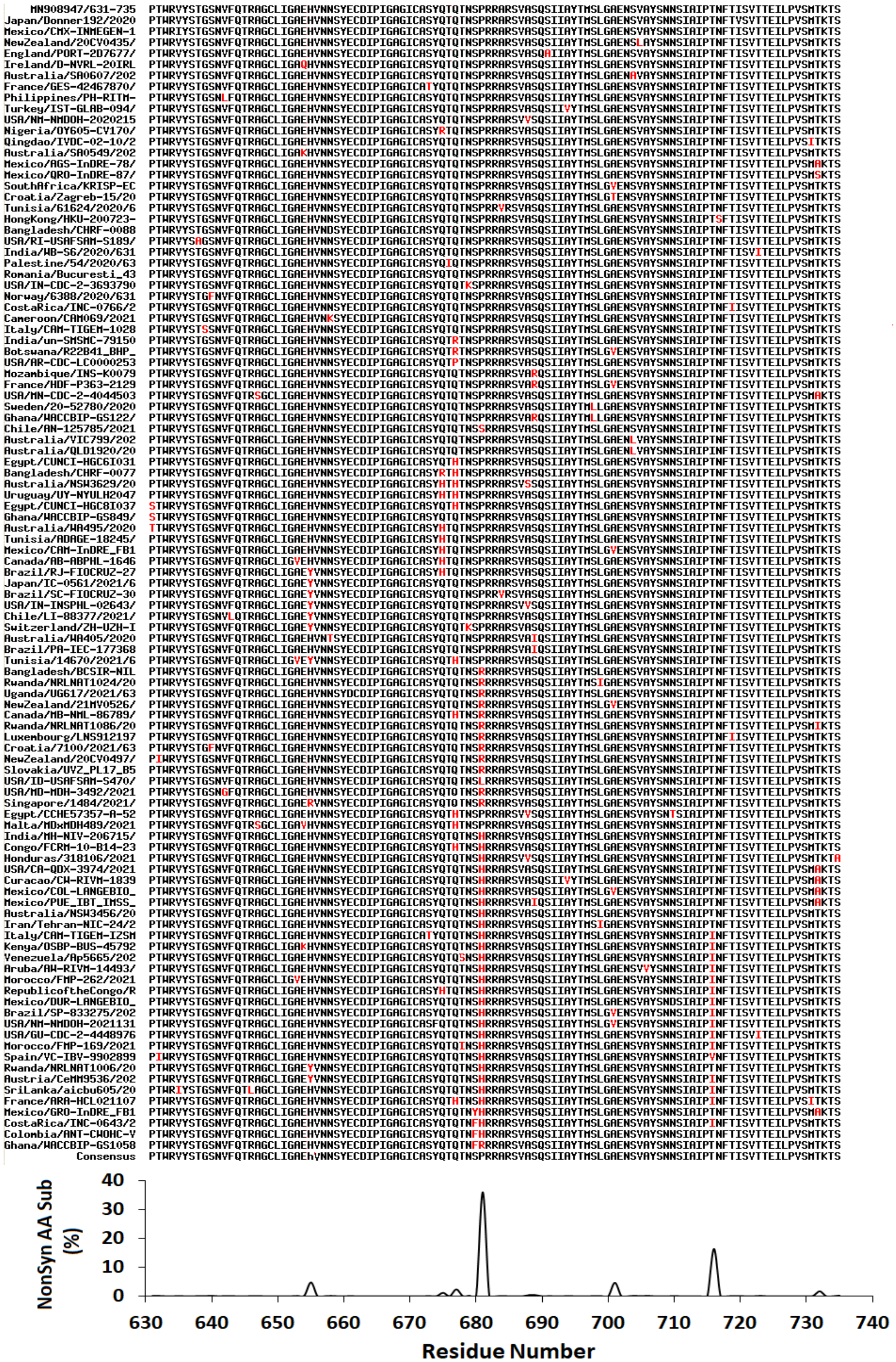


**Figure S4c.** MSA of selected non-redundant S protein sequences (residues 631-735) from set A1c against the human reference sequence (Wuhan-Hu-1) using the Multialin web server. **Top**: Sequences containing non-synonymous AA substitutions (red). The human reference sequence was removed to highlight the observed substitutions. **Bottom**: Percent non-synonymous AA substitution in each residue within the non-redundant E protein sequences. Percent substitution was calculated against the consensus sequence (bottom of MSA).

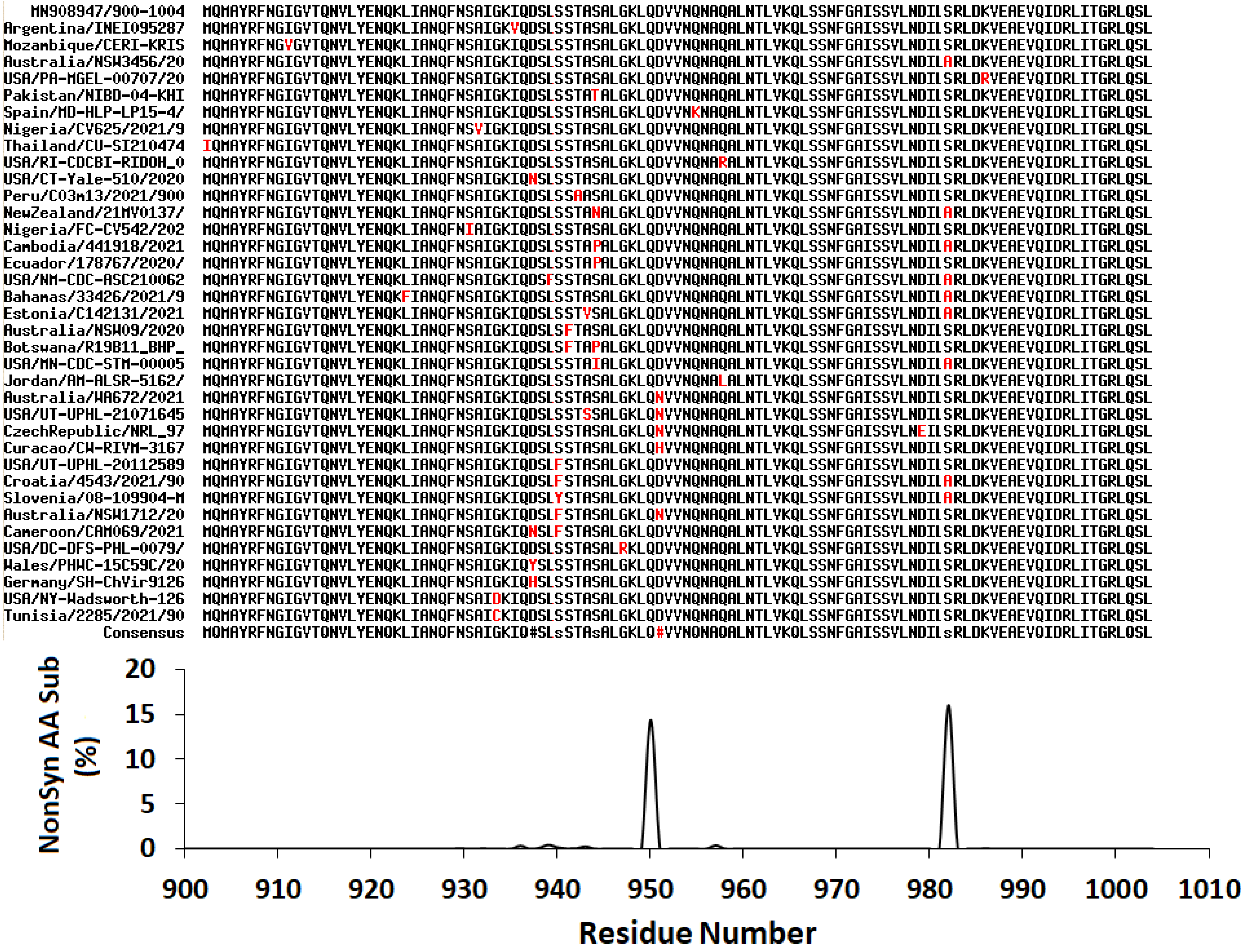


**Figure S4d.** MSA of selected non-redundant S protein sequences (residues 900-1004) from set A1c against the human reference sequence (Wuhan-Hu-1) using the Multialin web server. **Top**: Sequences containing non-synonymous AA substitutions (red). The human reference sequence was removed to highlight the observed substitutions. **Bottom**: Percent non-synonymous AA substitution in each residue within the non-redundant E protein sequences. Percent substitution was calculated against the consensus sequence (bottom of MSA).

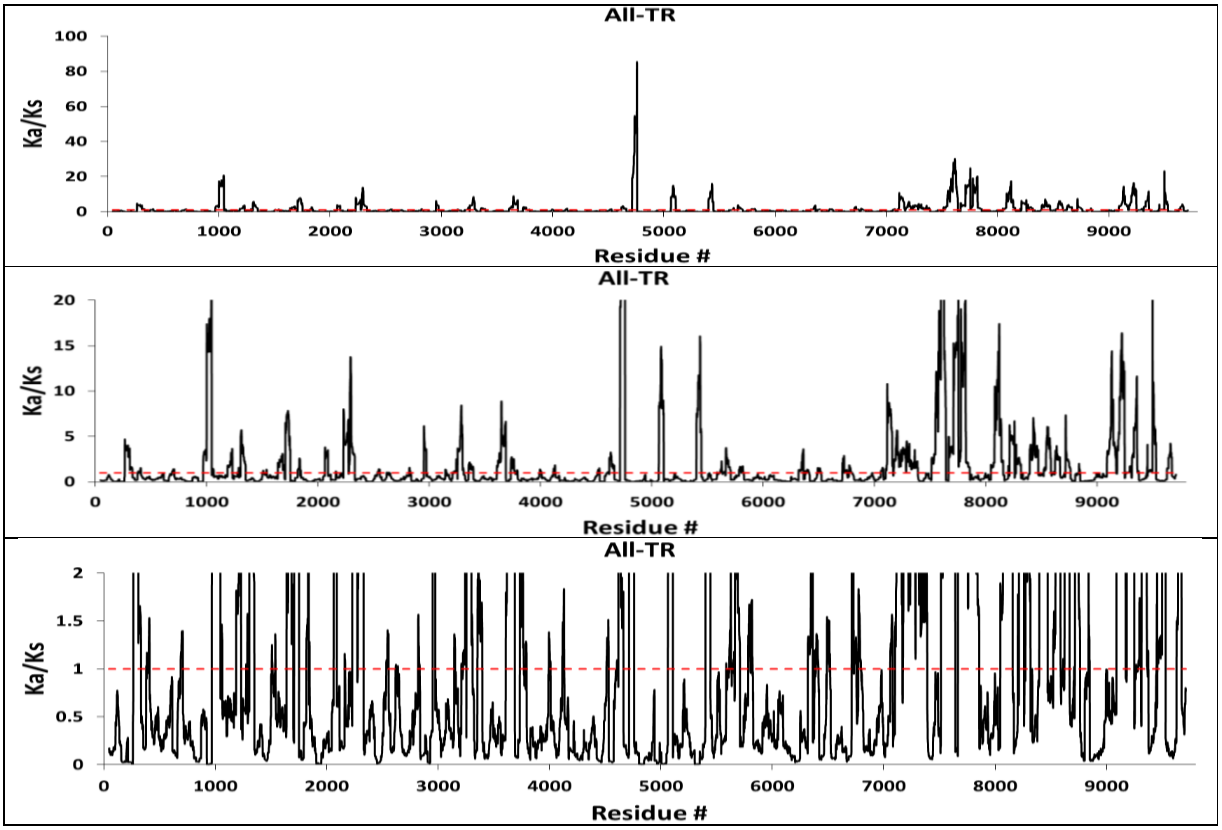


**Figure S5a.** Ka/Ks distribution of each codon in the proteome of each sequence in datasets A1a/A1b/A1c using the NG method with a sliding window of 45 AA residues. The red line represents Ka/Ks = 1.0 for sites likely under neutral selection pressure.

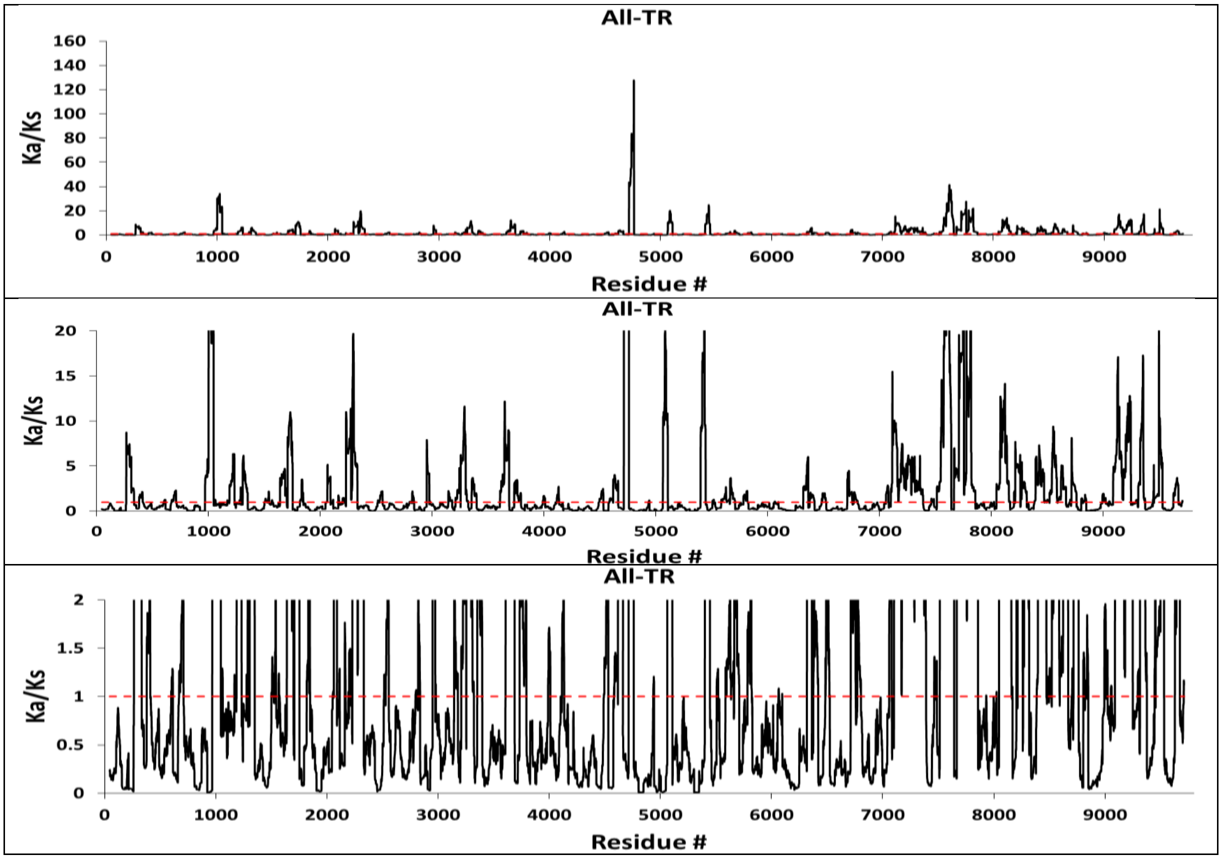

**Figure S5b.** Ka/Ks distribution of each codon in the proteome of each sequence in datasets A1a/A1b/A1c using the PBL method with a sliding window of 45 AA residues. The red line represents Ka/Ks = 1.0 for sites likely under neutral selection pressure.

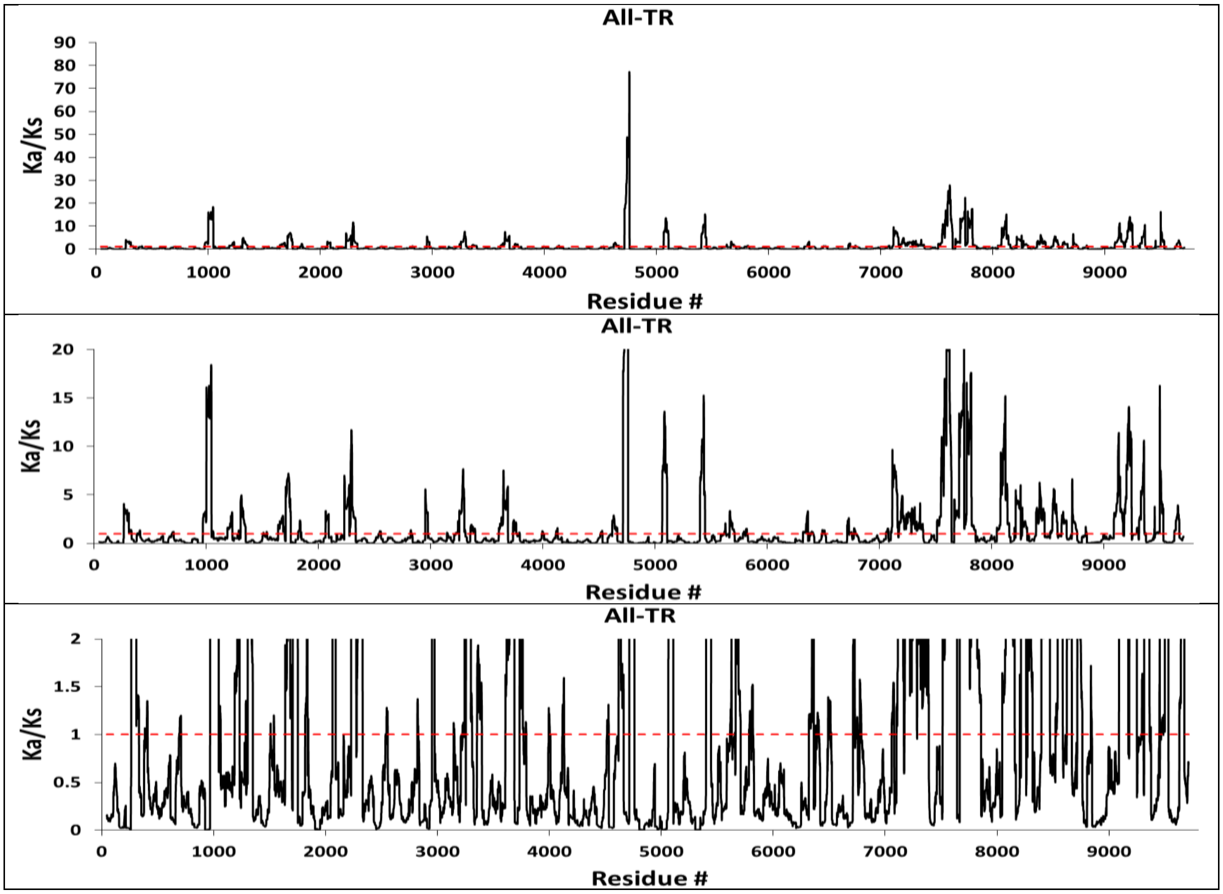


**Figure S5c.** Ka/Ks distribution of each codon in the proteome of each sequence in datasets A1a/A1b/A1c using the LWL method with a sliding window of 45 AA residues. The red line represents Ka/Ks = 1.0 for sites likely under neutral selection pressure.

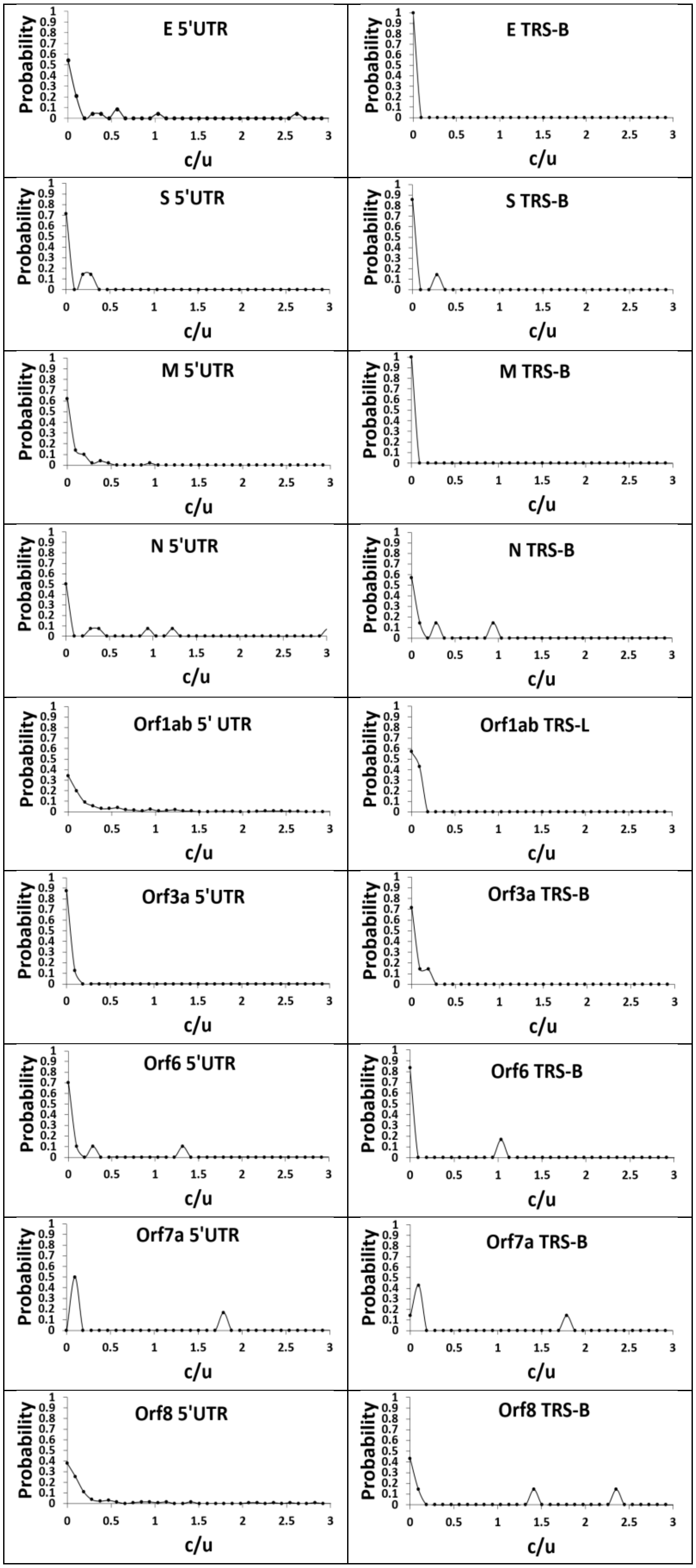
E 5'UTR
E TRS-B
S 5'UTR
S TRS-B
M 5'UTR
M TRS-B
N 5'UTR
N TRS-B
Orf1ab 5' UTR
Orf1ab TRS-L
Orf3a 5'UTR
Orf3a TRS-B
Orf6 5'UTR
Orf6 TRS-B
Orf7a 5'UTR
Orf7a TRS-B
Orf8 5'UTR
Orf8 TRS-B
Probability
c/u

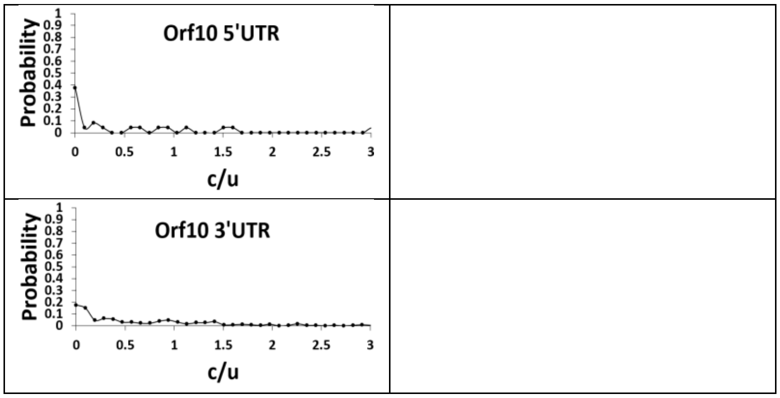


**Figure S6.** Probability distribution of the relative total NT substitution rate (*c/μ*) across each UTR and TRS across datasets A1a/A1b/A1c.

c = number of substitutions
μ = average number of substitutions per site per 19 months: 10.6 for the genome.
*Orf10 does not have a TRS-B, thus it is not shown.

| Total NT Substitutions | Synonymous NT Substitutions | Non-Synonymous NT Substitutions |
|---|---|---|

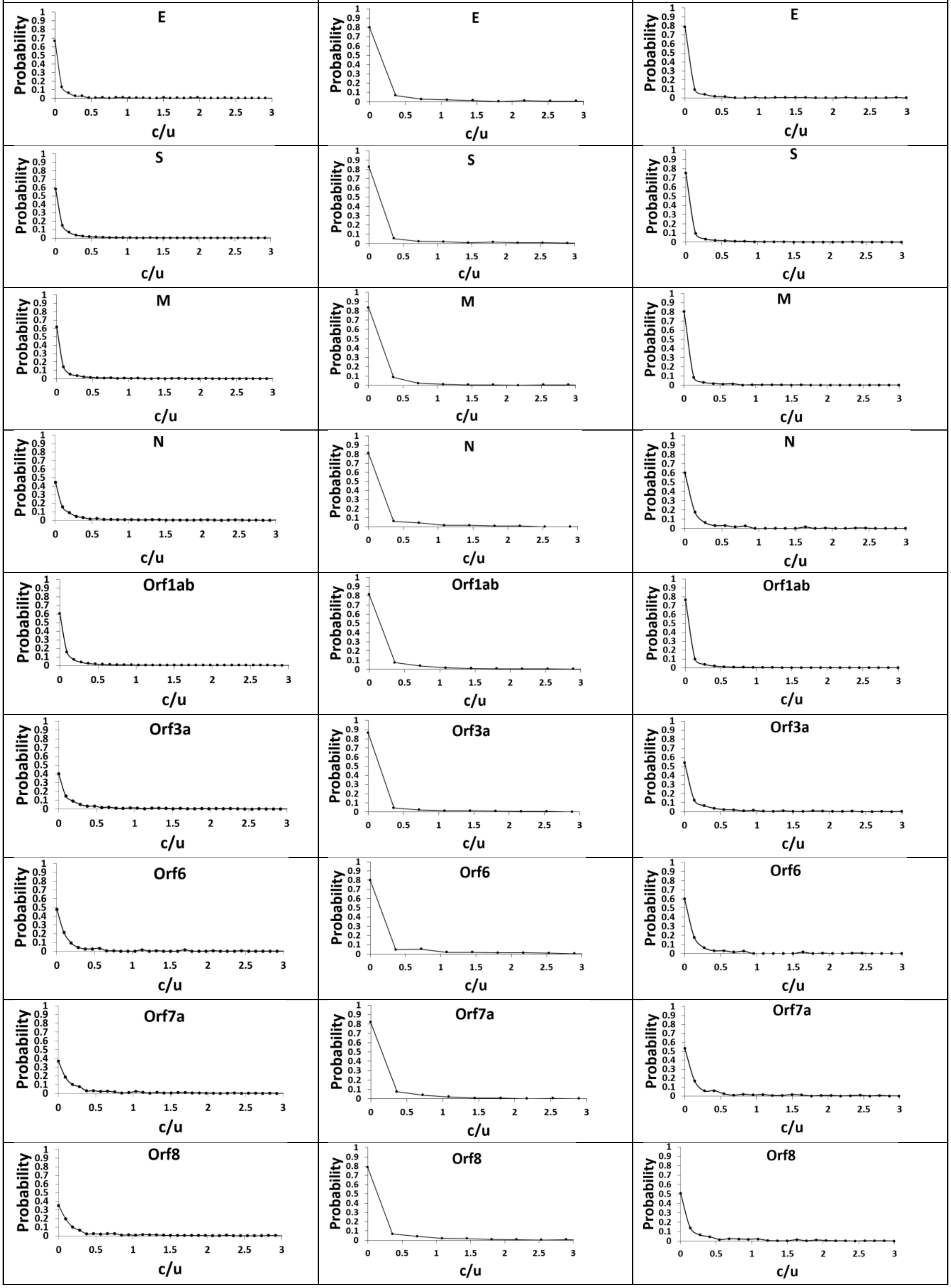

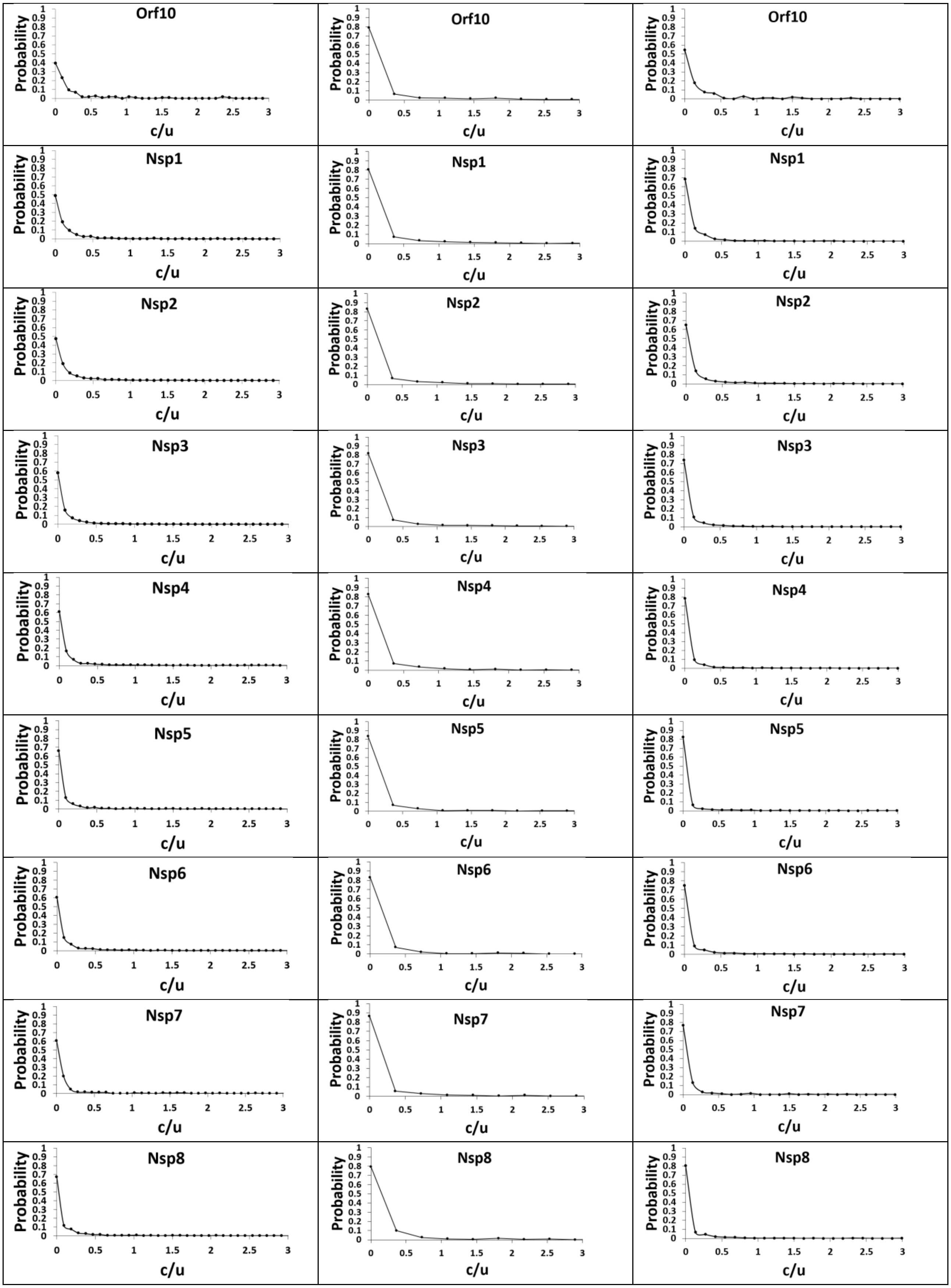

Orf10
Orf10
Orf10
Nsp1
Nsp1
Nsp1
Nsp2
Nsp2
Nsp2
Nsp3
Nsp3
Nsp3
Nsp4
Nsp4
Nsp4
Nsp5
Nsp5
Nsp5
Nsp6
Nsp6
Nsp6
Nsp7
Nsp7
Nsp7
Nsp8
Nsp8
Nsp8
Probability
c/u

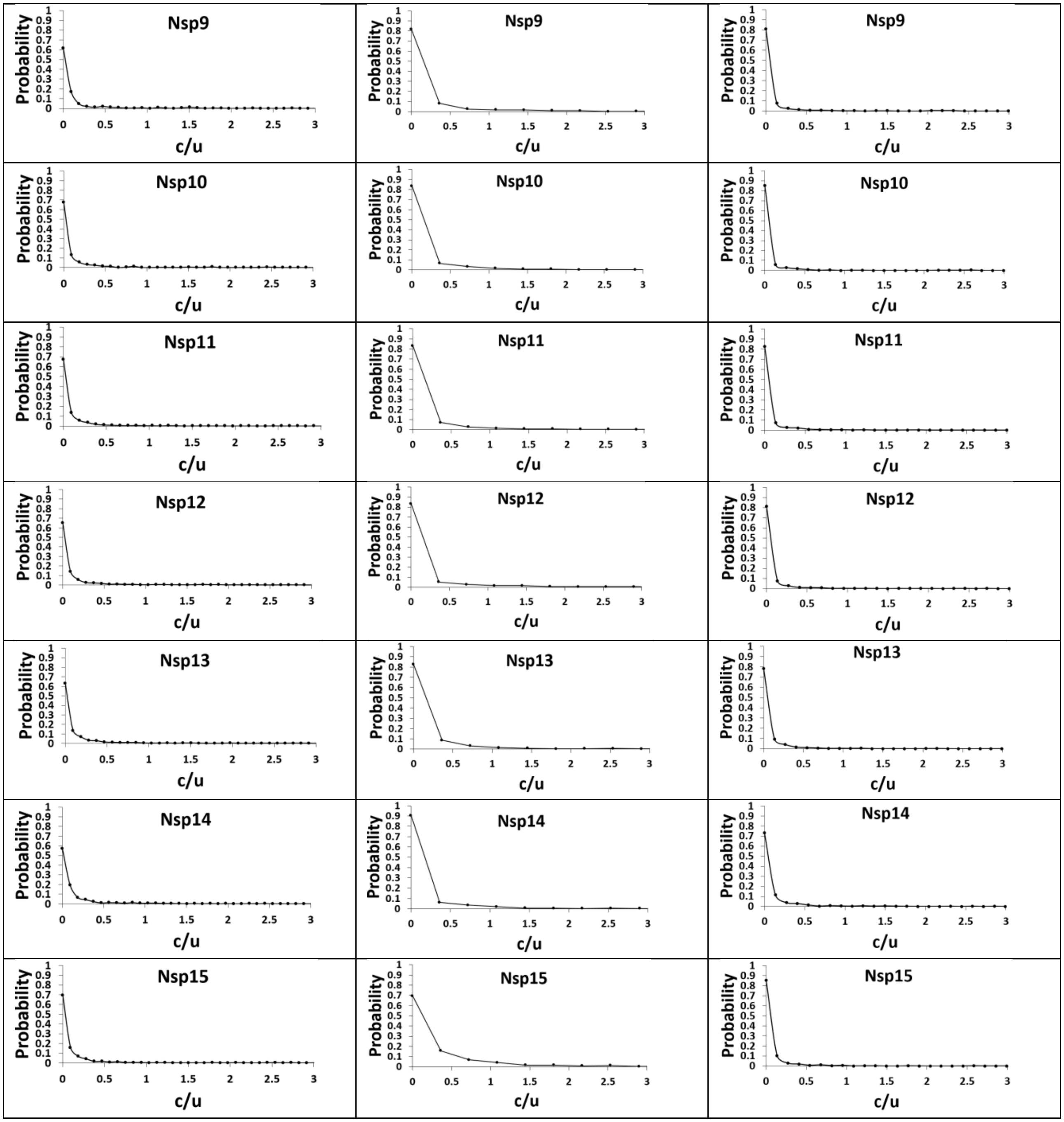


**Figure S7.** Probability distribution of the relative total NT substitution rate, relative Synonymous NT substitution rate and relative Non-Synonymous NT substitution rate (*c/μ*) for the major and accessory coding genes and Nsp1-15 across datasets A1a/A1b/A1c.

c = number of substitutions
μ = average number of substitutions per site per 19 months: **Total NT**: 10.6; **Synonymous NT:** 2.8; **Non-Synonymous NT**: 7.4

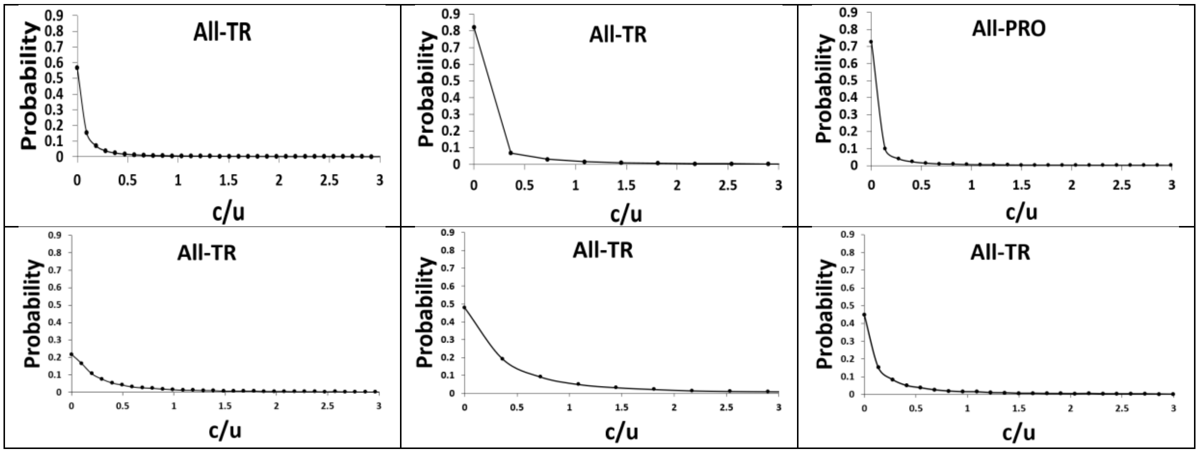


**Figure S8.** Observed L-shaped probability distribution of relative substitution rates (*c/μ*) for All-TR in the genome at the NT level (top row) and codon level (bottom row).

c = number of substitutions

μ = average number of substitutions per site per 19 months: **Total NT**: 10.6; **Synonymous NT:** 2.8; **Non-Synonymous NT**: 7.4

| | |
|---|---|
| Orf1ab 5’UTR | 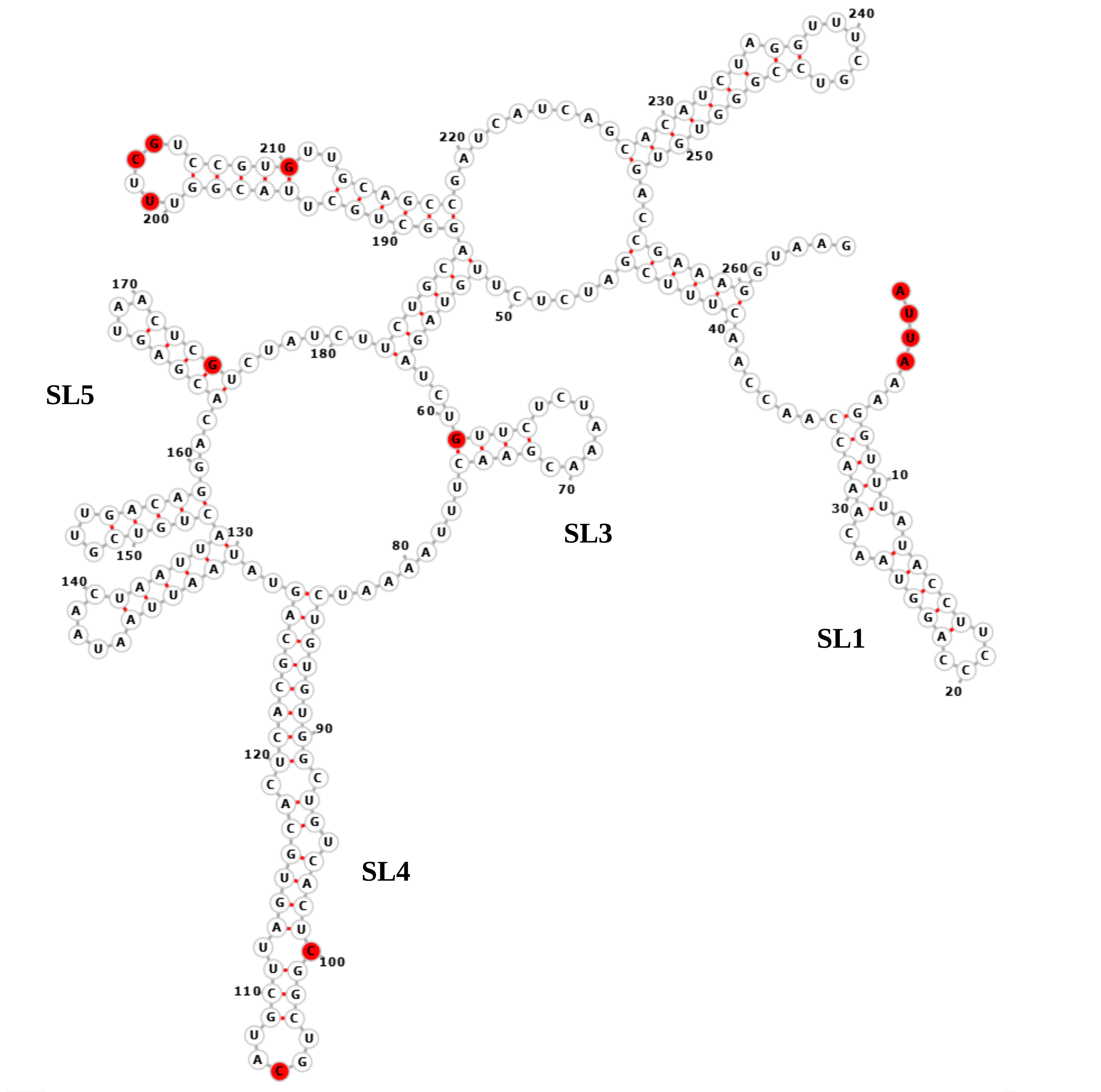  |
| E 5’UTR | 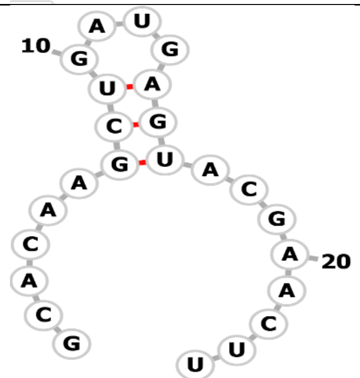 |

| | |
|---|---|
| M<br>5'UTR | 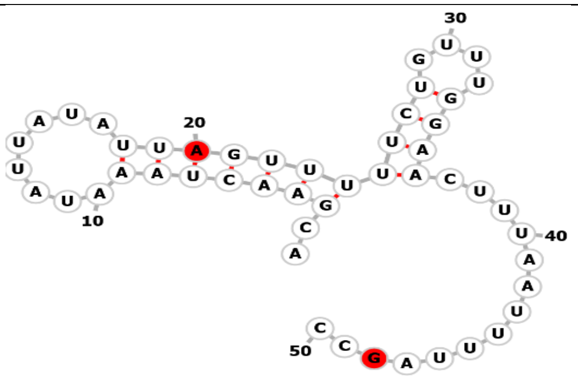  |
| N<br>5'UTR | 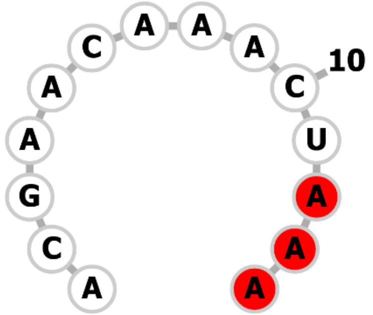  |
| Orf3a<br>5'UTR | 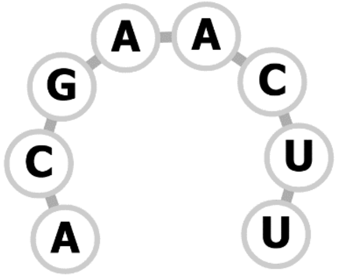 |
| Orf6<br>5'UTR | 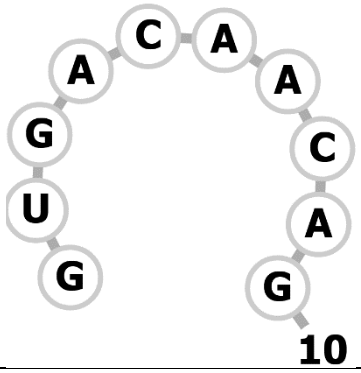  |
| Orf7a<br>5'UTR | 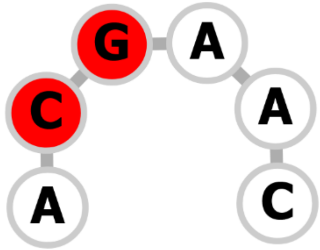 |

| | |
|---|---|
| Orf8 5’UTR | 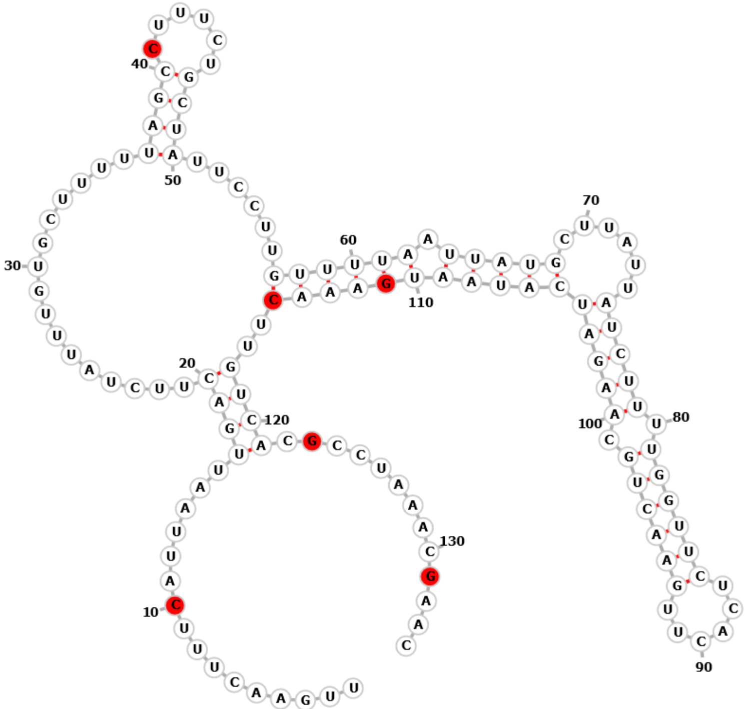  |
| Orf10 5’UTR | 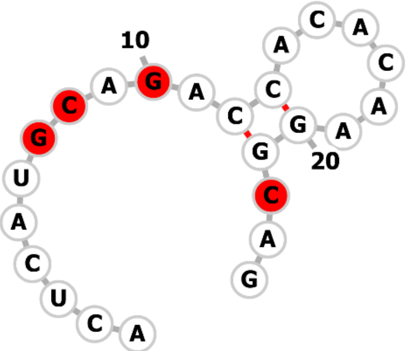  |

| | |
|---|---|
| Orf10<br>3’UTR | 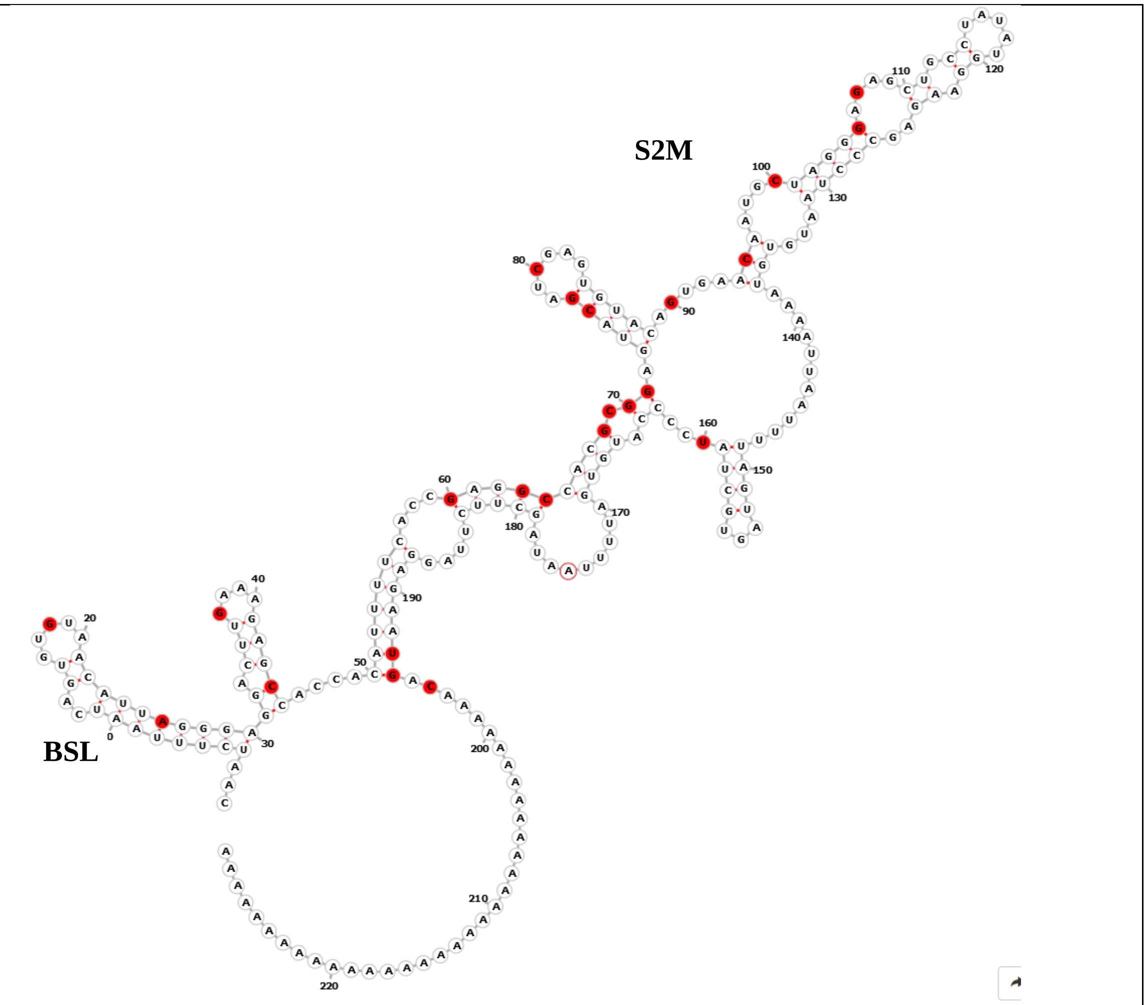 |

**Figure S9.** Thermodynamic ensemble prediction of the secondary structure of each UTR using RNAfold, using the default settings. The Wuhan-Hu-1 reference sequence was used as input for generating each secondary structure. The top 54 NTs which exhibited strong positive selection ($c$>30 and $c/\mu$>3.0) are colored in red.

*For Orf1ab 5’UTR, SL2 was not predicted by RNAfold; part of SL5 and the whole of SL6 and SL7 are not shown as we only show the first 250 NTs.

**SL: Stem-loop; BSL: Bulged stem loop; S2M: Stem-loop 2 motif

| | | |
|---|---|---|
| 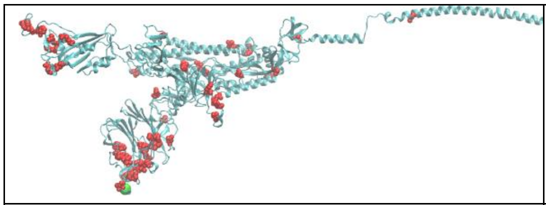 | 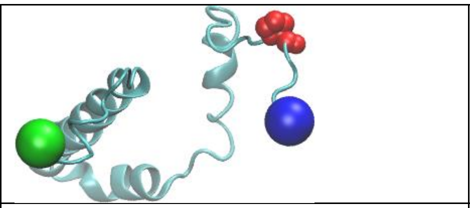 | 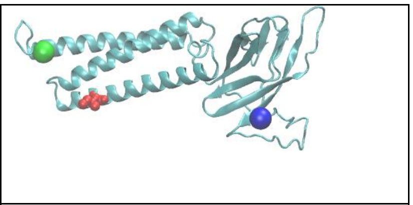 |
| **A** (Spike protein, 6vyb) | **B** (Envelope protein, 5x29) | **C** (Membrane protein, Zhanglab) |
| 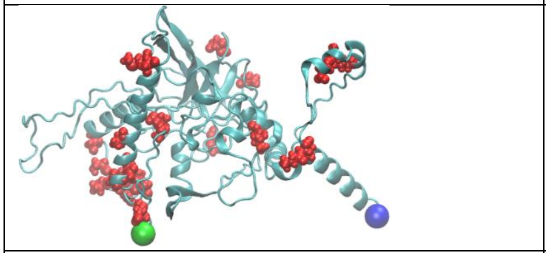 | 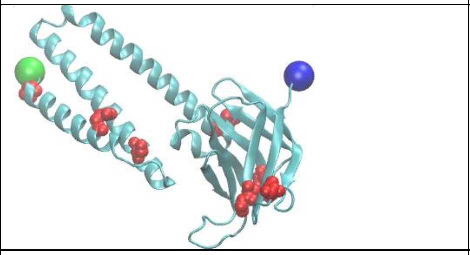 | 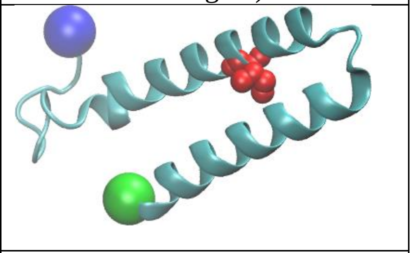 |
| **D** (Nucleocapsid protein, 6yi3 & 6yun) | **E** (Orf3a, 7kjr) | **F** (Orf6, Zhanglab) |
| 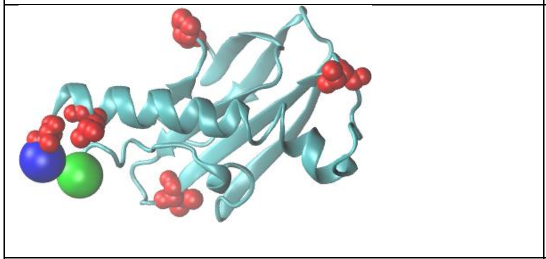 | 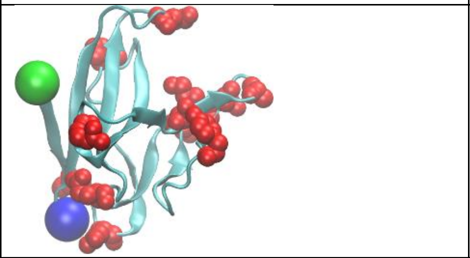 | 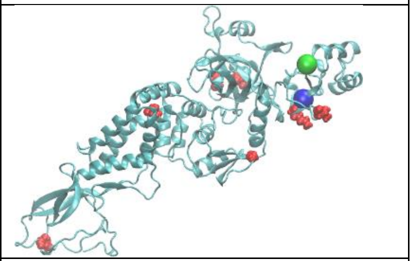 |
| **G** (Orf7a, 7ci3) | **H** (Orf8, 7jx6) | **I** (NSP2, 7msw) |
| 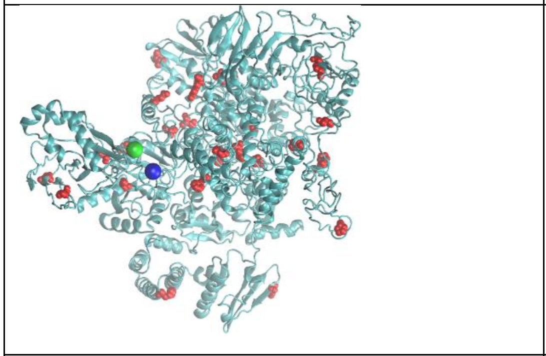 | 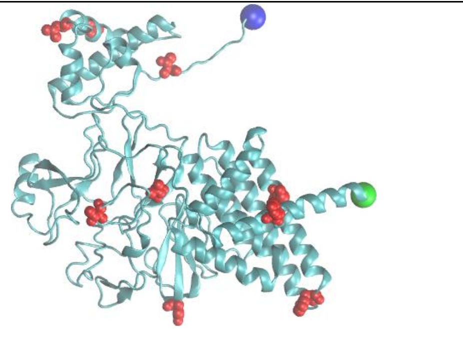 | 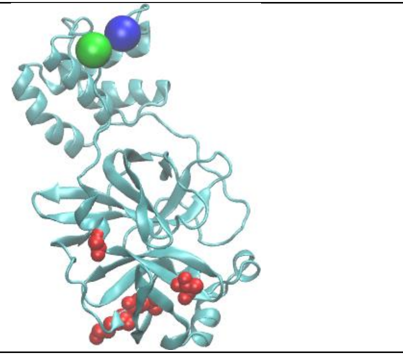 |
| **J** (NSP3, Zhanglab) | **K** (NSP4, Zhanglab) | **L** (NSP5, 7cb7) |
| 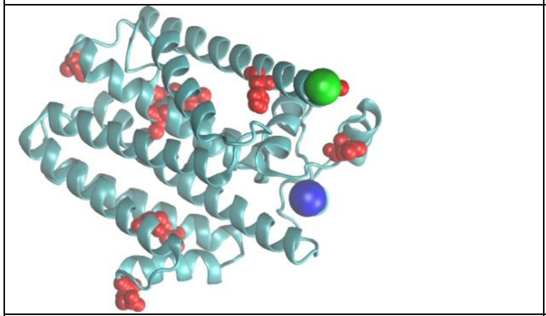 | 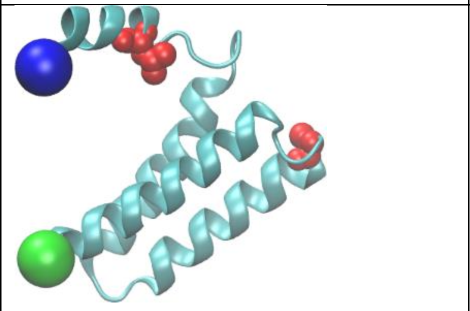 | 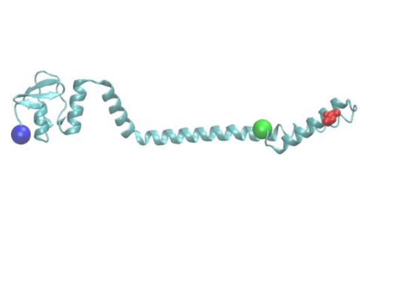 |
| **M** (NSP6, Zhanglab) | **N** (NSP7, 6m5i) | **O** (NSP8, 7cyq) |

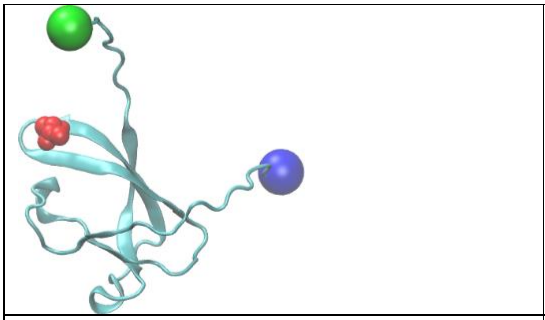

**P** (NSP9, 7kri)

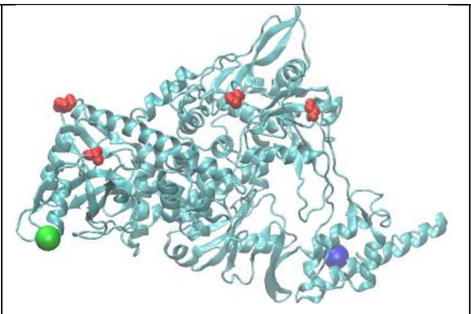

**Q** (NSP11, 7dte)

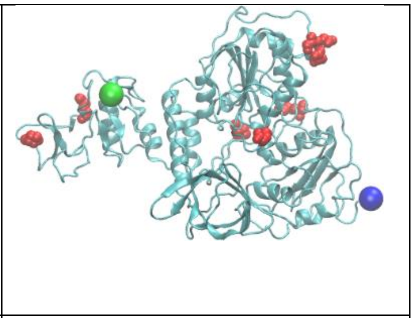

**R** (NSP12, 6xez)

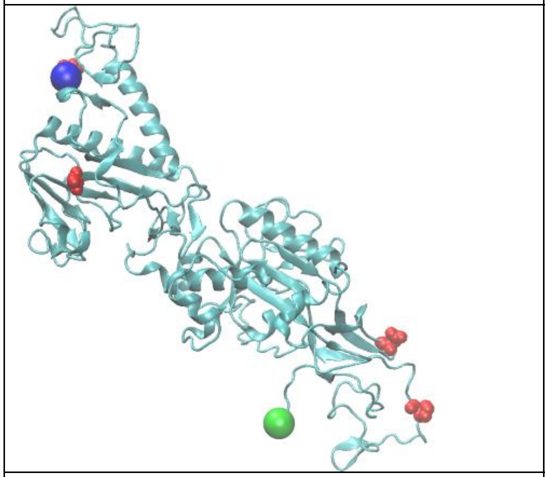

**S** (NSP13, 7egq)

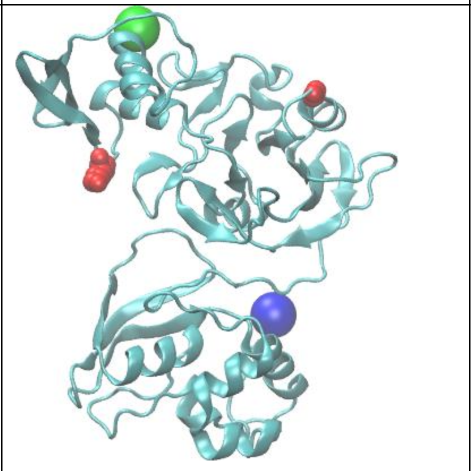

**T** (NSP14, 6wxc)

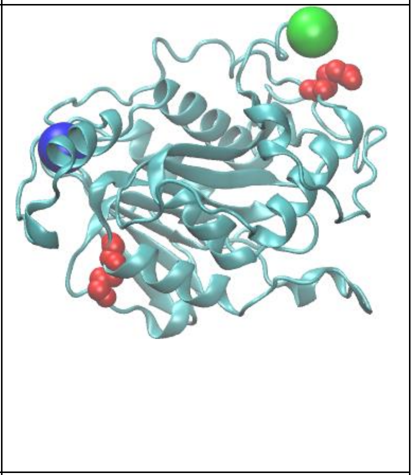

**U** (NSP15, 7I6r)

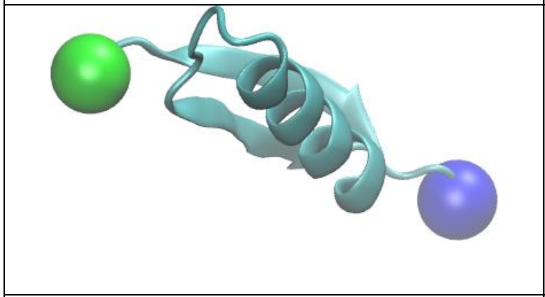

**V** (Orf10, Zhanglab)

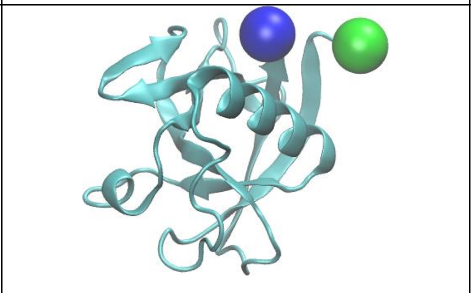

**W** (NSP1, 7k7p)

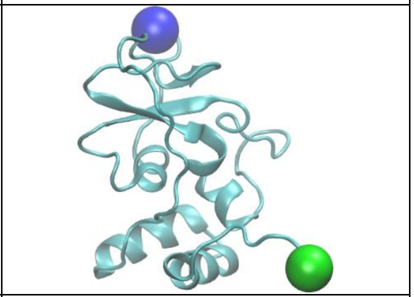

**X** (NSP10, 7n0d)

**Figure S10.** Top 247 AA sites (red spheres) in the proteome under positive selection. Protein structure models were generated from the PDB databank (with PDB ID: XXXX) or the Zhanglab homology models. Protein backbone is represented as blue ribbons. N- and C-termini are represented as green and blue VDW spheres, respectively.

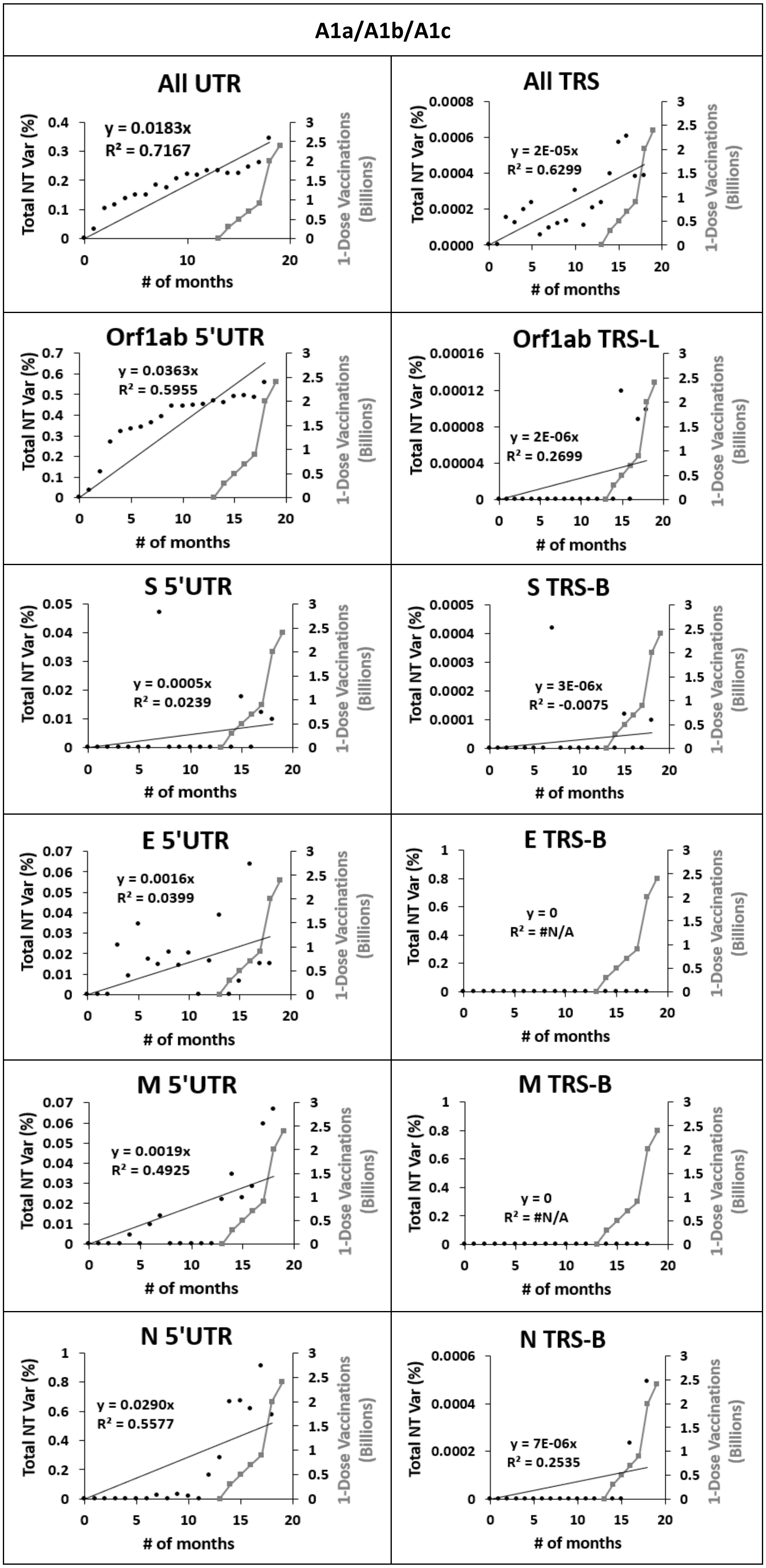
A1a/A1b/A1c
All UTR
y = 0.0183x
R² = 0.7167
All TRS
y = 2E-05x
R² = 0.6299
Orf1ab 5'UTR
y = 0.0363x
R² = 0.5955
Orf1ab TRS-L
y = 2E-06x
R² = 0.2699
S 5'UTR
y = 0.0005x
R² = 0.0239
S TRS-B
y = 3E-06x
R² = -0.0075
E 5'UTR
y = 0.0016x
R² = 0.0399
E TRS-B
y = 0
R² = #N/A
M 5'UTR
y = 0.0019x
R² = 0.4925
M TRS-B
y = 0
R² = #N/A
N 5'UTR
y = 0.0290x
R² = 0.5577
N TRS-B
y = 7E-06x
R² = 0.2535
Total NT Var (%)
1-Dose Vaccinations (Billions)
### of months

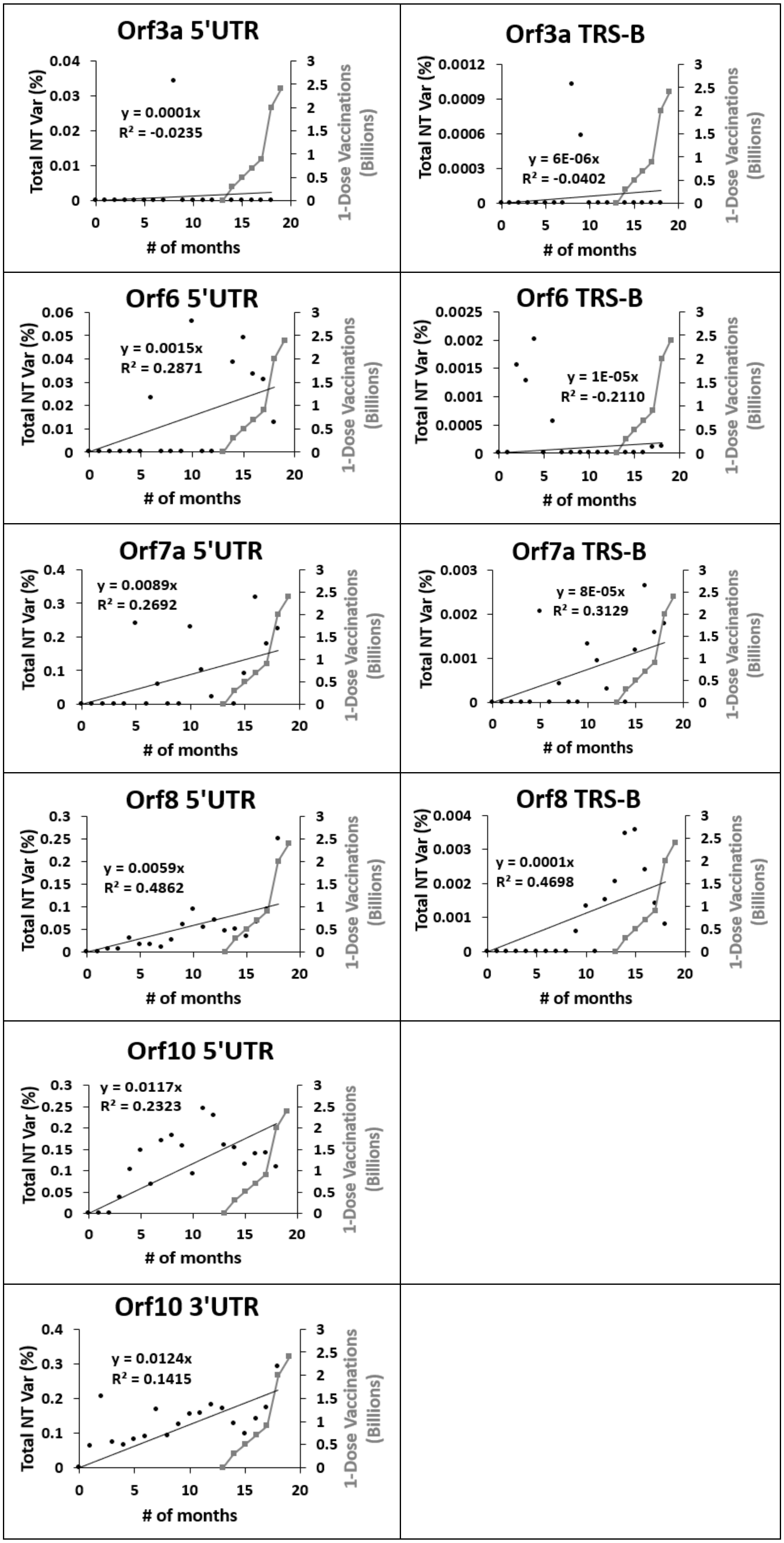

Orf3a 5'UTR
y = 0.0001x
R² = -0.0235
Orf3a TRS-B
y = 6E-06x
R² = -0.0402
Orf6 5'UTR
y = 0.0015x
R² = 0.2871
Orf6 TRS-B
y = 1E-05x
R² = -0.2110
Orf7a 5'UTR
y = 0.0089x
R² = 0.2692
Orf7a TRS-B
y = 8E-05x
R² = 0.3129
Orf8 5'UTR
y = 0.0059x
R² = 0.4862
Orf8 TRS-B
y = 0.0001x
R² = 0.4698
Orf10 5'UTR
y = 0.0117x
R² = 0.2323
Orf10 3'UTR
y = 0.0124x
R² = 0.1415
Total NT Var (%)
1-Dose Vaccinations (Billions)
### of months

**Figure S11.** Correlation between percent total NT variation timelines for each month over 19 months for UTR (left) and TRS (right) from the combined sets (A1a/A1b/A1c) and 1 dose vaccinations distributed to patients globally (grey closed squares).

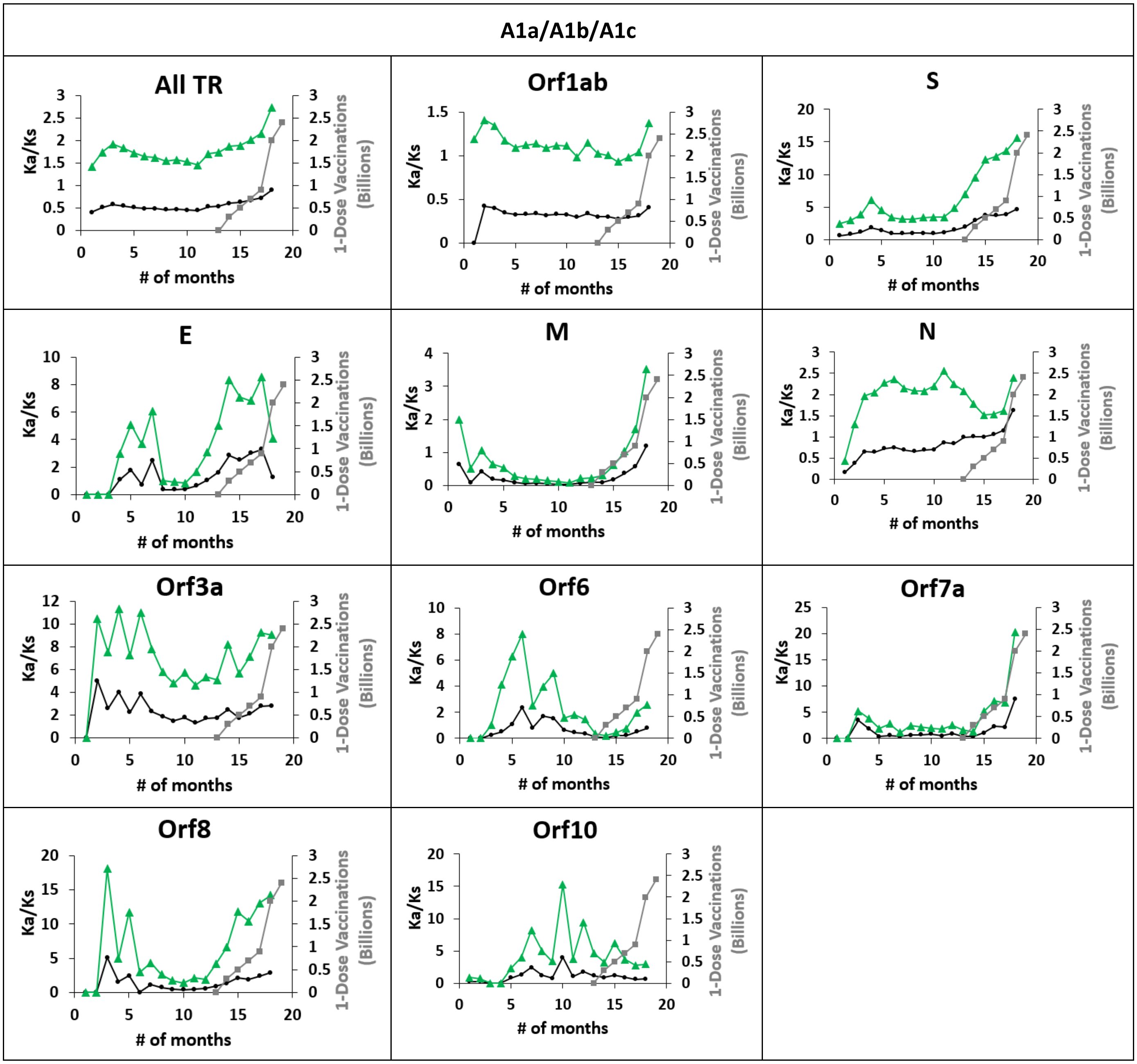


**Figure S12.** Ka/Ks timelines for each month over 19 months for All-TR, the major and accessory genes from the combined sets (A1a/A1b/A1c) using two methods: The Ka/Ks ratio using NG method (black closed circle); the Ka/Ks ratio by dividing non-synonymous NT substitution rate by synonymous NT substitution rate (not shown) derived from SP approximation (green closed triangle), and 1-dose vaccinations distributed to patients globally (grey closed squares).

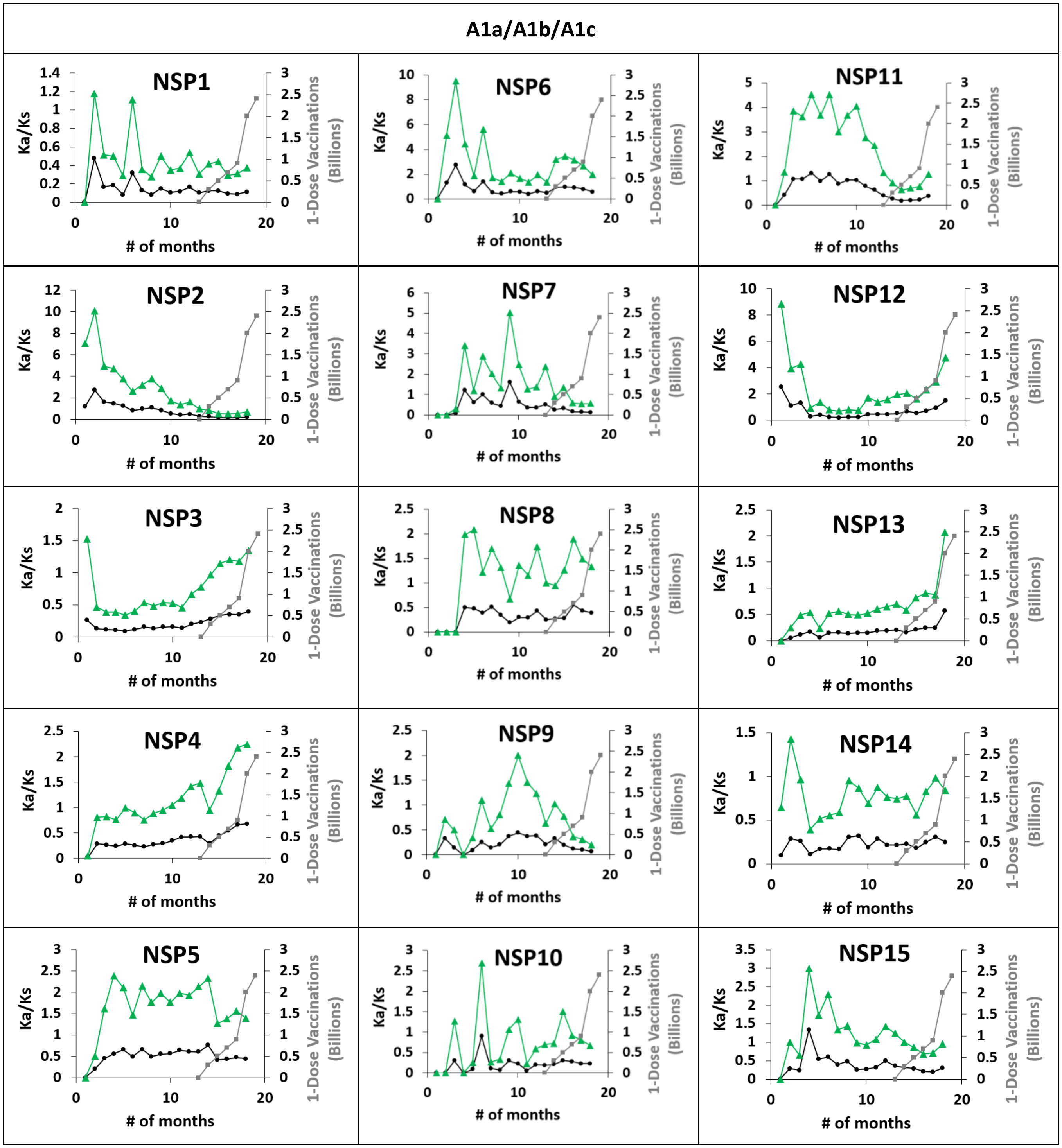


**Figure S13.** Ka/Ks timelines for each month over 19 months for Nsp1-15 from the combined sets (A1a/A1b/A1c) using two methods: The Ka/Ks ratio using NG method (black closed circle); the Ka/Ks ratio by dividing non-synonymous NT substitution rate by synonymous NT substitution rate (not shown) derived from SP approximation (green closed triangle), and 1-dose vaccinations distributed to patients globally (grey closed squares).

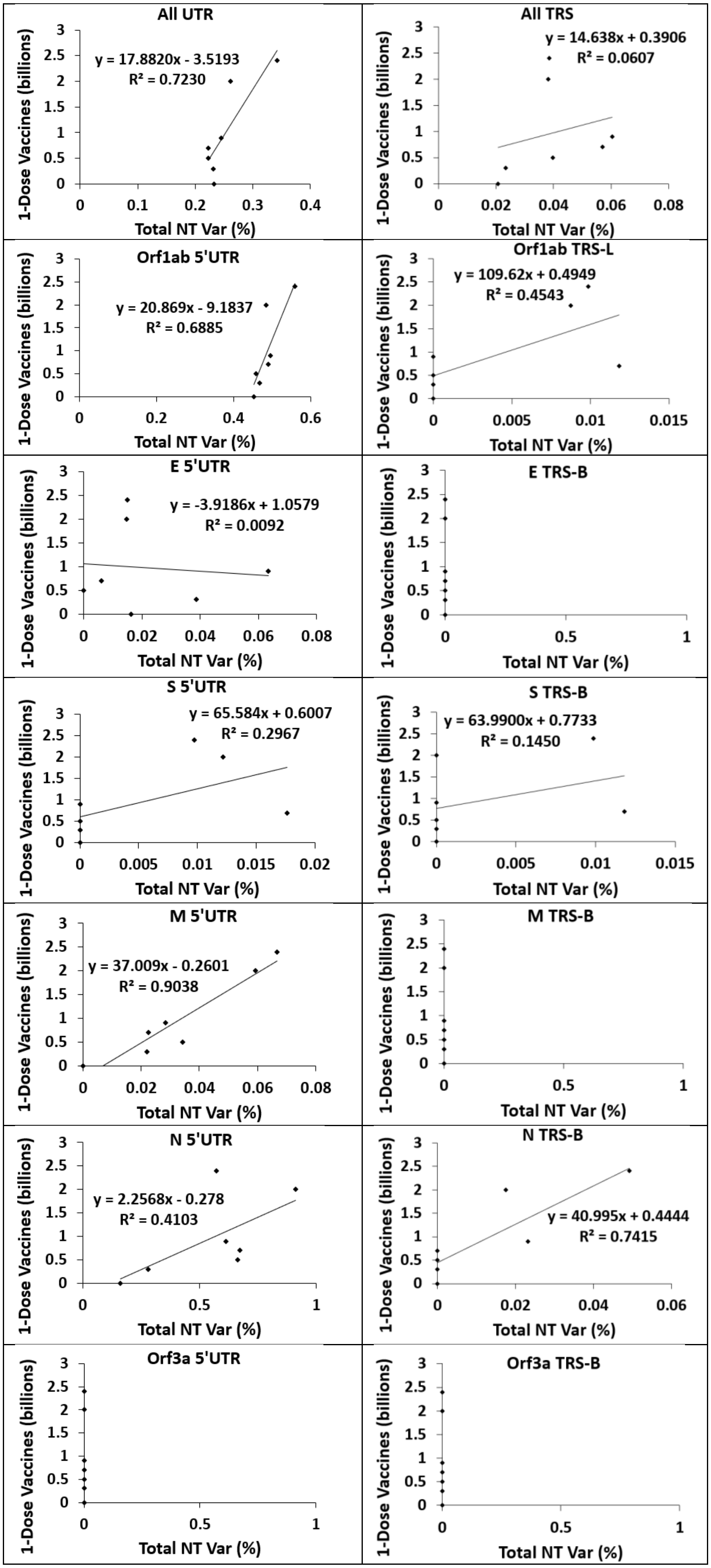
All UTR
y = 17.8820x - 3.5193
R² = 0.7230
All TRS
y = 14.638x + 0.3906
R² = 0.0607
Orf1ab 5'UTR
y = 20.869x - 9.1837
R² = 0.6885
Orf1ab TRS-L
y = 109.62x + 0.4949
R² = 0.4543
E 5'UTR
y = -3.9186x + 1.0579
R² = 0.0092
E TRS-B
S 5'UTR
y = 65.584x + 0.6007
R² = 0.2967
S TRS-B
y = 63.9900x + 0.7733
R² = 0.1450
M 5'UTR
y = 37.009x - 0.2601
R² = 0.9038
M TRS-B
N 5'UTR
y = 2.2568x - 0.278
R² = 0.4103
N TRS-B
y = 40.995x + 0.4444
R² = 0.7415
Orf3a 5'UTR
Orf3a TRS-B
1-Dose Vaccines (billions)
Total NT Var (%)

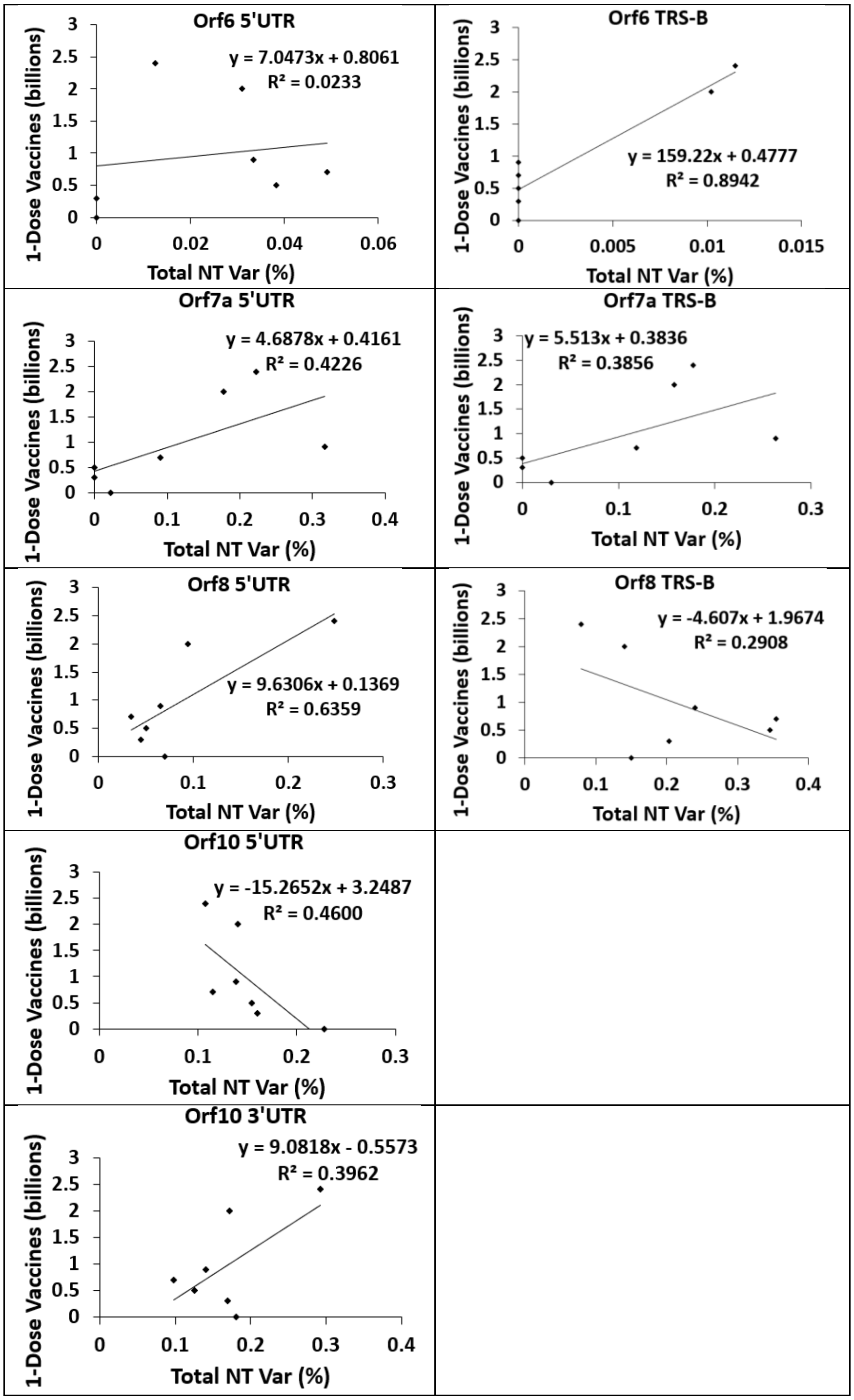


**Figure S14.** Correlation between 1-dose vaccination timeline and the percent total NT variation timeline for All-UTR, All-TRS, each UTR and TRS between December 2020-June 2021.

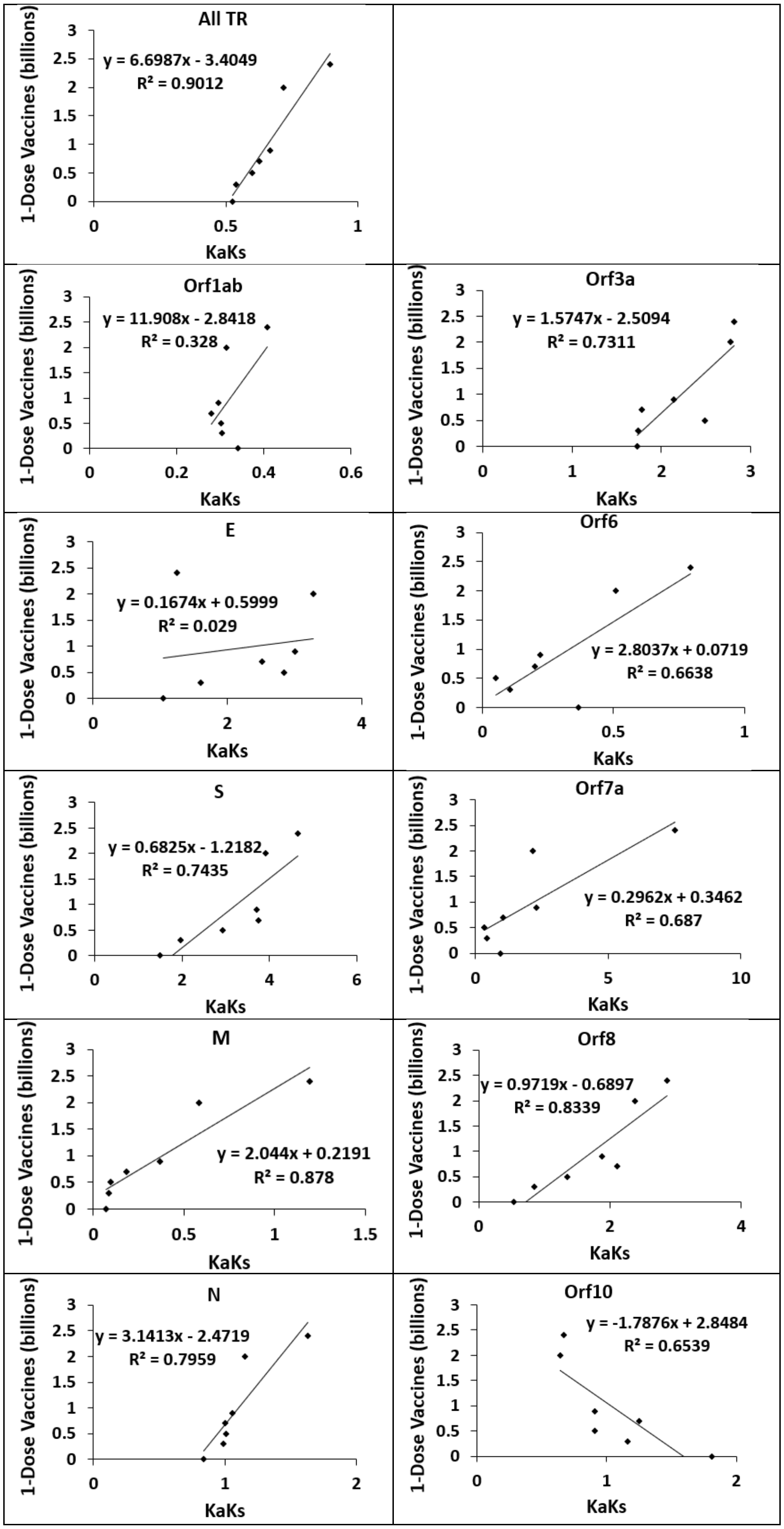
All TR
y = 6.6987x - 3.4049
R² = 0.9012
1-Dose Vaccines (billions)
0
0.5
1
1.5
2
2.5
3
0
0.5
1
KaKs
Orf1ab
y = 11.908x - 2.8418
R² = 0.328
0
0.5
1
1.5
2
2.5
3
0
0.2
0.4
0.6
1-Dose Vaccines (billions)
KaKs
Orf3a
y = 1.5747x - 2.5094
R² = 0.7311
0
0.5
1
1.5
2
2.5
3
0
1
2
3
1-Dose Vaccines (billions)
KaKs
E
y = 0.1674x + 0.5999
R² = 0.029
0
0.5
1
1.5
2
2.5
3
0
2
4
1-Dose Vaccines (billions)
KaKs
Orf6
y = 2.8037x + 0.0719
R² = 0.6638
0
0.5
1
1.5
2
2.5
3
0
0.5
1
1-Dose Vaccines (billions)
KaKs
S
y = 0.6825x - 1.2182
R² = 0.7435
0
0.5
1
1.5
2
2.5
3
0
2
4
6
1-Dose Vaccines (billions)
KaKs
Orf7a
y = 0.2962x + 0.3462
R² = 0.687
0
0.5
1
1.5
2
2.5
3
0
5
10
1-Dose Vaccines (billions)
KaKs
M
y = 2.044x + 0.2191
R² = 0.878
0
0.5
1
1.5
2
2.5
3
0
0.5
1
1.5
1-Dose Vaccines (billions)
KaKs
Orf8
y = 0.9719x - 0.6897
R² = 0.8339
0
0.5
1
1.5
2
2.5
3
0
2
4
1-Dose Vaccines (billions)
KaKs
N
y = 3.1413x - 2.4719
R² = 0.7959
0
0.5
1
1.5
2
2.5
3
0
1
2
1-Dose Vaccines (billions)
KaKs
Orf10
y = -1.7876x + 2.8484
R² = 0.6539
0
0.5
1
1.5
2
2.5
3
0
1
2
1-Dose Vaccines (billions)
KaKs

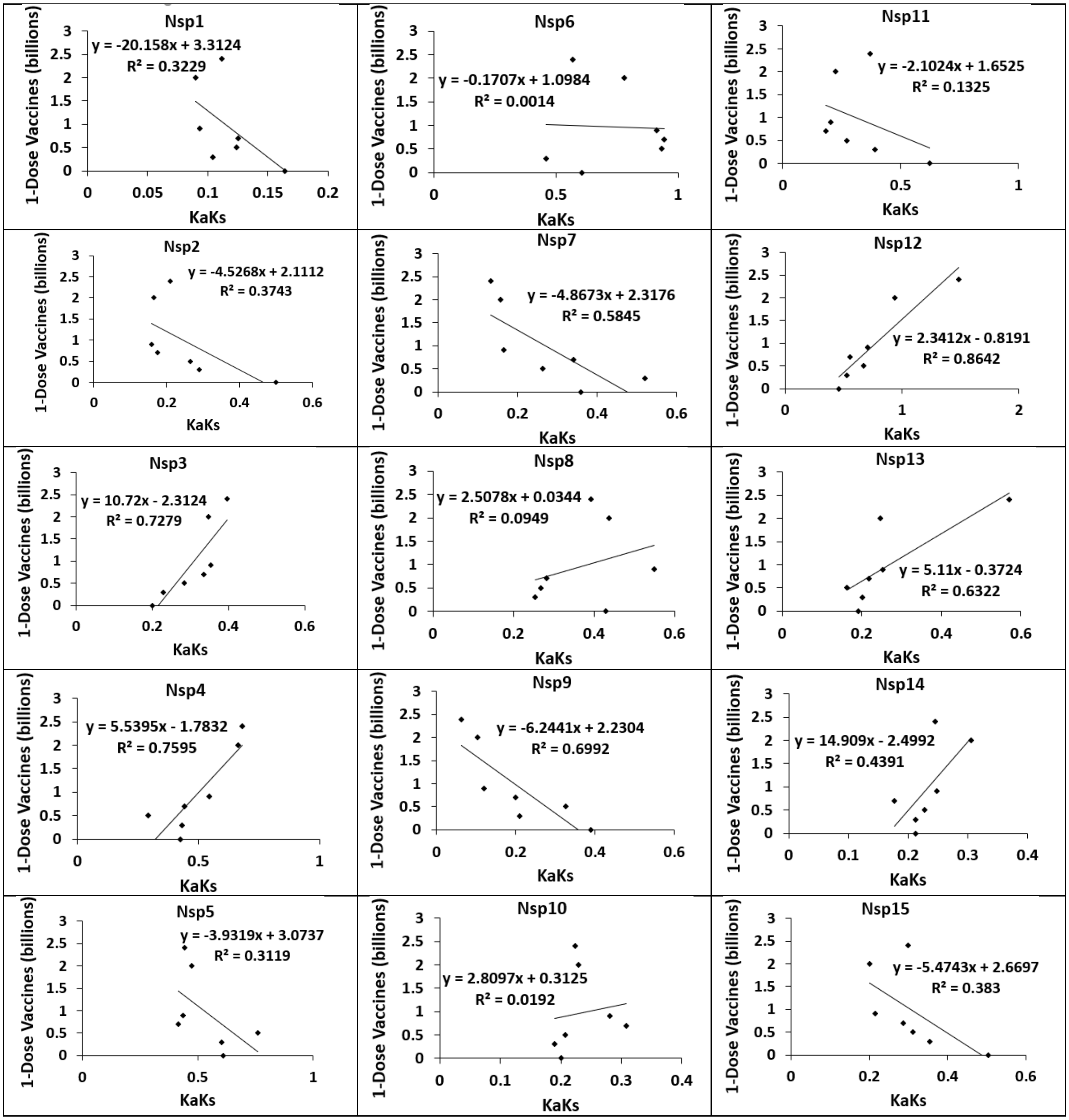


**Figure S15.** Correlation between 1-dose vaccination timeline and the NG Ka/Ks timeline for All-TR and each coding gene between December 2020-June 2021.
